# FALCON: Closed-Loop Multi-Objective Optimization of Lipid Nanoparticles for Cell-Selective mRNA Delivery

**DOI:** 10.64898/2026.06.13.731774

**Authors:** Wu Han Toh, Leonardo Cheng, Brandon Chang, Di Yu, Jingyao Ma, Xiuchun Huang, Gene Weng, Yining Zhu, Xiaoya Lu, Jinghan Lin, Jin Liu, Joseph Choy, Autumn Greco, Manav Jain, Joanna Yang, Milan Patel, Grace Shoemaker, Isabella Cozzone, Daniel Antov, Kevin Zhang, Sarp Kayabas, Charles Shin, Ataes Aggarwal, Jordan Green, Stephany Tzeng, Ramya Kumar, Maximilian F. Konig, Hai-Quan Mao

**Affiliations:** Institute for NanoBioTechnology, Johns Hopkins University, Baltimore, MD, USA; Department of Computer Science, Johns Hopkins University, Baltimore, MD, USA; Department of Biology, Johns Hopkins University, Baltimore, MD, USA; Translational Tissue Engineering Center, Johns Hopkins University School of Medicine, Baltimore, MD, USA; Department of Biomedical Engineering, Johns Hopkins University School of Medicine, Baltimore, MD, USA; Department of Materials Science and Engineering, Johns Hopkins University, Baltimore, MD, USA; Department of Biophysics, Johns Hopkins University, Baltimore, MD, USA; Department of Pathology, Johns Hopkins University, Baltimore, MD, USA; Department of Chemical and Biomolecular Engineering, Johns Hopkins University, Baltimore, MD, USA; Department of Chemical and Biological Engineering, Colorado School of Mines, Golden, Colorado, USA; Division of Rheumatology, Department of Medicine, Johns Hopkins University School of Medicine, Baltimore, MD, USA; Center for Autoimmunity and Immuno-Oncology, Johns Hopkins University School of Medicine, Baltimore, MD, USA

**Author notes:** These authors contributed equally to this work (Wu Han Toh, Leonardo Cheng).

## Abstract

Efficient, cell type-selective delivery of genetic payloads remains a central challenge in the development of gene and cell therapies. Lipid nanoparticles (LNPs) offer a versatile delivery platform, but their optimization is hindered by reliance on brute-force screening methods that are laborious, resource-intensive, and focus on single targets. Here, we present FALCON (Framework for Active Learning-driven Compositional Optimization of Nanoparticles), a closed-loop pipeline that leverages iterative screening, surrogate modeling, and multi-objective optimization to accelerate LNP compositional design. In B cell-targeted validation experiments, FALCON-optimized LNPs achieved a 1.8-fold increase in splenic B cell transfection *in vivo* compared with reference compositions. When optimized for selectivity, FALCON LNPs displayed an 84-fold improvement in selective transfection of splenic B cells over off-target liver populations and enabled spleen-tropic behavior across factorial panels of varying ionizable and helper lipid chemistries. In vaccine studies, these LNPs induced higher IgG2c antibody titers and a more Th1-biased immune profile. FALCON was also deployed to optimize LNPs for myeloid cell-selective delivery, achieving enhanced *in vivo* selectivity following systemic administration both across and within spleen and liver compartments. Our results establish FALCON as a useful tool for data-driven design of LNP compositions for precision gene delivery.

## INTRODUCTION

Achieving efficient, targeted delivery of genetic payloads remains a central focus for advancing gene and cell therapies. In recent years, lipid nanoparticles (LNPs) have emerged as leading non-viral carriers for mRNA delivery, providing a versatile platform for applications including mRNA vaccines, gene editing, cancer immunotherapy, and treatment for genetic disorders^1,2^. The utility of LNP-mediated delivery systems was demonstrated through the FDA approval of Onpattro, an siRNA-LNP for hereditary amyloidosis, as well as during the COVID-19 pandemic, with large-scale deployment of two LNP-based mRNA vaccines, Spikevax (Moderna) and Comirnaty (BioNTech/Pfizer), against SARS-CoV2^3–5^.

Despite their remarkable success, engineering LNPs poses an ongoing challenge due to the need for precise formulation optimization. Current FDA-approved LNP formulations consist of 4 components: ionizable cationic lipids, helper lipids, cholesterol, and polyethylene glycol (PEG)-modified lipids^6^. Numerous studies have shown that even subtle variations in component structure (i.e., lipid chemistry) and relative ratios (referred to hereon as composition) of LNPs can substantially alter their transfection efficiency and tissue-specific delivery^7–12^. Moreover, there is no universal LNP formulation; compositions must be carefully tailored for each therapeutic application^13,14^. Routine approaches for optimizing LNPs involve *in vitro* high-throughput screening (HTS) cellular assays, barcoded/cluster-mode screening *in vivo*, or a combination thereof^15–19^. Although these methods have been employed with some levels of success, they face several major obstacles.

First, the design space of LNP formulations is vast and poorly characterized, with limited insights to inform rational design. Conventional screening is performed through grid-search libraries where composition ratios are determined by empirical selections and factorial designs. However, grid-search methods frequently lack the resolution to capture critical interactions between formulation parameters, missing critical gaps in the design space, resulting in suboptimal LNP formulations, particularly for hard-to-transfect stem cells and immune cell populations. Such brute-force experimentation is not only labor-and resource-intensive but also tests only a small fraction of possible formulation permutations, yielding a handful of viable LNP formulations^20^. To overcome the low hit rate and inefficient experimental design of conventional combinatorial screening, more adaptive data-driven screening approaches are needed^14^.

Second, off-target delivery arises from a lack of consideration of biological contexts during formulation design, raising substantial downstream safety and efficacy concerns. Side effects caused by ectopic transgene expression remain one of the biggest challenges and needs for gene therapy^21–23^. Current screening methods, however, are limited to single-cell target optimizations that fail to account for the complexity of tissue microenvironments, cellular neighborhoods, and barriers associated with different administration routes. For instance, intravenously administered LNPs must navigate the bloodstream while avoiding rapid clearance by phagocytic immune cells^24^. Additionally, nonspecific uptake in off-target tissues, particularly the liver, spleen, and lungs, may generate unwanted side effects and should ideally be minimized^14^. Such factors are not captured in current *in vitro* screening models. Our recent study, along with others, has shown that the LNP transfection profiles of different cell types and subtypes are governed by distinct composition design rules^13,25^. As such, systematically enhancing LNP targeting precision and minimizing off-target effects will require approaches capable of multi-dimensional optimization across diverse cell types.

Machine learning (ML) offers a powerful tool for tackling these challenges^20,26^. By effectively modeling complex multifactorial relationships, ML has been successfully employed to predict the functional performance of various lipid-and polymer-based carriers using their physicochemical, structural, and compositional attributes^13,27–30^. In this work, we show that leveraging ML to guide experimental screening can direct formulation search in an efficient yet exhaustive manner, making it an attractive approach for LNP optimization tasks. Conventional ML methods rely on large, well-curated datasets, which present a substantial experimental burden. This challenge can be addressed through active learning, where models trained on sparse initial datasets iteratively identify samples to test^31–33^. In this work, we define active learning in an exploitative sense. Rather than selecting the most informative samples to label from a fixed pool, our framework generates new candidate designs by optimizing a surrogate model over a continuous, expandable composition space. The primary aim is not broad uncertainty reduction, but rapid identification of high-performing formulations. Accordingly, each iteration prioritizes regions of composition space with the greatest predicted objective potential, directing experimental resources toward performance-driven discovery. When combined with multi-objective optimization algorithms capable of balancing the competing transfection profiles of multiple cell types, this approach enables rapid and thorough discovery of selective LNP compositions that maximize delivery to a target cell type while minimizing delivery to off-target cell types^34,35^.

To date, most ML-based LNP screening approaches have focused on applying various deep learning, tree-based, and large language models to optimize the structures of ionizable lipids^36–41^. In this work, we focused on optimizing the compositional ratios of LNPs. While lipid chemistry is a major determinant of tissue and cell tropism, we demonstrate that even when lipid choices are held constant, the compositional ratios of the four lipid components and RNA cargo constitute a powerful and underexplored design space. Within a fixed lipid set, systematic ratio tuning can substantially alter both potency and selectivity. Moreover, current ML pipelines for LNP composition screening lack structured optimization frameworks capable of proposing novel formulations. Researcher-directed selection of LNP formulations is inefficient and biased, particularly when considering multi-objective optimization problems, such as cell type-specific mRNA delivery. Despite growing interest in ML-assisted LNP discovery, relatively few approaches directly integrate predictive composition–function models with formal optimization algorithms to guide rational formulation search.

Here, we present FALCON (Framework for Active Learning-driven Compositional Optimization of Nanoparticles), a closed-loop experimental-computational pipeline that integrates robotic LNP assembly and screening, supervised ML models, and multi-objective search algorithms to accelerate the optimization of cell type-selective mRNA LNPs. Inspired by the rise of self-driving labs, FALCON utilizes an iterative design-build-test-learn (DBTL) framework^32,42–44^ that optimizes LNP composition across successive experimental iterations, rapidly converging on regions of the parameter space with the highest cell type-selective transfection potential. To validate FALCON, we first optimized LNPs for potent mRNA delivery to B cells. We show that in several iterative cycles, FALCON converges on optimized LNPs that considerably outperform benchmark formulations, demonstrating superior transfection efficiency in both primary human B cells *ex vivo* and in an *in vivo* mouse reporter model. We then demonstrate FALCON’s ability to conduct multi-objective optimization by designing LNPs that maximize B-cell transfection while minimizing hepatocyte delivery. We show that this optimized composition achieves enhanced organ-level spleen-to-liver and *in vivo* single-cell B cell-selective transfection compared to benchmark LNPs and, notably, enables spleen-selective delivery across several ionizable and helper lipid chemistries while inducing enhanced IgG2c antibody responses in vaccine studies. Finally, we deploy FALCON in additional case studies, including optimizing LNPs for myeloid cell-selective delivery *in vivo*, to demonstrate FALCON’s utility in different cell targets. Through SHAP feature importance analysis, we investigate compositional design rules associated with cell-selective delivery. Together, our results establish FALCON as an effective and versatile platform for intelligent compositional optimization of LNPs.

## RESULTS

### Design of FALCON

FALCON uses ML-guided search algorithms instead of brute-force trial-and-error analysis to drive intelligent LNP formulation optimization (**Fig. 1**). Under this framework, researchers initiate and monitor FALCON while it proceeds through independent DBTL optimization cycles.

**Figure 1.**
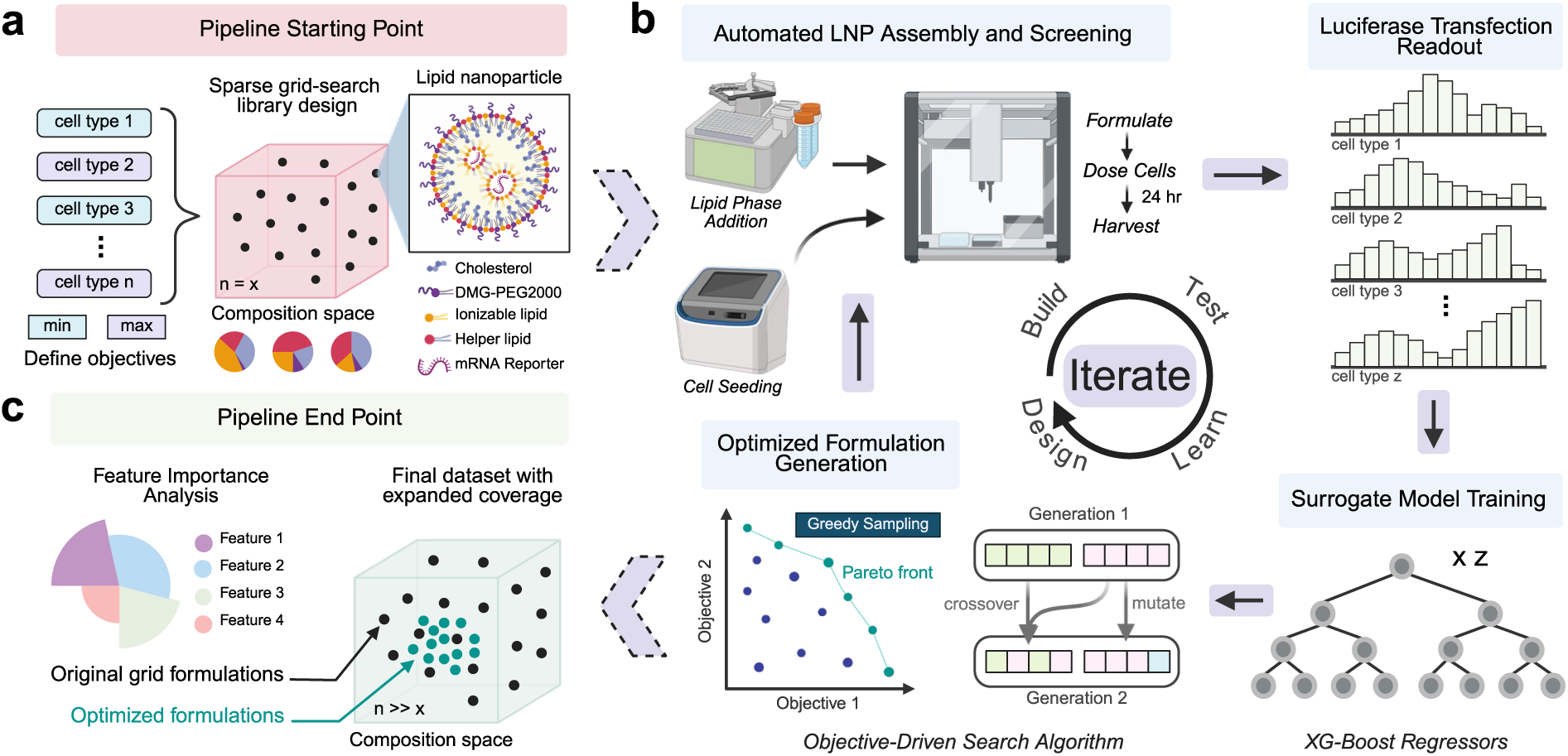
**Schematic overview of FALCON workflow**. **a.** Pipeline starting point. An initial sparse grid-search library was designed by varying parameters influencing the relative ratio of LNP lipid components and payload. Cell type objectives, *i.e.*, maximize or minimize transfection, were defined to enable cell-selective optimization. **b.** Illustration of the DTBL framework for ML-driven optimization. Each cycle was performed in 4 steps: (1) candidate LNPs were formulated and treated to cells using a robotic workstation, (2) transfection was quantified by luciferase expression; (3) the resulting composition-function dataset was used to train or refine a surrogate ML model, which then (4) guides an exhaustive search with single-or multi-objective optimization algorithms to identify LNPs for testing in the next iterative cycle. **c.** Pipeline endpoint following convergence on optimal LNP compositions. An expanded dataset was obtained and used to train a final model, which was analyzed using SHAP analysis to interpret design rules for cell-selective LNPs. This figure was created with BioRender.com and released under a Creative Commons Attribution-NonCommercial-NoDerivs 4.0 International license.

The first step of FALCON involves designing an initial training library. Compared to conventional ML approaches that rely on high-throughput screening to generate large training datasets, FALCON can be initiated from a sparse grid-search library, making it feasible to employ in routine screening (**Fig. 1a**). The initial libraries used in this study were generated by varying 4 independent composition parameters: (i) the total molar percentage of charged lipids, i.e., ionizable lipid (IL) and helper lipid (HL) (“(IL + HL)%”), (ii) helper lipid molar percentage of charged lipids (“HL/(IL + HL)%”), (iii) PEGylated lipid molar percentage of uncharged lipids, *i.e.*, cholesterol (Chol) and PEGylated lipid (PEG) (“PEG/(PEG + Chol)%”), and (iv) protonatable nitrogen to phosphate ratio (“N/P ratio”). (**Supplementary Tables 1–4**)^18^. As a preliminary validation that the datasets generated from our sparse library were sufficient to initiate FALCON, we ensured that the transfection readouts obtained spanned a broad performance range, and models trained on these datasets outperformed baseline on held-out samples while enabling stable acquisition of new samples, indicating the presence of learnable structure to guide early formulation search (**Supplementary Fig. 1**).

Following initial library design, the remaining phases of FALCON are carried out in a data-driven experimental loop consisting of four main stages: (1) candidate LNPs carrying firefly luciferase reporter mRNA are assembled and delivered to relevant cell types in parallel using a robotic workstation; (2) transfection is quantified through bulk luciferase expression in a 24-h assessment; (3) experimentally acquired composition-function data is used to train or refine an ML-based surrogate model; and (4) search algorithms that specify multiple optimization objectives suggest a new batch of LNP compositions with improved predicted outcomes for testing in the next cycle (**Fig. 1b**). Each FALCON iteration proceeds independently and can be repeated until predefined stopping conditions are met. This iterative framework is designed to progressively enrich the dataset with samples from high-performing regions of interest, allowing the surrogate model to refine its representation of the local design landscape. As additional data are incorporated, model-guided sampling can more efficiently focus experimental exploration on promising areas of the formulation space. Feature importance analysis can be performed on the final trained ML models to identify design rules for achieving cell-type-selective LNP compositions (**Fig. 1c**)^45,46^.

We deployed a robotic in-house setup to assemble and screen LNPs, increasing the speed of data generation while minimizing variation across iterations (**Supplementary Fig. 2**). The size, polydispersity, encapsulation efficiency, and transfection performance of LNPs assembled using this approach were comparable to those of LNPs manually formulated through established methods (**Supplementary Fig. 3**)^18^.

### Model training and search strategy for formulation optimization

ML models were trained to predict transfection performance as a function of the four LNP composition parameters described above. For each cell type objective, model training was performed independently. Datasets obtained from iterative screening underwent a series of data cleaning steps, including floor adjustment and min-max scaling (see “Methods”). Batch-wise normalization of transfection performance, obtained from bulk luciferase assay, was performed using a set of internal controls included in each screening cycle to correct for batch effects. Among a panel of regression models, the XGBoost model trained on our datasets achieved similar performance to kNN, RF, MLP, LGBM, and DT models and markedly better performance to MLR, PLS, and lasso (**Supplementary Fig. 4**). Given its strong predictive performance and clear interpretability for SHAP analysis, we selected XGBoost as our surrogate model for downstream optimization studies^13^. Prior to model training, data was randomly split into an 85:15 train-test ratio. A 5-fold cross-validation was performed on the training set to optimize hyperparameters, with the best-performing model selected based on the lowest Mean Absolute Error (MAE) across validation folds (see “Methods”). Following each iteration, models were retrained with updated datasets, incorporating newly acquired data to refine the model’s predictions. When evaluated on a fixed global hold-out set (15% of the final dataset, randomly sampled prior to model training and held constant across all iterations), predictive performance generally remained stable or improved across successive iterations (**Supplementary Fig. 5**).

We explored three model-guided optimization algorithms in this study: Bayesian Optimization (BO) and Dual Annealing (DA) for single-objective tasks, and Non-Dominated Sorting Genetic Algorithm II (NSGA-II) for multi-objective optimization tasks (**Supplementary Fig. 6**). BO is a surrogate model-guided strategy that uses an acquisition function to balance exploration and exploitation when proposing new candidate formulations^33^. DA is a global optimization algorithm that explores rugged, non-convex parameter spaces through controlled randomness while using the trained surrogate to evaluate candidate solutions^47^.

Multi-objective optimization of LNP composition across cell types, where transfection profiles can be contrasting or overlapping, requires algorithms capable of exploring trade-offs in LNP design. To achieve this level of control over delivery profiles, we employed NSGA-II, an evolutionary optimization method that evolves a population of Pareto-optimal solutions over multiple generations^48^. On-target and off-target transfection were optimized as independent objectives rather than being combined into a single selectivity score, thereby avoiding the optimization of ratio-based metrics that can artificially favor trivially low-transfection formulations. Following the completion of the algorithm, diversity-weighted greedy selection was performed to sample along the Pareto front, selecting a batch of LNPs for testing that spanned a wide range of trade-offs. Final candidate LNPs for validation were selected from the Pareto-optimal set by applying biologically motivated transfection thresholds and ranking the remaining formulations according to their selectivity scores (see “Methods”).

Leveraging these search algorithms, FALCON conducts a comprehensive virtual screening of thousands of LNP compositions before nominating the top-performing LNPs *in silico* to test experimentally, thus exploring the parameter space more extensively than through experimental screening alone. Representative plots of the convex hull volume encompassing the evaluated points of each algorithm are shown in **Supplementary Fig. 7**, illustrating the breadth and coverage of these evaluations across the search space.

### FALCON efficiently explores composition space to optimize LNPs for transfection of B cells

B cells play a critical role in humoral immunity and are increasingly being explored as engineered cell therapies for applications ranging from combating autoimmunity, cancer, and infectious diseases to the *in vivo* production of therapeutic proteins for inherited metabolic disorders^49,50^. However, efficient non-viral delivery systems for B cell transfection remain underdeveloped. As such, they were chosen as the first target for FALCON-driven LNP optimization (**Fig. 2**).

**Figure 2.**
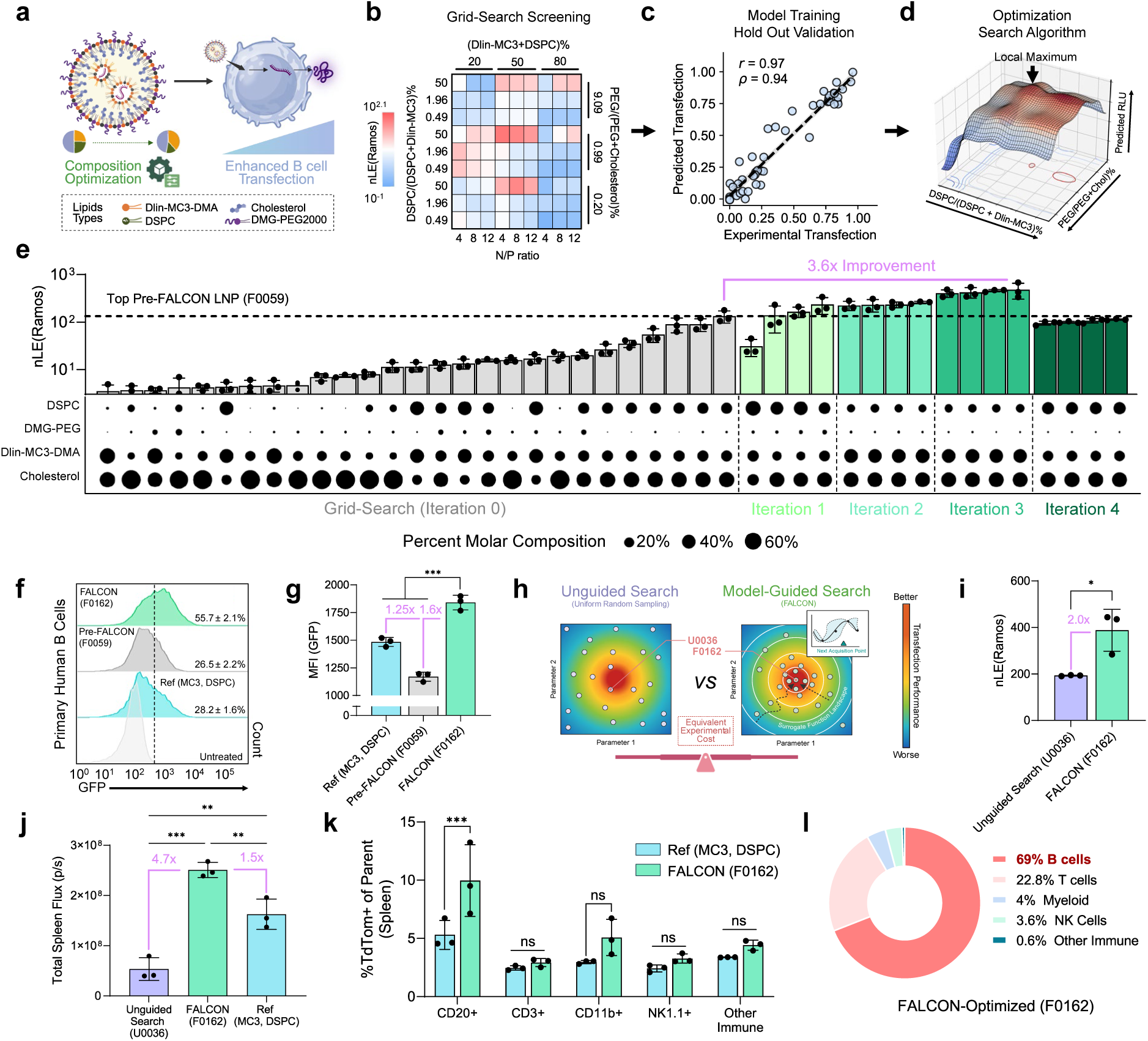
**FALCON-driven single-objective optimization of LNPs for B cell transfection**. **a.** Schematic of the FALCON objective, with lipid types fixed for compositional optimization. **b.** Initial grid-search library screen of 81 LNPs screened in Ramos human B cells. **c.** Representative hold-out validation, illustrating predictive performance of the surrogate model during optimization, in this case evaluated on data up to the final FALCON iteration. **d.** Contour plot showing the predicted transfection landscape used to guide the optimization process. **e.** Bar plot depicting the nLE of formulations tested in the grid-search screen and each subsequent iteration. Dot plots illustrate the percentage molar composition of each component in the displayed LNPs. **f**–**g.** Validation of the transfection efficiency of the top FALCON-optimized LNP against the Onpattro and the top grid-search LNP in primary human B cells, using flow cytometry to quantify the percentage of transfected cells (**f**) and mean fluorescence intensity (MFI) of GFP reporter expression among the transfected cell population (**g**). **h.** Schematic of the search methodology employed by FALCON model-guided search vs an unguided search control via uniform random sampling, **i.** *In vitro* validation of the top LNP compositions identified by FALCON (F0162) against the top composition identified by unguided search (U0036). **j.** Total spleen flux mice dosed with U0036, F0162, and a commercial reference formulation (MC3, DSPC). **k.** Percentage of tdTomato⁺ (transfected) cells among splenic immune subsets following LNP delivery. **l.** Transfection profile of F0162 in the spleen, showing the relative contribution of each immune cell type to the total TdTom⁺ cell population. nLE (normalized luciferase expression) refers to raw readings from *in vitro* luciferase transfection experiments, which were blank-subtracted and batch normalized against a set of internal controls. Data are presented as mean ± SEM from a representative experiment (n=3). The *p*-values were determined via one-way ANOVA with Tukey’s multiple comparisons test (**g, j**), two-tailed Student’s *t*-tests (**i**), and multiple *t*-tests (**k**). ns: *p* > 0.05, \**p* < 0.05, \*\**p* < 0.01, \*\*\**p* < 0.001. **Panels a** and **h** were created with BioRender and released under a Creative Commons Attribution-NonCommercial-NoDerivs 4.0 International license.

To initiate FALCON, a sparse grid-search screen of LNPs was performed in Ramos, an human B cell line. As our purpose was to examine the effects of optimizing formulation composition, the lipid components used were fixed to match those in the FDA-approved Onpattro formulation, *i.e.*, 1,2-distearoyl-sn-glycero-3-phosphocholine (DSPC), cholesterol, DMGPEG-2000, and Dlin-MC3-DMA, which was used as a reference (**Fig. 2a**). This reference formulation was not included in the initial grid-search library or surrogate model training as it was reserved as an external benchmark to enable independent evaluation of the performance of FALCON-driven optimization. Transfection of this initial library using luciferase reporter mRNA, quantified by normalized luciferase expression (nLE) readings, spanned a broad range of 10^-2.3^ to 10^2.1^, with a good distribution of high and low hits **(Fig. 2b)**. This data was preprocessed and used to train the initial surrogate XGBoost model, which was continuously refined after new data was acquired. **Fig. 2c** illustrates a representative holdout validation performance of the trained model, in this case trained on data up to and including the fourth iteration. These surrogate models were also trained on cumulative data generated across optimization campaigns within the same study. These trained models enabled mapping of the LNP design space, as visualized by the contour plot of predicted transfection efficiencies (**Fig. 2d**), allowing search algorithms to predict regions of potent B cell transfection to sample from.

In total, four iterations of FALCON-driven optimization were conducted, with 12 FALCON-suggested LNPs tested every experimental iteration (**Fig. 2e**). Scatter plots displaying all 129 tested formulations can be found in **Supplementary Fig. 8**. In the first iteration, FALCON formulations exhibited higher transfection efficiencies, with an average nLE of 49.1, approximately 6.8-fold higher than that of the original grid-search library. By Iteration 2-3, FALCON-generated formulations consistently achieved nLE values that surpassed the top formulation from grid-search screening. Principal component analysis (PCA) visualizations of the search space in two dimensions illustrate that the optimized formulations suggested by FALCON occupy a distinct, expanded composition space than that of the original grid-search library (**Supplementary Fig. 9**).

Of particular note, nLE values from the final fourth FALCON iteration decreased relative to the previous iteration. This dip in performance, when preceded by sustained improvement and convergence of suggested compositions, is consistent with diminishing returns following extensive sampling of high-performing regions within the explored formulation space (**Supplementary Note 1**). These observations suggest that additional iterations were unlikely to yield substantial performance gains under the current surrogate and acquisition framework. Accordingly, this behavior was used as a practical stopping criterion for iterative optimization.

To validate the performance of LNPs identified through FALCON, we evaluated the *ex vivo* transfection of our top FALCON-optimized LNP (F0162) in primary human B cells obtained from healthy donors. F0162 achieved markedly higher transfection efficiency (2.0-and 2.1-fold) and MFI among transfected cells (1.2-and 1.6-fold) compared to the reference and top grid-searched Pre-FALCON LNP (F0059), respectively (**Fig. 2f, g, Supplementary Fig. 12**). Because all three formulations contained the same lipid components, this enhanced potency was attributed to differences in composition.

FALCON employs a model-guided strategy to efficiently sample areas of the search space with the highest transfection potential. To demonstrate such an approach facilitates improved performance compared to simple naïve searches, we introduced an unguided search benchmark, obtained through uniform random sampling (**Fig. 2h, i, Supplementary Fig. 13**). For this control, 129 formulations were randomly sampled, corresponding to the number of formulations tested during initial library screening and 4 iterations of FALCON-driven B cell optimization. We also allowed random search to sample within the same expanded parameter bounds FALCON had discovered. In *in vitro* transfection assays, F0162 achieved 2-fold greater nLE than the top unguided search composition (U0036) we found, a result that supports FALCON’s ability to identify high-performing compositions more effectively than passive approaches.

Finally, to characterize the *in vivo* delivery performance of F0162, we administered mFluc-loaded LNPs intravenously at a dose of 10 µg mRNA (0.5 mg/kg) and harvested the spleen after 6 h for *ex vivo* luciferase assay. All *in vivo* tested LNP compositions are shown in **Supplementary Table 5**, with their respective size, PDI, zeta potential, and RNA encapsulation reported in **Supplementary Tables 6 and 7**. F0162 LNPs achieved 4.7-fold greater mFluc expression in the spleen than U0036, and 1.5-fold greater than the reference (**Fig. 2j)**. Similar transfection enhancement trends were also observed in the liver (**Supplementary Fig. 14**). Based on these results, we assessed *in vivo* single-cell transfection efficiency using a Cre-Ai9 reporter model dosed with 10 µg mRNA per mouse (0.5 mg/kg). In the spleen, F0162 transfected 9.97 ± 1.78% of CD20^+^ B cells, a 1.8-fold improvement over the reference (**Fig. 2k, Supplementary Figs. 15 and 16**). In addition, we observed no substantial difference in transfection among CD3^+^, CD11b^+^, NK1.1^+^, and other immune cell subsets, suggesting that the increased spleen signal of F0162 is mediated primarily by increased B cell delivery. On average, B cells made up approximately 69% of the total transfected cells in the spleen in the F0162-treated mice (**Fig. 2l**). We also examined single-cell transfection in the liver compartment, where we observed high overall transfection of non-immune cells in the liver, at levels similar to the reference (**Supplementary Figs. 17-19**). This was not surprising, given minimizing off-target transfection was not accounted for in our single-objective optimization workflow. This result justified the necessity of adopting a multi-objective strategy in order to enhance selective B cell delivery.

### Multi-objective FALCON optimization balances competing objectives to uncover LNPs with enhanced cell type-selectivity

To test the multi-objective capabilities of our platform, we deployed FALCON to guide the design of LNPs for selective transfection of Ramos B cells (target to maximize) over HepG2 cells (target to minimize), a hepatocyte cell line (**Fig. 3a**). This task is challenging for traditional grid search screening methods, as the response landscape between the two cell types is not clearly separable, making it hard to isolate a “B cell-selective” composition space without computational guidance (**Supplementary Fig. 20**).

**Figure 3.**
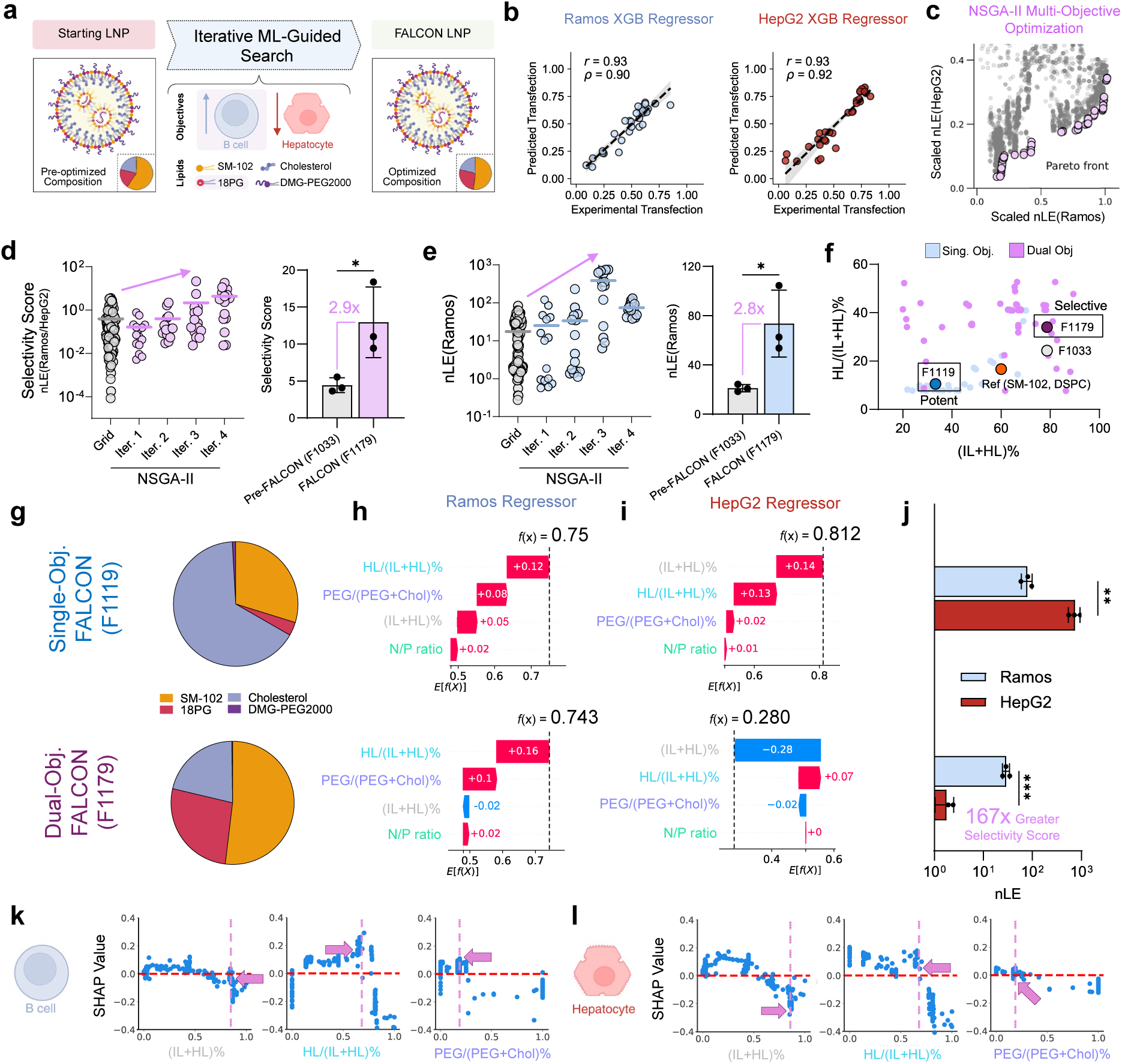
Multi-objective optimization of anionic LNPs for optimal B cell and minimal hepatocyte delivery. **a.** FALCON was applied to a 4-component anionic helper lipid LNP system to co-optimize for high Ramos B cell and low HepG2 hepatocyte delivery. **b**. Representative hold-out performance of both Ramos B and HepG2 transfection models. **c.** Representative Pareto front illustrating optimal trade-off solutions for selective Ramos B cell transfection identified via NSGA-II. **d.** Dot plot of selectivity scores achieved by LNPs tested in the initial grid-search screen and four subsequent FALCON-guided iterations. FALCON dual-objective optimized LNP F1179 is more selective than the most selective Pre-FALCON LNP from the initial grid-search (F1033). **e.** Dot plot of nLE values achieved by LNPs tested in the initial grid-search library and four subsequent FALCON-guided iterations. F1179 achieves a higher nLE in Ramos B cells than F1033. **f.** Visualization of formulations explored by single-and dual-objective FALCON algorithms, highlighting the reference benchmark, top FALCON single-objective optimized F1119, F1033, and F1179 compositions based on their HL/(IL + HL)% and IL + HL% parameters. **g.** Lipid percent molar compositions of the top potent F1119 and selective F1179 FALCON optimized LNPs. **h–i.** Waterfall plots from SHAP analyses of F1119 and F1179 showing the contribution of each parameter to the predicted normalized transfection in Ramos B cells (**h**) and in HepG2-trained models (**i**). **j.** Side-by-side comparison of the *in vitro* nLE of F1119 and F1179 in Ramos and HepG2 cell lines. **k–l.** SHAP dependence plots showing the correlation between each compositional parameter and its corresponding SHAP value in Ramos B (**k**) and HepG2 (**l**) models. Pink vertical dashed lines and an arrow indicate the feature values of the optimized F1179 LNP. nLE (normalized luciferase expression) refers to raw readings from *in vitro* luciferase transfection experiments, which were blank-subtracted and batch normalized against a set of internal controls. Selectivity score was calculated as the fold change in nLE of the target cell type over the off-target cell type. Data are presented as mean ± SEM from a representative experiment (n = 3). The *p*-values were determined via multiple *t*-tests with Benjamini-Krieger-Yekutieli correction (FDR Q = 1%) (**j**), and unpaired Student’s *t*-tests (**d**, **e**). \**p* < 0.05, \*\**p < 0.01,* \*\*\**p* < 0.001. **Panel a** was created with BioRender.com and released under a Creative Commons Attribution-NonCommercial-NoDerivs 4.0 International license.

We began with a parallel library screen of 108 SM102 and 18PG based LNP compositions in Ramos B and HepG2 cells (**Supplementary Table 2**). The resulting dataset and trained surrogate models were then fed into both FALCON’s multi-objective computational pipeline. **Fig. 3b** depicts representative hold-out performances for Ramos B cell and HepG2 cell transfection models. **Fig. 3c** depicts a representative Pareto front generated from the multi-objective NSGA-II search algorithm. The plot depicts a curved boundary formed by Pareto-optimal points (depicted in pink), indicating meaningful trade-offs between Ramos B and HepG2 transfection performance. Over four iterative cycles of multi-objective optimization, FALCON designed various LNPs with improved selectivity scores for Ramos B over HepG2 cells (**Fig. 3d**), with the top-nominated LNPs, F1179, demonstrating a 2.9-fold improvement over the top-selective Pre-FALCON F1033 formulation. We also observed improved *in vitro* transfection potency levels in FALCON-suggested LNPs with F1179 achieving a 2.8-fold improvement compared to Pre-FALCON F1033, demonstrating that FALCON can simultaneously search for selective designs while maintaining or improving on-target potency (**Fig. 3e**). For comparison, single-objective optimization for B cells was also performed, resulting in an optimized formulation F1119 (**Supplementary Fig. 21**).

For both single and dual optimization tasks, FALCON explored formulations outside of and between the initial grid-search composition space (**Supplementary Fig. 22**). More importantly, the optimal compositions identified for each task were distinct from each other. Regions explored by single-vs. dual-objective algorithms differed in key compositional parameters, suggesting different formulation requirements for selectivity optimization compared to simply maximizing target transfection (**Fig. 5f**). Specifically, we observed that low (IL + HL)% and lower HL/(IL + HL)% were more explored during single-objective optimization, while increased (IL + HL)% and HL/(IL + HL)% were explored more during Ramos B selective optimization.

Post hoc SHAP analysis was performed to interpret the final trained Ramos B and HepG2 models (**Supplementary Fig. 23**). SHAP was first leveraged to conduct local feature importance analysis of F1119 and F1179. **Fig. 3g** shows the molar composition of these LNPs (**Supplementary Table 5**). Results from local SHAP analysis are displayed as waterfall plots that visualize the direction and magnitude of each feature value’s impact on transfection in Ramos B cells (**Fig. 3h**) and HepG2 cells (**Fig. 3i**).

Notably, all parameter values in F1119 were optimized to increase Ramos B cell transfection (**Fig. 3h**). On the other hand, the higher (IL + HL)% of F1179 actually decreased the predicted Ramos B cell transfection, although only minimally (-0.02); but it is also this choice that substantially minimized the predicted HepG2 cell transfection of the LNP (by-0.28) (**Fig. 3h, i**). This result suggests that when FALCON was fitted with a multi-objective optimization strategy, it made strategic trade-off decisions, selecting parameter values that, while slightly decreasing Ramos B cell transfection, greatly reduced HepG2 cell transfection, ultimately enhancing overall selectivity by 167-fold *in vitro* (**Fig. 3j**). This tradeoff is not considered in the single-objective case. **Fig. 3k, l** displays Ramos B and HepG2 model SHAP values as a function of critical design parameter values and highlights the rational decisions made by FALCON to design the F1179 selective LNP (pink). Interestingly, these decisions were based not only on differences in preferred parameter values but also differences in the magnitude of each feature value’s impact on the transfection in the two cell types.

### FALCON-optimized LNPs mediate robust and selective transfection in splenic B cells *in vivo*

To characterize *in vivo* performance, mFluc-loaded LNPs were dosed intravenously at 10 µg per mouse (0.5 mg/kg) into C57/B6 mice, and their liver and spleen were harvested after 6 h for *ex vivo* luciferase assay. F1119 (SM102, 18PG) and F1179 (SM102, 18PG) both displayed robust delivery to the spleen. Notably, F1179 (SM102, 18PG) achieved a 75-fold lower off-target liver transfection compared to the reference (SM102, 18PG) LNP (**Fig. 4a, b**), reflecting about 10-fold biased transfection in the spleen over the liver. This is markedly more spleen-tropic than both the reference (SM102, 18PG) LNP (28-fold improvement) and F1119 (SM102, 18PG) (3.5-fold improvement) (**Fig. 4c**).

**Figure 4.**
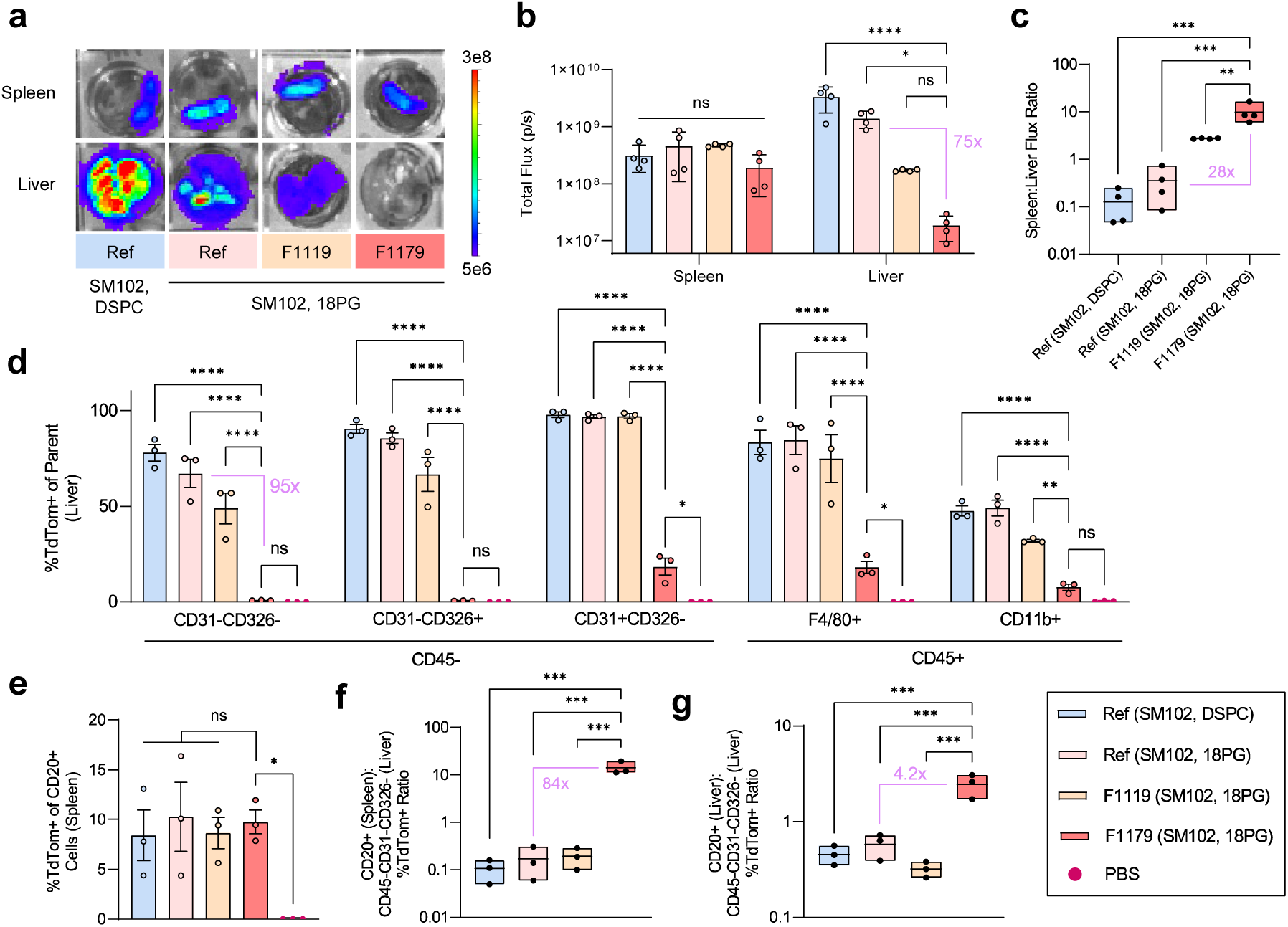
*In vivo* delivery of FALCON-optimized F1119 and F1179. **a.** *Ex vivo* luciferase assay by IVIS imaging for transfection in the spleen and liver, after intravenous dosage of 10 µg (0.5 mg/kg) mLuc-loaded LNPs in mice. Representative images are shown. **b.** Whole organ *ex vivo* luminescence flux measurements in the spleen and liver. **c.** The calculated total flux ratio between the spleen and the liver. **d**–**g.** Ai9 transgenic model where mice were dosed with Cre mRNA-loaded LNPs. The spleen and liver were harvested after 3 days for flow cytometry analysis (gating strategy is shown in **Supplementary Figs. 15–18**). **d.** Non-immune CD45^-^ cell and myeloid lineage transfection in the liver, where EpCAM^+^ epithelial cells are CD45^-^CD31^-^CD326^+^, endothelial cells are CD45^-^CD31^+^CD326^-^, other non-immune cells (consisting predominantly of hepatocytes) are CD45^-^CD31^-^CD326^-^, Kupffer cells are CD45^+^F4/80^+^, and other myeloid lineage cells are CD45^+^CD11b^+^. **e.** CD20^+^ B cell transfection in the spleen. **f–g.** Single-cell delivery selectivity comparing CD20^+^ cell transfection in the spleen (**f)** or liver (**g**) to CD45^-^CD31^-^CD326^-^ liver cell transfection. Data are presented as mean ± SEM from a representative experiment (n = 3 or 4). *p*-values were determined via one-way or two-way ANOVA followed by Šídák’s (**b**), Tukey’s (**c**) or Dunnett’s (**d–f**) multiple comparisons test. ns: *p* > 0.05, \**p* < 0.05, \*\**p* < 0.01, \*\*\**p* < 0.001, \*\*\*\**p* < 0.0001.

**Figure 5.**
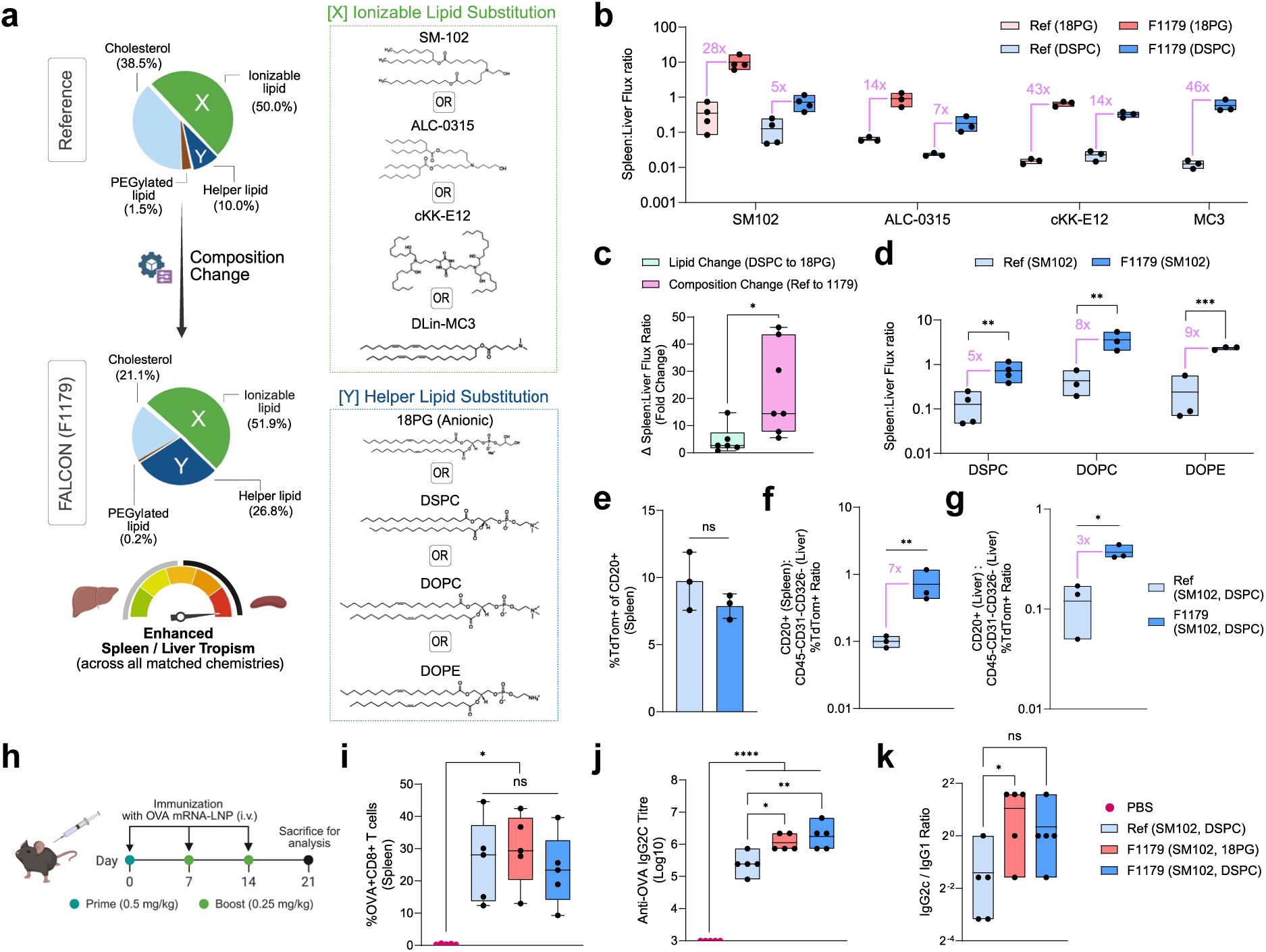
F1179 enables spleen-tropic delivery across several lipid structures and enhances vaccine responses. **a.** Design of factorial ionizable and helper lipid substitutions to investigate the effects of the F1179 composition across several lipid chemistries. **b.** Organ-level spleen-tropism of F1179 and reference composition LNPs across a panel of ionizable lipids and containing either 18PG or DSPC helper lipids. The Spleen-to-Liver flux ratio is calculated from the total bioluminescence flux measured for each organ using ex vivo luciferase assays and IVIS. **c.** Relative contributions to spleen-tropism of chemistry changes (switching DSPC to 18PG, composition held constant) and compositional changes (switching reference for F1179, lipid chemistry held constant). **d.** Organ-level spleen-tropism of SM102-based F1179 and reference composition LNPs containing either DSPC (shared data as in Fig. 5b, shown for comparisons), DOPC, or DOPE zwitterionic helper lipids. **e–g.** Single-cell transfection of SM-102 and DSPC-based LNPs varying composition (reference or F1179) dosed at 0.5 mg/kg. The gating strategy used is shown in **Supplementary Figs. 15–18.** Single-cell delivery potency comparing CD20^+^ cell transfection in the spleen (**e**). Single-cell delivery selectivity comparing CD20^+^ cell transfection in the spleen (**f)** or liver (**g**) to CD45^-^CD31^-^CD326^-^liver cell transfection. **h–k.** Systemic vaccination of mice with mOVA-LNPs and evaluation of adaptive immune response. Vaccination and experimental timeline (**h**). Day 21 unstimulated splenocytes were assessed via flow cytometry to determine the percentage of OVA-specific CD3^+^CD8^+^ T cells (by SIINFEKL-H-2Kb tetramer staining) (**i**). The gating strategy used is shown in **Supplementary Fig. 31. j–k.** Titers of OVA-specific serum IgG2c on day 21 (**j**) and IgG2c to IgG1 titer ratio (**k**) determined by ELISA. Data are presented as mean ± SEM from a representative experiment (n = 3 or 4). The *p*-values were determined via unpaired t-test (**c, e–g**), one-way or two-way ANOVA followed by Sidak’s (**d**) or Tukey’s multiple comparisons (**b**, **i–k**) test. ns: *p* > 0.05, *\*p* < 0.05, \*\**p* < 0.01, \*\*\**p* < 0.001, \*\*\*\**p* < 0.0001. SM102/18PG-based reference and F1179 datapoints shown in **b** and **c** were previously presented in Fig. 4 and are reproduced here for comparison with the expanded lipid chemistry panel. **Panels a** and **h** were created with BioRender, and released under a Creative Commons Attribution-NonCommercial-NoDerivs 4.0 International license.

We also characterized cell type-specific delivery in an Ai9 transgene murine model dosed at 10 µg per mouse (0.5 mg/kg). F1179 (SM102, 18PG) minimally transfected off-target cell populations in the liver, including CD45-CD31^+^ endothelial, CD45^-^CD326^+^ epithelial, and CD45^-^CD31^-^CD326^-^ cells liver cells, a population expected to consist predominantly of hepatocytes, compared to the other tested LNPs (**Fig. 4d**). Specifically, F1179 (SM102, 18PG) achieved a 95-fold reduction in off-target CD45^-^CD31^-^CD326^-^ cell transfection (<1% TdTom^+^) compared to the reference (SM102, 18PG) LNP, a difference attributable to FALCON-driven compositional changes. F1179 (SM102, 18PG) mediated robust splenic CD20^+^ cell transfection levels (∼10% TdTom^+^), although with lower levels of liver CD20^+^ cell transfection (**Fig. 4e, Supplementary Fig. 24-25**). Importantly, F1179 (SM102, 18PG) achieved more than 14:1 splenic CD20^+^ to liver CD45^-^CD31^-^CD326^-^ transfection ratio (*e.g.*, for every 14% of spleen CD20^+^ cells transfected, 1% of liver CD45^-^CD31^-^CD326^-^ are transfected), an 80-fold improvement for B cell specific delivery compared to the reference (SM102, 18PG) and 70-fold improvement compared to F1119 (SM102, 18PG) (**Fig. 4f**). When comparing the on-target liver CD20^+^ cell to off-target cell delivery, F1179 (SM102, 18PG) achieved more than 2:1 CD20^+^ to CD45^-^CD31^-^CD326^-^ transfection ratio in the liver, representing a 4-fold improvement for B cell selectivity compared to the reference (SM102, 18PG) (**Fig. 4g**).

### F1179 enables spleen-tropic behavior across several ionizable and helper lipid chemistries and enhances vaccine responses

Given the previously reported spleen-tropic nature of anionic helper lipid-containing LNPs (*e.g*. SORT LNPs), we investigated the contributions of chemical versus compositional changes towards the observed enhanced selectivity of F1179. We did this by testing the F1179 composition across factorial panels of ionizable lipid (IL) and helper lipid (HL) chemistries (anionic and zwitterionic) (**Fig. 5a)**. In the first factorial panel, we varied combinations of ionizable lipid (SM102, MC3, ALC-0315, cKK-E12) and helper lipid (anionic 18PG, zwitterionic DSPC) chemistries across the reference composition and the FALCON-identified F1179 composition. Following intravenous delivery, F1179 LNPs at 10 µg per mouse (0.5 mg/kg) displayed enhanced spleen-tropism across all IL and HL chemical combinations tested compared to their reference composition counterparts by >5-fold and up to 46-fold (**Fig. 5b, Supplementary Fig. 26, Supplementary Table 8)**. A comparative analysis of the chemical and compositional contributions on spleen-tropism within the tested LNPs uncovered that compositional changes (swapping composition from reference to F1179 while keeping helper lipid constant) displayed on average markedly stronger improvement in spleen-tropism than helper lipid chemical changes alone (swapping DSPC to 18PG while keeping composition constant) (**Fig. 5c**). We then investigated whether F1179 composition-based spleen tropism was also observed across other commonly used zwitterionic helper lipid chemistries (DOPC and DOPE). Our results show that the F1179 composition consistently improved spleen-tropism by more than 5-fold compared to reference compositions containing the same helper lipid (**Fig. 5d, Supplementary Fig. 27**).

Next, using Ai9 transgenic mice, we assessed single-cell transfection to see if the enhanced cell type selectivity observed for anionic F1179 (SM102, 18PG) LNPs were also present for DSPC zwitterionic formulations (SM102, DSPC). LNPs were formulated with the reference or F1179 compositions and dosed intravenously at 10 µg per mouse (0.5 mg/kg). Compared to the reference composition (SM102, DSPC), F1179 (SM102, DSPC) achieved similar levels of splenic B cell transfection with markedly lower off-target liver non-immune cell transfection (**Fig. 5e, Supplementary Fig. 28-30**). Resultantly, F1179 (SM102, DSPC) achieved improved splenic B cells transfection selectivity by 7-fold compared to the reference (**Fig. 5f**). Similarly, in the liver tissue, F1179 (SM102, DSPC) achieved a 3-fold enhancement in selective transfection of B cells (**Fig. 5g**). While these results mainly demonstrate transferability on a formulation level, interestingly, SHAP plots from models trained on SM102-based datasets using either DSPC or 18PG helper lipids both show similar predicted trends for several compositional features (**Supplementary Fig. 23**). We also note that several F1179 formulations exhibited larger particle sizes than their reference counterparts, whereas others retained similar physicochemical characteristics (**Supplementary Tables 6 and 7)**. These findings suggest that no single physicochemical parameter consistently explains the observed selectivity trends across lipid systems and that targeted studies will be required to establish causal relationships.

To examine the therapeutic potential of F1179 LNPs, we performed an OVA vaccination study in healthy mice, comparing the immune response generated by F1179 (SM102, 18PG) and F1179 (SM102, DSPC) to that of the reference LNPs (SM102, DSPC) (**Fig. 5h**). F1179 formulations generated similar percentages of antigen-specific CD8^+^ T cells in the spleen as the reference LNPs, indicating that liver de-targeting and modest changes in on-target cell transfection did not compromise T cell activation (**Fig. 5i, Supplementary Fig. 31)**. Evaluation of serum anti-OVA IgG displayed that F1179 LNPs achieved more than 3-fold higher IgG2c titers and an increased IgG2c/IgG1 ratio relative to the reference formulation, suggesting a stronger Th1-skewed immune response associated with cell-mediated immunity (**Fig. 5j and k**, **Supplementary Fig. 32**). These findings are consistent with the possibility that reduced antigen expression in the tolerogenic liver microenvironment may contribute to improved vaccine responses^51^. Together, these results support the potential translational relevance of FALCON-optimized formulations for vaccine and immunotherapy applications.

### FALCON identifies enhanced compositions for biological contexts beyond B cell delivery

To demonstrate the utility of the FALCON platform for applications beyond B cell delivery, we deployed FALCON to optimize LNP compositions for myeloid cell-selective delivery relevant to vaccine and CAR-immune cell applications (**Fig. 6**). FALCON was configured to optimize transfection in THP-1 human monocytes and minimize transfection in HepG2 human hepatocyte cells (**Fig. 6a, Supplementary Fig. 33**). We began optimization with an initial grid-search library of 36 unique LNPs, and conducted iterative search in batches of 20 LNPs: 10 suggested by NSGA-II (optimize LNPs for selectivity) and 10 proposed by I-optimal algorithms (minimize average predictive variance), to demonstrate the possibility of integrating sampling methods that focus on model uncertainty, allowing efficient learning from a comparatively sparser starting dataset (**Supplementary Table 9, see Methods**).

**Figure 6.**
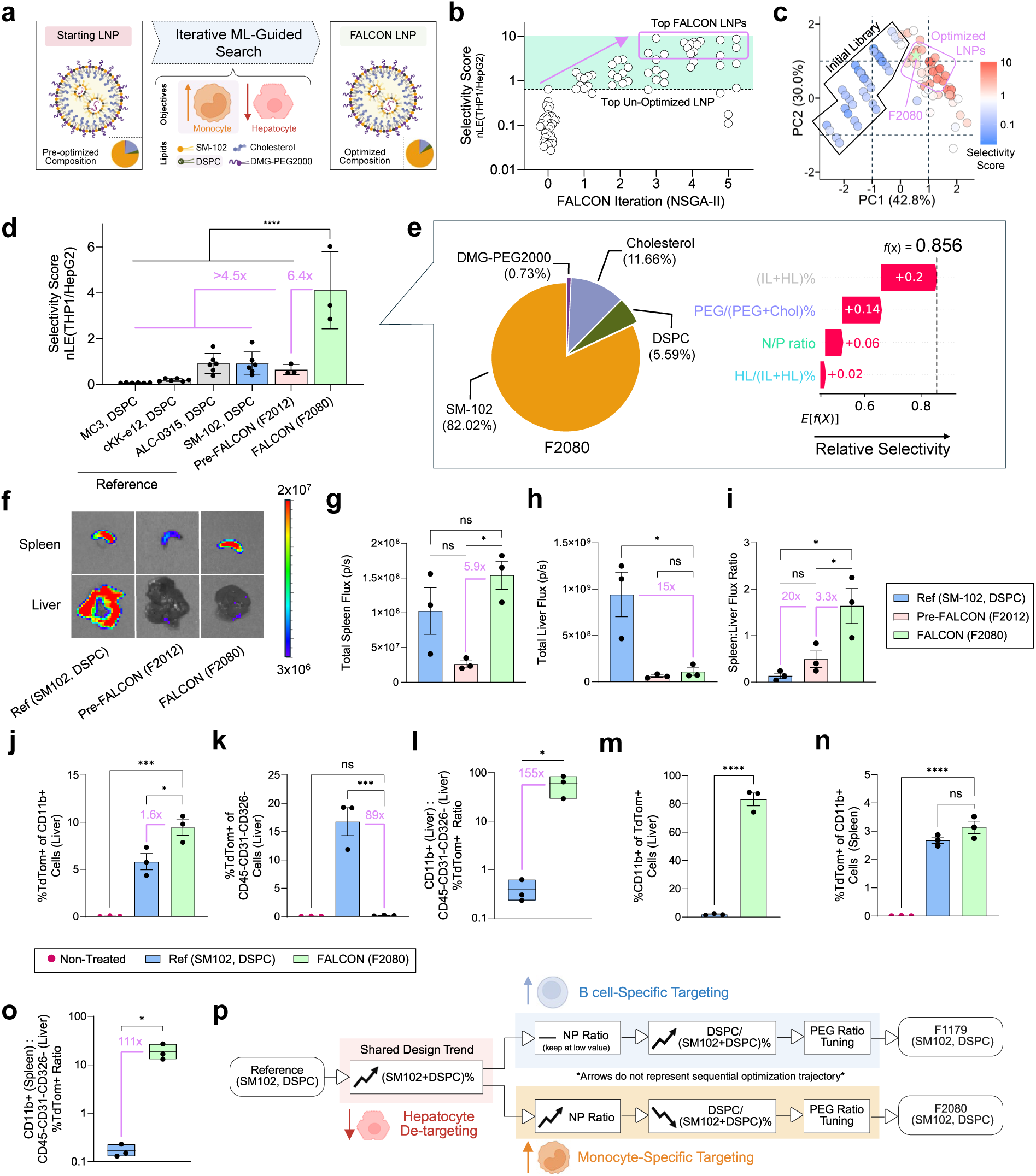
FALCON-guided LNP optimization for selective delivery to myeloid cells. **a.** Schematic of dual-objective FALCON optimization for high myeloid and minimal hepatocyte delivery, depicting optimization targets and lipid types used. **b.** Iterative scatter plot of the selectivity score of LNPs screened in THP-1 monocytes and HepG2 hepatocyte cell lines during each FALCON iteration. Formulations that lie in the green region demonstrate enhanced selectivity over the top Pre-FALCON formulation from the initial grid-library library (iteration 0). **c.** PCA visualization of all tested LNP compositions, colored by selectivity score. The search spaces encompassed by the initial grid-search library and by FALCON optimized LNPs are labelled. **d.** Side-by-side comparison of the *in vitro* selectivity score of the FALCON-optimized formulation (F2080) against commercial reference benchmarks, and the top Pre-FALCON optimized LNP (F2012). **e.** Percent molar composition of F2080 and the predicted contribution of each of its individual feature values towards its enhanced selectivity score (waterfall plot). **f.** Representative *in vivo* luciferase imaging of the spleen and liver in B6 mice injected intravenously with F2080, F2012, and the reference benchmark. **g–i.** Quantified total spleen flux (photons/sec) (**g**), total liver flux (photons/sec) (**h**), and the spleen-to-liver flux ratio (**i**) following LNP delivery. **j–o.** Single-cell transfection in Ai9 mice of F2080 and the reference LNPs dosed at 0.25 mg/kg. The gating strategy used is shown in **Supplementary Figs. 15–18.** Transfection of CD11b^+^ **(j)** and CD45^-^CD31^-^CD326^-^ **(k)** cells in the liver and the ratio of the two transfected cell types (**l**). Percentage of CD11b^+^ cells among all TdTom^+^ cells in the liver, indicating transfection specificity in the liver microenvironment (**m**). CD11b^+^ cell transfection in the spleen (**n**) and the ratio of transfected splenic CD11b^+^ cells to CD45^-^CD31^-^CD326^-^ liver cells (**o**). **p.** Summary of SHAP analysis trends displaying design rules driving liver de-targeting, B-cell-specific targeting, and myeloid-specific targeting. The nLE (normalized luciferase expression) refers to raw readings from *in vitro* luciferase transfection experiments, which were blank-subtracted and batch normalized against a set of internal controls. Selectivity score was calculated as the fold change in nLE of the target cell type over the off-target cell type. The nLE values from Fig 6b, c are displayed on a logarithmic scale, to allow visualization of the full dataset distribution. Data are presented as mean ± SEM from a representative experiment (n = 3). *p*-values were determined via unpaired t-test (**l–m, o**) or one-way ANOVA with Tukey’s (**d, g–h**) or Dunnett’s (**j–k, n**) multiple comparisons test. ns: *p* > 0.05, \**p* < 0.05, \*\*\**p* < 0.001, \*\*\*\**p* < 0.0001. **Panels a** and **p** were created with BioRender, and released under a Creative Commons Attribution-NonCommercial-NoDerivs 4.0 International license.

After 5 iterations, a plateau in LNP performance was observed (iterations 3–5), corresponding to the identification of a region of the design space with improved myeloid selectivity, which was outside of the initial library space (**Fig. 6b, c**). The top selected FALCON LNP (F2080) demonstrated 6.5-fold enhanced *in vitro* transfection selectivity for THP-1 cells over HepG2 cells compared to the top selective pre-FALCON LNPs (F2012) and over 4.5-fold improved selectivity over reference LNPs (**Fig. 6d**). Local SHAP analysis of F2080 suggest that high total (IL+HL)% content (82.02% of all lipids) as well as its optimized PEG/(PEG+Chol)% (5.59%) likely played the most substantial roles in enhancing THP-1 cell-selective transfection (**Fig. 6e**). To investigate *in vivo* delivery profiles, we treated C57/BL6 mice with 5 µg per mouse (0.25 mg/kg) of mLuc-loaded LNPs intravenously (**Fig. 6f**). We observed similar spleen transfection signal in F2080 LNPs compared to reference LNPs (with matched lipid chemistries SM102 and DSPC) and enhanced delivery compared to F2012 (5.9-fold) (**Fig. 6g)**. Both Pre-FALCON and F2080 LNPs displayed 15-fold lower liver transfection compared to the reference (**Fig. 6h**). As a result, F2080 achieved markedly improved spleen selectivity over the liver, with a 3.3-fold and 20-fold selectivity improvement compared to Pre-FALCON and reference LNPs, respectively (**Fig. 6i**).

Given the promising organ-level results, we compared the single-cell transfection of F2080 against the reference LNP. Ai9 mice were dosed intravenously at 0.25 mg/kg, and the spleen and liver were harvested 24 h afterward for flow cytometry analysis. In the liver, F2080 achieved robust CD11b^+^ myeloid cell transfection, while displaying minimal off-target delivery (<1%) in CD45^-^CD31^-^CD326^-^ cell populations. Specifically, F2080 achieving a 1.6-fold increase in CD11b^+^ myeloid cell transfection and over 89-fold reduction in off-target liver cell transfection, corresponding to a 155-fold improvement in delivery selectivity (**Fig. 6j–l, Supplementary Figs. 34 and 35)**. Analysis of the cellular composition of the transfected (TdTom^+^) population revealed that among all TdTom^+^ harvested cells in the liver, CD11b^+^ cells represented over 80% of the TdTom^+^ cells in F2080 treated mice livers, while this number was less than 3% in reference LNP-treated mice livers (**Fig. 6m)**. In the spleen, F2080 achieved similar myeloid transfection to reference LNPs, leading to a 111-fold higher ratio of transfected splenic myeloid cells to CD45^-^CD31^-^CD326^-^ liver cells (**Fig. 6n, o, Supplementary Fig. 36**). We also observed a more myeloid-biased transfection profile within the spleen, as F2080 mediated a similar level of CD11b^+^ transfection as the reference LNPs, but a substantially lower level of transfection in splenic CD20^+^ cells (**Supplementary Fig. 36**). Together, these analyses demonstrate that F2080 exhibits a strongly myeloid-biased transfection profile both within the liver and spleen microenvironment. To better understand the factors underlying the selective delivery phenotypes identified by FALCON, we next analyzed composition-dependent design features and biodistribution behavior.

Comparing FALCON’s designs for B cell (F1179) and myeloid-selective (F2080) LNP compositions via SHAP, we identified design rules for SM102-and DSPC-based LNPs that de-target the liver and improve cell-specific delivery (**Fig. 6p, Supplementary Figs. 37 and 38**). We found that increasing the combined molar percentage of SM102 and DSPC, corresponding to decreased cholesterol content, was associated with liver de-targeting. This observation is consistent with previous studies showing that reduced cholesterol content can diminish ApoE-mediated hepatic delivery^52^. Importantly, within this shared liver de-targeting composition space, FALCON optimization further tuned N/P ratio and DSPC content to drive transfection either towards B cells (high DSPC content, a property recently shown to enhance delivery to marginal zone B cells^53^) or myeloid cells (low DSPC content and high N/P ratio). The stronger preference of THP-1 cells for a high N/P ratio may be due to greater expression of scavenger receptors, which are primed to internalize more cationic particles^54,55^. Additionally, we characterized the organ-level biodistribution of the top FALCON LNPs and found they share a similar distribution with the reference LNPs, including in the liver, where we observed the most significant transfection differences. These findings are consistent with the possibility that differences in cell-specific uptake and/or intracellular processing contribute to the observed delivery profiles (**Supplementary Fig. 39**).

Finally, to demonstrate FALCON’s applicability outside of immune cell delivery, we also conducted a preliminary optimization of LNPs for delivery to pancreatic β cells (**Supplementary Result 1**)^56^. FALCON was also used to perform *in vitro* optimization for (1) selective transfection of Ramos B and THP-1 cells against each other, to compare compositional features distinguishing B cell-and monocyte-selective transfection, and (2) a quadruple-objective optimization exercise to explore the feasibility of extending FALCON-guided formulation search to the many-objective case (**Supplementary Result 2**).

## DISCUSSION

The therapeutic potential of gene carriers depends on the ability to engineer delivery systems that efficiently transfect biological targets while avoiding off-target effects^14^. FALCON presents a practical realization of this design philosophy. By integrating machine learning and multi-objective algorithms into a closed loop workflow, FALCON enables efficient exploration and optimization of cell-type selective LNP compositions. We demonstrate that FALCON-optimized LNPs, obtained from 3 to 5 iterations in under 2 weeks, achieve improved performance over commercial and grid-search benchmarks in predefined objectives, underscoring the utility of this framework.

For our initial application, we employed FALCON to optimize LNP composition to target human B cells. Engineered B cells represent a promising and yet underutilized therapeutic platform for human diseases, in part due to their traditionally difficult-to-transfect nature. Reprogramming of B cells via potent mRNA delivery *in situ* promises to provide a scalable and safe approach to express therapeutic payloads, such as broadly neutralizing antibodies against HIV and cancer vaccine antigens^50,57,58^. Furthermore, selectivity-optimized delivery of toxic payload templates to B cell cancers may offer opportunities to overcome the limitations of protein toxins whose clinical utility has been curtailed by off-target toxicity after systemic administration^59^. Beyond B cells, proof-of-concept FALCON datasets for two potential applications, developing LNPs to (1) target myeloid cells for systemically delivered mRNA vaccines or CAR-constructs and (2) transfecting pancreatic β cells for T1D autoimmune disease, were briefly explored.

The surrogate models used in FALCON were intentionally selected to balance predictive accuracy, computational efficiency, and interpretability in a low-data iterative optimization setting. For the composition-based datasets investigated here, tree-based ensemble methods like XGBoost provided sufficient predictive performance while enabling rapid retraining and model interpretation. FALCON shares conceptual similarities with traditional DOE-based formulation optimization approaches in that both seek to identify formulations with desired biological performance while minimizing experimental burden. However, whereas DOE is commonly used to characterize formulation-response relationships and identify promising design regions, FALCON combines machine-learning models with closed-loop Pareto optimization to adaptively navigate formulation space and quantitatively map trade-offs between competing delivery objectives. This framework enables iterative refinement of formulations based on experimental feedback while capturing complex nonlinear relationships between composition and biological performance.

FALCON-optimized LNPs achieved enhanced cell-type selectivity in B cell-and myeloid cell-targeted case studies, both across organ compartments (spleen versus liver) and within the liver microenvironment. Both forms of selectivity are important for systemic delivery applications. Notably, FALCON identified a formulation that simultaneously increased CD11b^+^ cell transfection potency (1.6-fold) and selectivity (155-fold) within the liver. For our B cell-targeted formulation, improved selectivity was achieved primarily through reduced off-target liver transfection with minimal changes in on-target activity. Our findings suggest that the relationship between selectivity and potency was formulation dependent. Gains in selectivity do not necessarily require reductions in potency but rather depend on the biological targets and formulation space being explored. From a translational perspective, the significantly lower off-target transfection achieved by our FALCON-optimized LNPs may reduce side effects, improving the therapeutic index of these formulations^60^. Our results suggest that enhanced delivery selectivity may also increase vaccine potency, potentially by limiting antigen expression within the tolerogenic hepatic microenvironment^51^.

An important feature of FALCON is its ability to function effectively with sparse starting datasets, enabling efficient optimization without the need for prohibitively large combinatorial libraries. FALCON’s sampling efficiency thus makes it well-suited for resource-constrained applications such as hard-to-culture primary cells or even direct *in vivo* screening setups. This presents a future direction to strengthen translational alignment, a limitation of our current work, and other high-throughput screening studies that employ a staged setup to select candidate formulations before *in vivo* validation. Currently, the lightweight surrogate models FALCON employs are trained for specific optimization tasks and are not expected to generalize broadly across chemically distinct formulation spaces. Although our cross-lipid validation experiments showed that an optimized formulation displayed similar delivery profiles across several related lipid systems, these results should not be interpreted as evidence of model-level generalization or broadly transferable design rules. Rather, FALCON is intended as a flexible optimization framework that can be readily redeployed to accelerate refinement for new delivery applications, using data generated within a relevant design space. In this context, the value of FALCON lies not in establishing universal formulation principles, but in reducing the experimental burden required to identify high-performing formulations for a given delivery objective. We anticipate that the FALCON framework may be adaptable to the design of delivery systems across payload applications (*e.g.*, plasmid DNA, mRNA, siRNA, gRNA, etc.)^18,27,62–64^, and therapeutics such as cell engineering (*e.g.*, CAR-T, CAR-NK cells)^65,66^, mRNA LNP-based vaccine development^67,68^, and therapeutic protein production^69^.

Under the current LNP screening paradigm, efforts are generally directed toward optimizing delivery to a singular cell target. Our results suggest that this approach may be insufficient in contexts that require greater selectivity. Through SHAP analysis, we uncovered quantitative feature design rules governing cell-selective transfection, identifying both shared tradeoffs contributing to liver de-targeting and cell type specific tradeoffs that differ between myeloid and B cell targeted formulations. By navigating these quantitative trends, FALCON designed tailored compositions with different transfection profiles. While further *in vivo* studies are needed to verify these relationships and explain their mechanism, our findings show designing LNP formulations for selective delivery is not merely a matter of maximizing transfection but requires strategic navigation of competing design pressures, a concept that, while intuitive, has not been systematically quantified or incorporated into screening workflows^48^.

Through model-driven Pareto-optimization, FALCON enables quantitative evaluation of achievable delivery trade-offs without requiring a priori weighting of competing objectives. This is particularly valuable in delivery applications where it is often unclear what combinations of potency and selectivity are experimentally achievable within a given design space. Rather than imposing subjective weighting factors before the achievable landscape is known (for example, a scientist may assign selectivity twice the weight of potency yet have no prior knowledge of whether such a preference will result in a modest tradeoff or a severe one), FALCON first identifies the full set of optimal tradeoff solutions within the explored formulation space. Researchers can then apply application-specific criteria, constraints, or utility functions for the therapeutic context of interest to select formulations most appropriate for downstream validation. As quantitative relationships among competing delivery objectives and their relative importance become better established, future versions of the framework may benefit from incorporating validated utility functions directly into the optimization process.

There are several areas where FALCON could be improved. Currently, the platform focuses exclusively on compositional ratios and does not account for lipid chemical identities, particularly ionizable and helper lipids, well-known determinants of LNP performance^9,70^. While compositional changes can modulate potency and selectivity, composition alone presents a limited design space and may be insufficient to achieve substantial improvements across multiple delivery objectives. Accordingly, the current implementation of FALCON should be viewed as a framework for task-specific optimization within a defined formulation space, rather than a universally transferable model for LNP design. One potential future direction would be to incorporate lipid structural optimization engines such as AGILE and LiON, to enable joint exploration of lipid composition and chemical structures^36,39^. Such an extension would not be a direct continuation of the current composition-optimization framework but require the integration of a distinct modeling layer that incorporates molecular structure as an explicit design variable. Additionally feature engineering beyond the current implementation, including molecular representations, structure-based fingerprints, or chemical descriptors, as well as predictive models capable of effectively learning from these inputs (*e.g.*, graph neural networks) would be required. As such, this represents a distinct modeling framework from the composition-based surrogate models employed in the present study.

While our use of separately trained models for each objective proved effective, future implementations could leverage multi-task models to streamline the optimization process, enabling shared representations that capture both conserved and cell type-selective design features underlying LNP-mediated delivery^71,72^. Adopting more sophisticated strategies to direct formulation search, including batch and constrained optimization^33,34^, and third-generation genetic algorithms (*e.g.*, NSGA-III) to handle cases of many or tiered objectives^73^, could further enhance the effectiveness of FALCON’s design framework. Looking ahead, we envision expanding FALCON’s optimization objectives beyond transfection outcomes towards more holistic optimization frameworks. Integrating intermediate readouts of biological processes, such as hepatic clearance, cellular uptake, endosomal escape, or immunogenicity profiles, would enable multi-parametric design of delivery vehicles tailored for complex therapeutic applications^74^.

In summary, FALCON presents a framework for moving beyond empirical LNP screening for singular cell targets toward data-driven, multi-objective design for cell-type-selective delivery. By efficiently uncovering both high-performance and selectively targeted formulations, FALCON suggests that rational, interpretable design of gene delivery systems is achievable. As the field continues to explore increasingly complex delivery objectives, integrating computationally guided optimization strategies offers a promising path towards accelerating the development of precision nanomedicines.

## METHODS

### Materials

DLin-MC3-DMA was purchased from MedKoo Biosciences. SM-102 (BP-25499) was from BroadPharm. cKK-E12 (36700) was purchased from Cayman Chemical. ALC-0315, DSPC, 18PG (sodium salt), DOPC, DOPE, and DOTAP were from Avanti Polar Lipids. Cholesterol was from Sigma-Aldrich. DMG-PEG (MW 2000) (DMG-PEG2000) was from NOF America Corporation. Cell lines obtained from the American Type Culture Collection (ATCC) include Ramos B (human Burkitt’s lymphoma B cell line, ATCC CRL-1596), THP-1 (human monocytic leukemia cell line, ATCC TIB-202), Hep G2 (human hepatocellular cell line, ATCC HB-8065), NIH-3T3 (NIH/Swiss mouse embryonic fibroblast cell line, ATCC CRL-1658), and C2C12 (immortalized mouse myoblast cell line, ATCC CRL-1772). DC 2.4 cells were a gift from the lab of Prof. Jonathan Schneck at Johns Hopkins University School of Medicine, Department of Pathology. INS-1E cells were a gift from the lab of Prof. Dax Fu at Johns Hopkins University School of Medicine. Reporter lysis buffer and luciferin assay solution were purchased from Promega. All mRNA was purchased from TriLink BioTechnologies at a stock concentration of 1 mg/mL and capped using the TriLink CleanCap co-transcriptional capping method.

### FALCON Experimental Workflow

#### Preparation and Characterization of mRNA-LNPs

mRNA-LNPs were synthesized using the MANTIS liquid handler (Formulatrix) and Opentrons Flex lab robot. Dynamic light scattering (Nano ZS90 Zetasizer, Malvern Analytical) was used to measure the size (z-average), polydispersity index (PDI), and ζ-potential of synthesized LNPs. For the ζ-potential, LNPs were diluted 1:10 in PBS. Quant-it RiboGreen assay (ThermoFisher, R11490) was used to measure LNP encapsulation efficiency. The physicochemical properties of LNP formulations generated in this study are summarized in **Supplementary Tables 6 and 7**.

#### In vitro Transfection and Luciferase Assay

For *in vitro* library and iterative screening experiments, suspension cells (Ramos B and THP-1) were seeded at a density of 50,000 cells per well on the day of transfection. Adherent cells (HepG2, INS-1E, DC2.4, C2C12, NIH-3T3) were seeded at a cell density of 10,000 cells (HepG2, INS-1E) or 5,000 cells (DC2.4, C2C12, NIH-3T3) per well on the day before transfection. Cells were seeded in media containing 10% fetal bovine serum and 1% penicillin-streptomycin. Unless specified otherwise, firefly luciferase encoding mRNA-LNPs were dosed at the predetermined dosage level for each cell type (150, 25, 5, and 10 ng/well for Ramos B, THP-1 cells, HepG2, and INS-1E/DC2.4/C2C12/NIH-3T3, respectively). For the experiments described in Fig. 6, THP-1 and HepG2 cells were treated with mRNA-LNPs at 10 and 2.5 ng/well, respectively. After 24 h of incubation, cells were lysed, and luciferase expression was analyzed using a luminometer following a standard protocol.

### FALCON Computational Pipeline

#### Data Preprocessing

Prior to model training, the datasets generated were preprocessed in order to reduce experimental noise and equalize feature contributions. Raw luciferase signals were first transformed to our luciferase expression metric (LE), defined as the fold-change in treatment signal over no treatment signal, typically around 10^1.1^, as shown in **Equation 1**:

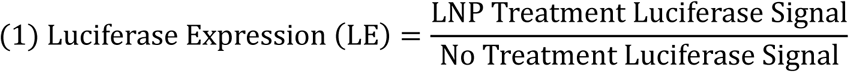

These values were then natural log-transformed to improve distribution symmetry. Both input features and output values were then scaled to a 0–1 range using min–max normalization. As datasets in this study were generated and analyzed in an iterative fashion, batch-wise normalization was performed using a consistent set of internal control LNP formulations to ensure comparability across experimental runs.

A total of six internal controls were used, selected to include both historically high-and low-performing formulations or established reference compositions. For each control formulation, the mean Ln(LE) across all batches was treated as the reference value (Ln(LE)^ref^). The difference between this reference and the measured value (Ln(LE)^batch^) in a given batch was computed for each control, and the average of these six differences was used as the batch-specific normalization factor. This factor was then applied uniformly to all formulations within that batch. The normalized Ln(nLE) of a given formulation j, denoted Ln(nLE)_j_, with control formulations i from 1 to 6, was calculated as shown in **Equation 2**:

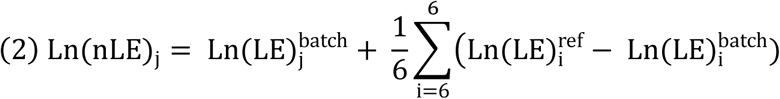

#### Model Training and Evaluation

Model training was performed as previously described in *Cheng et al.,* using established procedures for hyperparameter optimization and nested cross-validation from the Aspuru–Guzik group’s GitHub repository^13^. Briefly, the sci-kit learn (v1.2.2) and XGBoost (v1.7.1) Python packages were used to train XGBoost regression models on pre-processed input features, with performance evaluated via a 5-fold nested cross-validation strategy following an 85:15 train/test split. For each outer loop, an inner loop conducts a 4-fold split of the training data and runs 100 iterations of hyperparameter optimization for each split to minimize the validation mean absolute error (MAE) using a randomized search strategy. The final model was selected from the fold with the smallest validation-test MAE gap and retrained on the full training set. Final model performance was evaluated on the final 15% hold-out set using MAE, Pearson’s correlation coefficient *r*, and Spearman’s rank correlation *ρ*.

#### Closed-Loop Optimization and Sampling Strategy

Closed-loop optimization was performed to automate the generation of novel LNP compositions predicted to maximize transfection efficiency and cell type-selectivity. Trained XGBoost models were used as the objective function and guided the optimization of formulation composition parameters in normalized (0–1) space with expanded bounds. Alternatively, in cases where certain parameter values were known to be unfeasible, a hard cutoff for raw feature suggestion bounds was implemented. These configurations are described in **Supplementary Table 9.**

Two global optimization methods were implemented for single-objective compositional optimization: Bayesian Optimization (BO) and Dual Annealing (DA). BO was implemented using the bayes-opt package (v2.0.4), employing the Expected Improvement acquisition function with 10 random initialization points and 50 optimization iterations. Dual annealing was implemented using the scipy.optimize (v1.10.1) package, configured with a high initial temperature (20,000) and visit parameter of 2.8 to enable broad exploration. Each algorithm was iterated with greedy selection of the top-predicted formulation until a batch of distinct experimentally feasible LNP compositions was generated. In a preliminary test of the two iterative search strategies, both BO and DA identified and converged on similar composition regions, demonstrating comparable final performance (**Supplementary Fig. 48**). Subsequent studies used DA for single-objective tasks.

Multi-objective optimization was performed using the NSGA-II algorithm to identify Pareto-optimal formulations that balance on-target transfection and off-target de-targeting. Specifically, target-cell nLE was treated as an objective to maximize, while off-target cell nLE was treated as an objective to minimize. This configuration allows NSGA-II to identify Pareto-optimal trade-off solutions without requiring the a priori specification of a composite selectivity metric.

The search was implemented using the pymoo package (v0.6.1.3). Latin Hypercube Sampling was used for initial population generation. To maintain population diversity, adaptive genetic operators were introduced: Adaptive Simulated Binary Crossover (base probability = 0.9, η = 15) and Adaptive Polynomial Mutation (base probability = 0.1, η = 20). The algorithm was run for 100-500 generations starting from an initial population of 500-1000, with duplicate solutions eliminated. After completion of the algorithm, the Pareto front was extracted. During iterative optimization, a diversity-weighted greedy selection strategy was applied where points were chosen by maximizing Euclidean distance in objective space from all previously selected points, to select a subset of formulations diverse in objective space for experimental synthesis.

Following the final FALCON iteration, formulations on the final Pareto-optimal set were filtered using biologically motivated transfection thresholds. For the case studies explored in the main text, formulations were required to exhibit on-target Ln(nLE) > 2 and off-target Ln(nLE) values at least one unit lower than the reference formulation. The remaining formulations were then ranked according to their natural log transformed Selectivity Score, defined as the ratio of target-cell nLE to off-target-cell nLE (**Equation 3**):

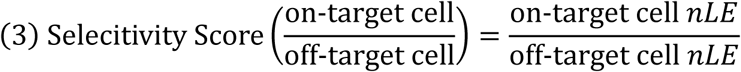

Following threshold-based filtering, formulations were prioritized for downstream validation using the Selectivity Score as a secondary ranking metric. The Selectivity Score was not used during optimization and served exclusively for post hoc candidate ranking and visualization of optimization progress.

When initial library screening was particularly sparse, an objective-driven search was complemented with I-optimal sampling at each iteration to reduce model uncertainty and improve coverage of the design space. I-optimal design was implemented by fitting a Gaussian Process surrogate to XGBoost predictions and iteratively selecting formulations that minimized the average predictive variance. In later studies, a minimum diversity threshold (*δ*) weighted by absolute SHAP values for each feature was explored (such that differences in highly important features were prioritized over variations in weaker features) to ensure that sampled formulations were diverse in experimentally meaningful dimensions of the design space. A candidate formulation x was accepted only if it was sufficiently different from the set of all previously selected formulations S, according to **Equation 4**:

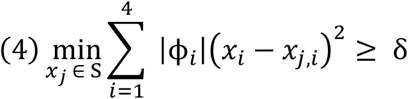

where *x_i_* and *x_j,i_* denote feature *i* values for the candidate and previously selected formulations, respectively, and |ϕ_*i*_| is the absolute SHAP importance of feature *i*. Search strategy configurations, including algorithms used, batch size, and search bounds for each study can be found in **Supplementary Table 9**.

### Post Hoc FALCON Interpretation and Analysis

#### PCA Visualization

PCA, a dimensionality reduction technique, was performed in R using the prcomp function from the base stats (v4.5.0) package. For each analyzed dataset, the two principal components capturing the greatest variance among all tested formulations were extracted and used to visualize the compositional search space in 2D plots with the ggplot2 (v3.5.2) package.

#### SHAP Feature Importance Analysis

Model interpretation was performed using the SHAP package (v0.42.1) to evaluate the contribution of each LNP compositional parameter to predicted transfection outcomes. SHAP beeswarm plots and dependence plots were used for global feature importance analysis, while waterfall plots were used to show per-feature contributions for selected individual formulations.

### Cell Transfection Validation Assays

#### Co-Culture Transfection Assays

For co-culture experiments, cells of interest were co-seeded into the same well at equal ratios. Cells were harvested for flow cytometry readouts 24h after LNP dosage.

#### Transfection of Primary Cells

Peripheral blood mononuclear cells (PBMCs) were isolated from a healthy donor leukopak (STEMCELL) by Ficoll-Paque PLUS (Cytiva) gradient centrifugation and frozen in CryoStor CS10 (BioLife Solutions). Primary human CD19^+^ B cells were isolated from healthy donor PBMCs by immunomagnetic positive selection (STEMCELL) and expanded *ex vivo* in serum-free human B cell expansion media (ImmunoCult, STEMCELL) for 3 days. Activated, expanded B cells were frozen at 10^7^ cells/mL in RPMI 1640 media (ATCC) supplemented with 10% fetal bovine serum (FBS), 10% DMSO, and stored in liquid nitrogen. B cells were thawed and plated in RPMI 1640 medium supplemented with 10% FBS and 1% penicillin-streptomycin for experiments.

### Ribogreen Assay

Encapsulation efficiency and encapsulated RNA concentration of LNPs were determined using Quant-it^TM^ RiboGreen (Thermo Fisher Scientific) and calculated using a standard curve of respective unencapsulated mRNA following standard protocol. Briefly, LNPs were incubated with Ribogreen reagent with or without 0.5% w/v Triton X-100, and the fluorescence intensity was measured to quantify free RNA content, where the measurement of LNPs without Triton X-100 represents “unencapsulated RNA” and LNPs with Trition X-100 represents “Total RNA”. The following equations (**Equations 5** and **6**) were used to calculate the encapsulated RNA concentration and the encapsulation efficiency.

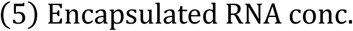

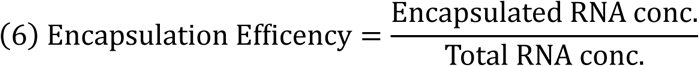

### Animals

All animal procedures were performed under an animal protocol approved by the Johns Hopkins Institutional Animal Care and Use Committee (protocol no. MO23E31). Male C57BL/6 mice (8–10 weeks old) were purchased from the Jackson Laboratory. Male Ai9 mice (8–10 weeks old) were bred in Johns Hopkins Animal Facilities and randomly grouped. The mice were supplied with free access to water and pelleted feed containing 5% fiber, 20% protein, and 5–10% fat. Mice typically provided 3–5 ml of water (150 mL/ kg body weight) and 4–5 g of pelleted feed (120 g/kg body weight) per day. The temperature of the mouse rooms was maintained at 18–26 °C (64–79 °F) with a relative humidity of 30–70%, and a minimum of 10 room air changes per hour. Standard shoebox cages with corncob as bedding were used to house the mice.

### *In vivo* LNP delivery for luciferase and Ai9 models

For the experiments in both C57BL/6 and Ai9 mice, LNP treatment was dosed intravenously retro-orbitally or intraperitoneally at either 10 µg per mouse (0.5 mg/kg) or 5 µg per mouse (0.25 mg/kg) as described in the main text. All LNPs were dialyzed and concentrated to their final injection concentration in 1x PBS. To minimize formulation-dependent variation in delivered mRNA, injection doses were normalized based on encapsulated mRNA concentrations determined by the RiboGreen assay. Mice were monitored for 15 min post-LNP injection for acute toxicity.

### Flow Cytometry

To validate cell-specific transfection profiles of candidate LNPs, cells were treated with GFP encoding mRNA-LNPs and incubated for 24 h at 37 °C. For Ramos B and THP-1 co-culture experiments, cells were first incubated with Fc receptor blocking solution (TruStain FcX™, BioLegend #422302) to reduce non-specific antibody binding, then washed and stained with BV421-anti-human CD20 (BioLegend #302330) and Zombie Yellow™ viability dye (BioLegend #423104). For Ai9 studies (**Figs. 2, 4, 5, and 6**), immune cell panels for spleen and liver were stained with BV421-anti-mouse CD11b (BioLegend #101236), BV605-anti-mouse CD45 (BioLegend #157217), BV785-anti-mouse NK1.1 (BioLegend #156539), AF488-anti-mouse CD20 (BioLegend #150432), APC-anti-mouse CD3 (BioLegend #100236), and Fixable NearIR Live/Dead (ThermoFisher #L10119). For non-immune and myeloid lineage liver cell staining, BV421-anti-mouse CD11b (BioLegend #101236), BV605-anti-mouse CD45 (BioLegend #157217), BV785-anti-mouse-CD326 (BioLegend #118245), APC-anti-mouse CD31 (BioLegend #102510), AF488-anti-mouse-F4/80 (BioLegend #123120), and Fixable NearIR Live/Dead (ThermoFisher #L10119). For Ai9 studies on pancreatic islet cells, cells were stained with BV421-anti-mouse-CD45 (BioLegend #147719) and Fixable Green Live/Dead (ThermoFisher #L34969). For OVA-specific T cell staining, APC-anti-mouse-CD3 (Biolegend #100236), APC-Cy7-anti-mouse-CD8a (Biolegend #100714), FITC-anti-mouse-CD45 (Biolegend #103108), BV421-OVA SIINFEKL tetramer (NIH tetramer core facility), and Zombie Yellow™ viability dye (BioLegend #423104). Staining steps followed the manufacturer’s protocols, and stains were diluted in PBS according to the manufacturer’s recommendations. Flow data were acquired using an Attune NXT flow cytometer and analyzed with FlowJo software.

### ELISA

Serum antibody responses were evaluated by measuring antigen-specific IgG, IgG1, and IgG2c levels with ELISA. Flat-bottom 96-well Nunc plates were coated with OVA protein at 2 μg per well in 100 mM carbonate buffer (pH 9.6) overnight at 4°C. Plates were then blocked for 2 h at room temperature using blocking buffer composed of 1% BSA in PBS. Mouse serum samples were first diluted 1:1000 in blocking buffer, followed by threefold serial dilution. After blocking, plates were washed with PBS-T consisting of PBS with 0.05% Tween-20. Diluted serum samples were added to the wells and incubated overnight at 4°C. HRP-conjugated goat anti-mouse IgG, IgG1, or IgG2c secondary antibodies (Southern Biotech Associates) were prepared in blocking buffer at dilutions of 1:2000, 1:4000, and 1:4000, respectively. Plates were incubated with the corresponding secondary antibody for 1 h at room temperature. After the 1 h incubation, plates were washed with PBS-T, TMB ELISA substrate solution was added and allowed to develop at room temperature for 30 min. The reaction was stopped by adding 50 μL of 4 N sulfuric acid to each well. Absorbance was measured at 450 nm using a plate reader. A sample was defined as positive when its absorbance was at least two times higher than that of the negative control.

## Statistical Analysis

Statistical analysis was performed using GraphPad Prism 8. Two-tailed Student’s *t*-tests were used for comparisons between two groups, and one-way analysis of variance (ANOVA) or two-way ANOVA followed by Tukey’s or Dunnett’s multiple comparisons test was used for comparisons involving more than two groups. To compare transfection efficiencies between two cell types across multiple formulations, multiple *t*-tests were conducted with Benjamini–Krieger–Yekutieli false discovery rate (FDR) correction (Q = 1%). Unless otherwise noted, all data are presented as mean ± standard error of the mean (SEM), performed in triplicate. A *p*-value of less than 0.05 was considered statistically significant.

## Acknowledgements

This study was partially supported by the National Institutes of Health under grant R01 CA293906-01A1 from the National Cancer Institute (to H.-Q.M. and R.K.) and grant R21 AI176764-01 from the National Institute of Allergy and Infectious Diseases (to M.F.K.).

## Author Contributions

W.H.T., L.C., and H.Q.M. conceptualized and designed this study. W.H.T., L.C., C.S., A.A., and R.K. developed and implemented the computational pipeline. W.H.T., L.C., D.Y., B.C., G.W., S.K., J.M., J. Lin, X.L., J.C., A.G., S.T., J. Liu, M.J., J.Y., M.P., G.S., I.C., D.A., X.H., and K.Z. performed the wet-lab experiments. W.H.T. and L.C. conducted data analysis. W.H.T., L.C., H.-Q.M., M.F.K., S.T., J.G., R.K., J.M., J. Liu, Y.Z., and D.Y. contributed to data interpretation and scientific discussion. H.-Q.M., M.F.K., and R.K. obtained funding and supervised this study. The manuscript was prepared by W.H.T., L.C., and H.-Q.M., with revisions and input from all authors. All authors approved the submitted manuscript version.

## Data Availability

The main data supporting the results of this study are available within the Article and its Supplementary Information. Raw and analyzed datasets generated during the study are available upon reasonable request from the corresponding author. Formatted screening datasets used for model training will be provided with this paper upon publication.

## Code Availability

The code used in this study for surrogate model training and optimization are publicly available under the MIT license at the GitHub repository (https://github.com/MaoResearchGroup/falcon-lnp-optimization).

