## Supplementary Material for "FALCON: Closed-Loop Multi-Objective Optimization of Lipid Nanoparticles for Cell-Selective mRNA Delivery"

### Contents

**Supplementary Table 1 |** Formulation composition feature ranges of initial grid-search library for FALCON single-objective optimization for B cell delivery.

**Supplementary Table 2 |** Formulation composition feature ranges of initial grid-search library for FALCON dual-objective optimization for selective B cell delivery against Hepatocytes.

**Supplementary Table 3 |** Formulation composition feature ranges of initial grid-search library for FALCON dual-objective optimization for selective monocyte delivery against hepatocytes.

**Supplementary Table 4 |** Formulation composition feature ranges of initial grid-search library for FALCON single-objective optimization for pancreatic beta cell delivery.

**Supplementary Table 5 |** Formulation composition and molar percentages of benchmark and FALCON-optimized LNPs described in the main text.

**Supplementary Table 6 |** Z-average size, polydispersity index, zeta potential, and encapsulation efficiency of reference composition-based LNPs described in the main text.

**Supplementary Table 7 |** Z-average size, polydispersity index, zeta potential, and encapsulation efficiency of FALCON-optimized LNPs described in the main text.

**Supplementary Table 8** | One-way ANOVA analysis followed by Tukey's multiple comparisons test and adjusted p-value corrected for multiple comparisons (6 within Ionizable lipid class, 6 between ionizable lipid classes) for **Figure 5b**.

**Supplementary Table 9 |** FALCON iterative search strategy modes and configurations.

**Supplementary Note 1 |** Stopping Criteria for Iterative Optimization.

**Supplementary Result 1 |** Application of FALCON for Pancreatic β Cell Optimization.

**Supplementary Result 2 |** Additional *In Vitro* FALCON Configurations.

**Supplementary Fig. 1 |** Initial dataset coverage, baseline model performance, and acquisition stability for representative Ramos B and THP-1 cell transfection prediction.

**Supplementary Fig. 2** | Process of robotic LNP assembly and screening.

**Supplementary Fig. 3** | Comparison of physicochemical attributes and transfection performance of robotic versus manually formulated LNPs.

**Supplementary Fig. 4 |** Boxplot of hold-out set error of panels of tested ML models.

**Supplementary Fig. 5** | Global hold-out performance of models at each stage of iterative FALCON optimization.

**Supplementary Fig. 6** | Illustration of search algorithms employed for *de novo* formulation search: dual annealing, Bayesian optimization, and NSGA-II.

**Supplementary Fig. 7** | Visualization of the convex hull volume encompassing the evaluated compositions of each search algorithm.

**Supplementary Fig. 8 |** Scatter plot of all tested LNPs during FALCON-driven compositional optimization for B cell transfection.

**Supplementary Fig. 9 |** PCA visualization of all tested compositions for FALCON-driven single-optimization for B cells, colored by nLE(Ramos).

**Supplementary Fig. 10 |** Example FALCON iterative optimization plot annotated with search behavior and stopping criteria used.

**Supplementary Fig. 11 |** Comparison of NSGA-II and I-optimal sampling performance across FALCON iterations.

**Supplementary Fig. 12 |** Gating strategy for the *in vitro* assessment of human primary B cell transfection via flow cytometry.

**Supplementary Fig. 13 |** Unguided search control for FALCON-driven single optimization for B cells.

**Supplementary Fig. 14 |** *Ex vivo* luminescence assay of liver after intravenous injection of MC3, DSPC-based Singel-Objective FALCON LNPs.

**Supplementary Fig. 15 |** Gating strategy for *in vivo* assessment of spleen immune compartment cell transfection in Ai9 mice via flow cytometry.

**Supplementary Fig. 16 |** Representative gating for tdTomato expression in CD45+CD20+ splenocytes.

**Supplementary Fig. 17 |** Gating strategy for *in vivo* assessment of liver non-immune and myeloid lineage cell transfection in Ai9 mice via flow cytometry.

**Supplementary Fig. 18 |** Gating strategy for *in vivo* assessment of liver immune compartment cell transfection in Ai9 mice via flow cytometry.

**Supplementary Fig. 19 |** Liver single cell transfection of FALCON B cell single-objective optimized LNP (F0162) in Ai9 mice.

**Supplementary Fig. 20** | Transfection correlation plot between Ramos B and HepG2 cells.

**Supplementary Fig. 21 |** Scatter plot of tested LNPs during FALCON-driven single-objective optimization of anionic LNPs for B cell transfection.

**Supplementary Fig. 22 |** PCA visualization of initial grid-search, single-objective (DA)- and dual-objective (NSGAII)-optimized compositions using FALCON-driven optimization of anionic LNPs.

**Supplementary Fig. 23 |** SHAP summary plots of models trained on anionic (18PG) or zwitterionic (DSPC) SM102 FALCON datasets for Ramos B and HepG2 transfection.

**Supplementary Fig. 24 |** *In vivo* Ai9 model transgene expression in spleen immune compartment and B cell selectivity.

**Supplementary Fig. 25 |** *In vivo* Ai9 model transgene expression in liver immune compartment and B cell selectivity.

**Supplementary Fig. 26 |** *Ex vivo* luminescence assay of spleen and liver after intravenous injection of F1179 and reference LNPs containing various IL and HL combinations.

**Supplementary Fig. 27 |** *Ex vivo* luminescence assay of spleen and liver after intravenous injection of SM102-based F1179 and reference LNPs containing different zwitterionic HLs.

**Supplementary Fig. 28 |** *In vivo* Ai9 model transgene expression in liver immune compartment cells comparing reference and F1179 LNPs with zwitterionic DSPC helper lipid.

**Supplementary Fig. 29 |** *In vivo* Ai9 model transgene expression in spleen immune compartment cells comparing reference and F1179 LNPs with zwitterionic DSPC helper lipid.

**Supplementary Fig. 30 |** *In vivo* Ai9 model transgene expression in liver non-immune and myeloid lineage cells comparing reference and F1179 LNPs with zwitterionic DSPC helper lipid.

**Supplementary Fig. 31 |** Gating strategy for *in vivo* assessment of splenic OVA-specific CD8+ T cells.

**Supplementary Fig. 32 |** Serum anti-OVA IgG titer collected from mice after vaccination with reference and F1179 LNPs.

**Supplementary Fig. 33** | Transfection correlation plot between THP-1 and HepG2 cells.

**Supplementary Fig. 34 |** *In vivo* Ai9 model transgene expression in liver immune-cell populations comparing F2080 with the reference LNPs.

**Supplementary Fig. 35 |** *In vivo* Ai9 model transgene expression in liver non-immune and myeloid cell populations comparing F2080 with the reference LNPs.

**Supplementary Fig. 36 |** *In vivo* Ai9 model transgene expression in spleen immune cell populations comparing F2080 with the reference LNPs.

**Supplementary Fig. 37 |** SHAP analysis of models trained on FALCON datasets for THP-1 and HepG2 transfection with SM102 and DSPC-based LNPs.

**Supplementary Fig. 38 |** SHAP analysis of models trained on FALCON datasets for Ramos and HepG2 transfection with SM102 and DSPC-based LNPs.

**Supplementary Fig. 39 |** Relative *In Vivo* biodistribution of Top FALCON and Reference LNPs.

**Supplementary Fig. 40 |** FALCON-guided single-objective compositional optimization of LNPs for delivery to pancreatic β cells.

**Supplementary Fig. 41 |** Gating strategy for *in vivo* assessment of pancreas cell transfection in Ai9 mice via flow cytometry after intraperitoneal administration of LNPs.

**Supplementary Fig. 42 |** Dual-objective FALCON-driven compositional optimization of LNPs for selective B cell and monocyte transfection.

**Supplementary Fig. 43 |** Gating strategy for co-culture assessment of LNP selectivity via flow cytometry.

**Supplementary Fig. 44 |** Dual-objective optimization of LNPs for selective monocyte transfection *in vitro*.

**Supplementary Fig. 45 |** SHAP feature importance analysis of Ramos B and THP-1 FALCON models.

**Supplementary Fig. 46** | Quadruple-objective optimization of LNPs in Ramos, C2C12, DC2.4, and NIH-3T3 cells.

**Supplementary Fig. 47** | Representative pairwise Pareto plots illustrating the FALCON quadruple-objective optimization process.

**Supplementary Fig. 48 |** Comparison of dual annealing and Bayesian optimization search algorithm performance.

**Supplementary Table 1 |** Formulation composition feature ranges of initial grid-search library for FALCON single-objective optimization for B cell delivery.

| Composition Parameters | Parameter Values |
|---|---|
| N/P ratio | 4, 8, 12 |
| (Dlin-MC3 + DSPC)% | 20, 50, 80 |
| DSPC/(Dlin-MC3 + DSPC)% | 0.49, 1.96, 50 |
| PEG/(PEG + Cholesterol)% | 0.20, 0.99, 9.09 |
| | Total: n = 81 |

**Supplementary Table 2 |** Formulation composition feature ranges of initial grid-search library for FALCON dual-objective optimization for selective B cell delivery against Hepatocytes.

| Composition Parameters | Parameter Values |
|---|---|
| N/P ratio | 4, 7, 10 |
| (SM-102 + 18PG)% | 17.9, 46.7, 77.8 |
| 18PG/(SM-102 + 18PG)% | 0.8, 7.7, 24.3, 42.9 |
| PEG/(PEG + Cholesterol)% | 0.20, 1, 5 |
| | Total: n = 108 |

**Supplementary Table 3 |** Formulation composition feature ranges of initial grid-search library for FALCON dual-objective optimization for selective monocyte delivery against hepatocytes.

| Composition Parameters | Parameter Values |
|---|---|
| N/P ratio | 4.2, 6.3 (for DSPC/(SM-102 + DSPC)% = 5)*;<br>5, 7.5 (for DSPC/(SM-102 + DSPC)% = 20)*;<br>6.7, 10 (for DSPC/(SM-102 + DSPC)% = 40)* |
| (SM-102 + DSPC)% | 40, 60, 80 |
| DSPC/(SM-102 + DSPC)% | 5, 20, 40 |
| PEG/(PEG + Cholesterol)% | 3.75, 5 |
| | Total: n = 36 |

*N/P ratio of the initial library was designed to hold total moles of SM102 constant across two SM102 only effective N/P ratios of 4 and 6. N/P ratios in the table above have been numerically adjusted to account for the number of nitrogens from both SM102 and DSPC, without changing the formulation itself.

**Supplementary Table 4 |** Formulation composition feature ranges of initial grid-search library for FALCON single-objective optimization for pancreatic beta cell delivery.

| Composition Parameters | Parameter Values |
|---|---|
| N/P ratio | 4, 7, 10 |
| (SM-102 + DOTAP)% | 20, 50, 80 |
| DOTAP/(SM-102 + DOTAP)% | 1, 10, 30, 50 |
| PEG/(PEG + Cholesterol)% | 0.2, 1, 5 |
| | Total: n = 108 |

**Supplementary Table 5 |** Formulation composition and molar percentages of benchmark and FALCON-optimized LNPs described in the main text.

| FALCON ID | N/P | IL% | HL% | Chol% | PEG% | (IL+HL)% | HL/(IL+HL)% | PEG/(PEG+Chol)% |
|---|---|---|---|---|---|---|---|---|
| Ref. | 6 | 50 | 10 | 38.5 | 1.5 | 60 | 16.7 | 3.8 |
| U0036 | 6.0 | 50.8 | 2.0 | 43.0 | 4.2 | 6.0 | 50.8 | 2.0 |
| F0059 | 8.0 | 25.0 | 25.0 | 49.9 | 0.1 | 50.0 | 50.0 | 0.2 |
| F0162 | 25.1 | 38.7 | 14.0 | 46.8 | 0.5 | 52.7 | 26.6 | 1.1 |
| F0225 | 5.0 | 23.0 | 29.4 | 44.0 | 3.6 | 52.4 | 56.1 | 7.6 |
| F0192 | 12.2 | 74.8 | 0.8 | 24.4 | 0.0 | 75.6 | 1.0 | 0.2 |
| F0178 | 21.4 | 60.0 | 6.0 | 33.9 | 0.1 | 66.0 | 9.1 | 0.4 |
| F1033 | 4.0 | 59.6 | 19.1 | 21.1 | 0.2 | 78.7 | 24.3 | 1.0 |
| F1119 | 4.1 | 29.8 | 3.5 | 65.8 | 0.9 | 33.3 | 10.5 | 1.3 |
| F1179 | 5.6 | 51.9 | 26.8 | 21.1 | 0.2 | 78.7 | 34.0 | 1.1 |
| F2080 | 8.8 | 82.0 | 5.6 | 11.7 | 0.7 | 87.6 | 6.4 | 5.9 |
| F2012 | 6.3 | 76.0 | 4.0 | 19.0 | 1.0 | 80.0 | 5.0 | 5.0 |
| F2124 | 13.9 | 83.9 | 2.7 | 12.9 | 0.6 | 86.5 | 3.1 | 4.2 |
| F2103 | 18.9 | 82.3 | 2.4 | 14.4 | 0.9 | 84.7 | 2.8 | 6.1 |
| F2083 | 14.7 | 73.1 | 7.0 | 18.7 | 1.2 | 80.1 | 8.8 | 6.1 |
| F3017 | 7 | 25.0 | 25.0 | 49.9 | 0.1 | 50.0 | 31.4 | 0.6 |
| F3118 | 24.3 | 34.3 | 15.7 | 49.6 | 0.3 | 61.3 | 37.2 | 4.8 |

**Supplementary Table 6 |** Z-average size, polydispersity index, zeta potential, and encapsulation efficiency of reference composition-based LNPs described in the main text.

| FALCON ID | Ionizable Lipid | Helper Lipid | Size (nm) | PDI | Zeta Potential (mV) | Encapsulation Efficiency (%) |
|---|---|---|---|---|---|---|
| Ref. | Dlin-MC3-DMA | DSPC | 145.5 ± 0.9 | 0.14 ± 0.01 | -6.0 ± 0.4 | 99.5% ± 0.4% |
| Ref. | SM-102 | DSPC | 121.5 ± 2.7 | 0.17 ± 0.05 | -3.7 ± 0.7 | 98.4% ± 0.5% |
| Ref. | SM-102 | 18PG | 164.2 ± 0.8 | 0.20 ± 0.03 | -5.9 ± 0.7 | 99.8% ± 0.2% |
| Ref. | SM-102 | DOPE | 182.6 ± 3.1 | 0.01 ± 0.01 | -5.7 ± 7.5 | 99.0% ± 0.4% |
| Ref. | SM-102 | DOPC | 165.7 ± 1.4 | 0.08 ± 0.02 | 3.5 ± 0.4 | 95.4% ± 0.4% |
| Ref. | ALC-0315 | DSPC | 188.8 ± 7.6 | 0.14 ± 0.04 | 0.5 ± 0.2 | 96.0% ± 0.4% |
| Ref. | cKK-e12 | DSPC | 192.4 ± 6.2 | 0.13 ± 0.08 | 0.0 ± 0.2 | 99.5% ± 0.4% |
| Ref. | ALC-0315 | 18PG | 174.1 ± 2.5 | 0.11 ± 0.02 | -15.3 ± 2.8 | 98.9% ± 0.2% |
| Ref. | cKK-e12 | 18PG | 184.3 ± 1.8 | 0.03 ± 0.01 | -5.4 ± 0.6 | 90.9% ± 0.3% |
| Ref. | SM-102 | DOTAP | 200.1 ± 9.4 | 0.16 ± 0.04 | 0.5 ± 0.1 | 96.1% ± 0.4% |

**Supplementary Table 7 |** Z-average size, polydispersity index, zeta potential, and encapsulation efficiency of FALCON-optimized LNPs described in the main text.

| FALCON ID | Ionizable Lipid | Helper Lipid | Size (nm) | PDI | Zeta Potential (mV) | Encapsulation Efficiency (%) |
|---|---|---|---|---|---|---|
| U0036 | Dlin-MC3-DMA | DSPC | 103.3 ± 2.2 | 0.13 ± 0.02 | -4.1 ± 3.3 | 99.8% ± 0.2% |
| F0059 | Dlin-MC3-DMA | DSPC | 257.1 ± 17.7 | 0.13 ± 0.06 | 1.6 ± 0.2 | 97.5% ± 0.0% |
| F0162 | Dlin-MC3-DMA | DSPC | 116.1 ± 2.1 | 0.21 ± 0.03 | 1.1 ± 0.3 | 96.1% ± 1.7% |
| F0225 | Dlin-MC3-DMA | DSPC | 115.5 ± 2.2 | 0.19 ± 0.03 | 0.0 ± 0.6 | 97.2% ± 5.2% |
| F0192 | Dlin-MC3-DMA | DSPC | 104.7 ± 2.4 | 0.20 ± 0.07 | -13.0 ± 1.3 | 78.2% ± 4.1% |
| F0178 | Dlin-MC3-DMA | DSPC | 92.5 ± 2.4 | 0.25 ± 0.02 | 0.0 ± 0.2 | 92.3% ± 2.3% |
| F1033 | SM-102 | 18PG | 371.9 ± 30.4 | 0.10 ± 0.04 | 1.0 ± 0.1 | 99.1% ± 0.2% |
| F1119 | SM-102 | 18PG | 236.7 ± 12.9 | 0.14 ± 0.02 | 0.1 ± 0.1 | 99.9% ± 0.0% |
| F1179 | SM-102 | 18PG | 392.8 ± 12.2 | 0.20 ± 0.06 | -12.5 ± 0.2 | 93.5% ± 1.0% |
| F1179 | ALC-0315 | DSPC | 189.9 ± 5.1 | 0.25 ± 0.02 | -7.4 ± 0.3 | 76.3% ± 1.2% |
| F1179 | cKK-e12 | DSPC | 309.7 ± 6.6 | 0.29 ± 0.02 | 4.6 ± 0.6 | 40.5% ± 3.2% |
| F1179 | Dlin-MC3-DMA | DSPC | 184.4 ± 1.0 | 0.19 ± 0.01 | 1.4 ± 0.4 | 86.6% ± 0.4% |
| F1179 | ALC-0315 | 18PG | 242.0 ± 5.8 | 0.17 ± 0.06 | -4.4 ± 2.5 | 99.2% ± 0.4% |
| F1179 | cKK-e12 | 18PG | 284.4 ± 10.3 | 0.13 ± 0.08 | -17.3 ± 0.9 | 53.0% ± 0.8% |
| F1179 | Dlin-MC3-DMA | 18PG | Visible flocculation, not tested further | | | |
| F1179 | SM-102 | DSPC | 196.2 ± 1.5 | 0.21 ± 0.02 | -3.3 ± 0.8 | 85.4% ± 1.2% |
| F1179 | SM-102 | DOPE | 362.3 ± 15.6 | 0.13 ± 0.13 | 5.4 ± 1.2 | 94.3% ± 0.5% |
| F1179 | SM-102 | DOPC | 166.9 ± 2.2 | 0.12 ± 0.02 | 13.1 ± 4.8 | 94.0% ± 0.3% |
| F2080 | SM-102 | DSPC | 167.4 ± 7.8 | 0.05 ± 0.04 | -2.2 ± 0.3 | 87.6% ± 0.5% |
| F2012 | SM-102 | DSPC | 151.8 ± 8.6 | 0.08 ± 0.09 | -5.9 ± 4.3 | 83.0% ± 1.5% |
| F3017 | SM-102 | DOTAP | 204.9 ± 8.7 | 0.20 ± 0.04 | 0.1 ± 0.2 | 97.4% ± 0.5% |
| F3118 | SM-102 | DOTAP | 260.0 ± 12.2 | 0.19 ± 0.03 | 0.0 ± 0.3 | 98.6% ± 0.5% |

**Supplementary Table 8 |** One-way ANOVA analysis followed by Tukey's multiple comparisons test and adjusted p-value corrected for multiple comparisons (6 within Ionizable lipid class, 6 between ionizable lipid classes) for **Figure 5b**.

| Comparison | SM102 | ALC0315 | cKK-E12 | MC3 |
|---|---|---|---|---|
| Ref (18PG) vs. F1179 (18PG) | 0.0006 | 0.0006 | 0.0006 | n/a |
| Ref (18PG) vs. Ref (DSPC) | 1.5174 | 0.2412 | 1.6152 | n/a |
| Ref (18PG) vs. F1179 (DSPC) | 1.692 | 0.2754 | 0.0006 | n/a |
| F1179 (18PG) vs. Ref (DSPC) | 0.0006 | 0.0006 | 0.0006 | n/a |
| F1179 (18PG) vs. F1179 (DSPC) | 0.0054 | 0.015 | 0.162 | n/a |
| Ref (DSPC) vs. F1179 (DSPC) | 0.0636 | 0.0048 | 0.0006 | 0.0072 |

**Supplementary Table 9 |** FALCON iterative search strategy modes and configurations.

| Experiment | Search Algorithm(s) | Objectives | Initial Library Size | Batch Size per Iteration | Batch Selection Strategy | Search Bounds |
|---|---|---|---|---|---|---|
| 0 (Fig. 2) | Dual Annealing | Single (↑Ramos) | 81 | 12 | Greedy | (0,3) in normalized space |
| 0 (Suppl) | Dual Annealing | Single (↑Ramos) | 81 | 12 | Greedy | (0,3) in normalized space |
| 0 (Suppl) | NSGA-II | Dual (↑Ramos, ↓THP-1) | 81 | 12 | Diversity-guided greedy Pareto selection | (0,1) in normalized space |
| 0 (Suppl) | NSGA-II | Dual (↑THP-1, ↓Ramos) | 81 | 12 | Diversity-guided greedy Pareto selection | (0,1) in normalized space) |
| 1 (Fig. 3) | NSGA-II | Dual (↑Ramos, ↓HepG2) | 108 | 15 | Diversity-guided greedy Pareto selection | (0,1.5) in normalized space |
| 1 (Fig. 3) | Dual Annealing | Single (↑Ramos) | 108 | 10 | Greedy | (0,3) in normalized space |
| 2 (Fig. 6) | NSGA-II, I-Optimal | Dual (↑THP-1, ↓HepG2) | 36 | 20 (10 per method) | Diversity-guided greedy Pareto selection with SHAP-weighted diversity threshold | Raw bounds: N/P ratio: [2,20] PEG/(Chol+PEG)%: [0.1,20] (IL+HL)%: [20,90], HL/(IL+HL)%: [0,80] |
| 3 (Suppl.) | Dual Annealing | Single (↑INS-1E ) | 108 | 5 | Greedy | (0,3) in normalized space |
| 4 (Suppl.) | NSGA-II | Quadruple (↑Ramos, ↓C2C12, ↓DC2.4, ↓NIH-3T3) | 108 | 12 | Diversity-guided greedy Pareto selection | (0,1.5) in normalized space |
| 4 (Suppl.) | Dual Annealing | Single (↑Ramos) | 108 | 12 | Greedy | (0,3) in normalized space |

### Supplementary Note 1 | Stopping Criteria for Iterative Optimization.

As FALCON's iterative optimization progresses, we expect to discover novel composition spaces with improved performance. For single-objective optimization tasks, this improvement is reflected in increases in the experimental objective metric (e.g., nLE). For multi-objective optimization tasks, although NSGA-II did not directly optimize a selectivity ratio, post hoc selectivity metrics were monitored as practical indicators of optimization progress. Thus, iterative optimization was continued as long as successive experimental rounds yielded meaningful improvements in the relevant performance metrics.

We terminated iterative optimization when additional rounds were unlikely to yield meaningful performance gains under the current surrogate and acquisition framework. This observation of convergence was defined by the following stopping criteria:

1. **Plateau in performance improvement**: When we began to observe that formulations suggested by FALCON were not resulting in significant performance gains, we took this as a sign that its search algorithm was concentrating sampling within locally optimal region(s) of the design space, where additional sampling yielded diminishing or minimal differences in performance gains. This was observed in iterations 3-5 of the annotated plot below.
2. **Algorithm has exploited the optimal region and is forced to explore marginally worse formulations.** Due to the exploitation–exploration balance inherent in FALCON's acquisition strategy, once high-performing regions were densely sampled, two outcomes were observed:
    - <u>Case 1</u>: The algorithm is fairly confident in an optimal region, and so instead elects for exploration of more uncertain regions in adjacent formulation spaces that have a low probability of yielding improved performance, and this can be reflected in an observed decrease in experimental performance. This behavior was observed in iteration 5 of the annotated plot below.
    - <u>Case 2</u>: The algorithm is very confident in its narrow optimal region, so it does not do any more exploration, and because of a minimal diversity threshold implemented to avoid repeated sampling of formulations that are not meaningfully distinct, the algorithm does not identify enough unique new formulations to sample in the optimal region and terminates. This was observed when we tried to generate a hypothetical 6$^{th}$ FALCON iteration for the annotated plot below.

**Supplementary Figure 10** is provided as an annotated version of **Figure 6b** as an example of the behaviors described above.

To assess whether the observed performance dip was attributable to diminishing returns within the explored design space rather than premature convergence of the optimization algorithm, we compared the performance trajectory of formulations selected by our primary objective-driven search algorithm (NSGA-II) with that of formulations selected using I-optimal sampling, an exploratory strategy designed primarily to improve predictive accuracy by sampling broadly across the design space.

If the objective-driven search had prematurely converged on a non-optimal region, we would expect I-optimal exploration to identify formulations with higher selectivity scores than those identified by NSGA-II, indicating the presence of superior regions of the search space that exploitative search had overlooked.

As shown in the accompanying analysis plot (**Supplementary Fig. 11**), this pattern was not observed. During the first exploration iteration, I-optimal sampling identified formulations with selectivity scores comparable to those identified by NSGA-II, suggesting that substantial opportunities for improvement still existed early in the optimization process. However, in subsequent iterations, NSGA-II consistently identified formulations with higher selectivity scores than those obtained through I-optimal exploration. From Iterations 2 to 4, the performance gap between NSGA-II-selected and I-optimal-selected formulations progressively increased. Even in the final iteration, I-optimal exploration did not identify formulations that exceeded the best-performing NSGA-II candidates from prior iterations.

This behavior suggests that NSGA-II identified a highly selective region of the explored formulation space and that additional sampling yielded diminishing performance gains. Notably, this occurred despite I-optimal sampling being specifically designed to explore under-sampled regions of the formulation space. While these analyses cannot exclude the existence of superior solutions elsewhere in the broader design space, they do not support progressive model overfitting or premature convergence as the primary explanations for the observed performance dip.

### Supplementary Result 1 | Application of FALCON for Pancreatic β Cell Optimization.

To demonstrate FALCON's applicability outside of immune cell delivery, we conducted a preliminary FALCON optimization of LNP delivery into pancreatic β cells relevant to Type 1 diabetes gene therapy. Recent work has shown that intraperitoneal administration of cationic helper lipid (DOTAP)-based LNPs improves selective gene delivery to the pancreas[56]. Building on this work, we conducted single-objective optimization of 4-component DOTAP-containing LNPs in INS-1E rat β cells (**Supplementary Fig. 40**). After 3 rounds of iterative screening (5 suggested LNPs per round, 15 FALCON suggested LNPs total), an optimized F3118 LNP was discovered outside of the initial library space that achieved more than 3-fold higher transfection in the isolated pancreatic islet CD45- cells in mice to compared to the reference (SM102, DSPC) and top initial library formulation (SM102, DOTAP), but no transfection differences in the CD45+ cells (**Supplementary Figs. 40** and **41**). Global SHAP analysis found that β cell transfection was associated with an intermediate (IL+HL)% content (~50% of all lipids) and a high N/P ratio (24.3) (**Supplementary Fig. 40)**. Recognizing the potential toxicity associated with higher NP ratios, future applications of FALCON could incorporate toxicity-related endpoints as additional objectives to identify formulations that balance delivery and safety.

### Supplementary Result 2 | Additional *In Vitro* FALCON Configurations.

In two final FALCON configurations, we performed 1) dual-objective optimization for selective transfection of Ramos B and THP-1 cells against each other in order to investigate compositional features distinguishing B cell- and monocyte-selective transfection (**Supplementary Figs. 42–45**), and also deployed FALCON for 2) quadruple-objective optimization to simultaneously minimize transfection in 3 off target cell types, demonstrating the feasibility of extending FALCON-guided formulation search to the many-objective case (**Supplementary Figs. 46** and **47**). In the first setup, FALCON-driven dual-objective optimization of B cells against monocytes identified LNPs with substantial B cell selectivity (~76.8% of transfected cells) in a co-culture of the two cell types, thus validating the utility of the model-guided optimization process (**Supplementary Figs. 42** and **43**). THP-1 single-objective potent and dual-objective selective FALCON LNPs both demonstrated enhanced monocyte transfection with minimal B cell transfection *in vitro* (**Supplementary Fig. 44**). These optimization experiments were performed using the same initial library and were included in the same dataset as the experiments reported in **Figure 2**.

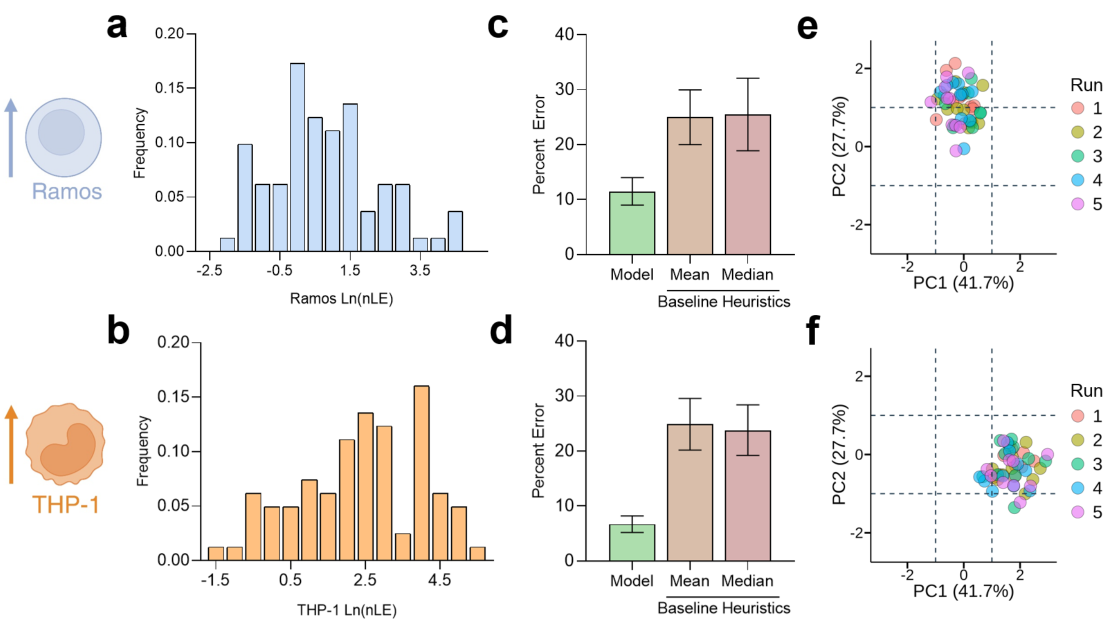


**Supplementary Fig. 1 | Initial dataset coverage, baseline model performance, and acquisition stability for representative Ramos B and THP-1 cell transfection prediction. a, b.** Histogram distributions of normalized Ln(LE) values for Ramos B cells (**a**) and THP-1 monocytes (**b**) in the initial base library show a range of transfection values across tested compositions. Ln(LE) represents natural log-transformed raw luciferase readings. **c,d.** Comparison of the initial trained model performance with baseline heuristics. The mean absolute percent error (MAE) from hold-out validation is lower than that of predictions based on the mean or median transfection values for both the Ramos (**c**) and THP-1 (**d**) models. **e,f.** PCA plots for the top 10 suggested LNP compositions from 5 independent runs reveal tight clustering across runs, indicating high acquisition stability and robustness of the model-guided search process for both Ramos (**e**) and THP-1 (**f**). The schematics in this figure were created with BioRender and released under a Creative Commons Attribution-NonCommercial-NoDerivs 4.0 International license.

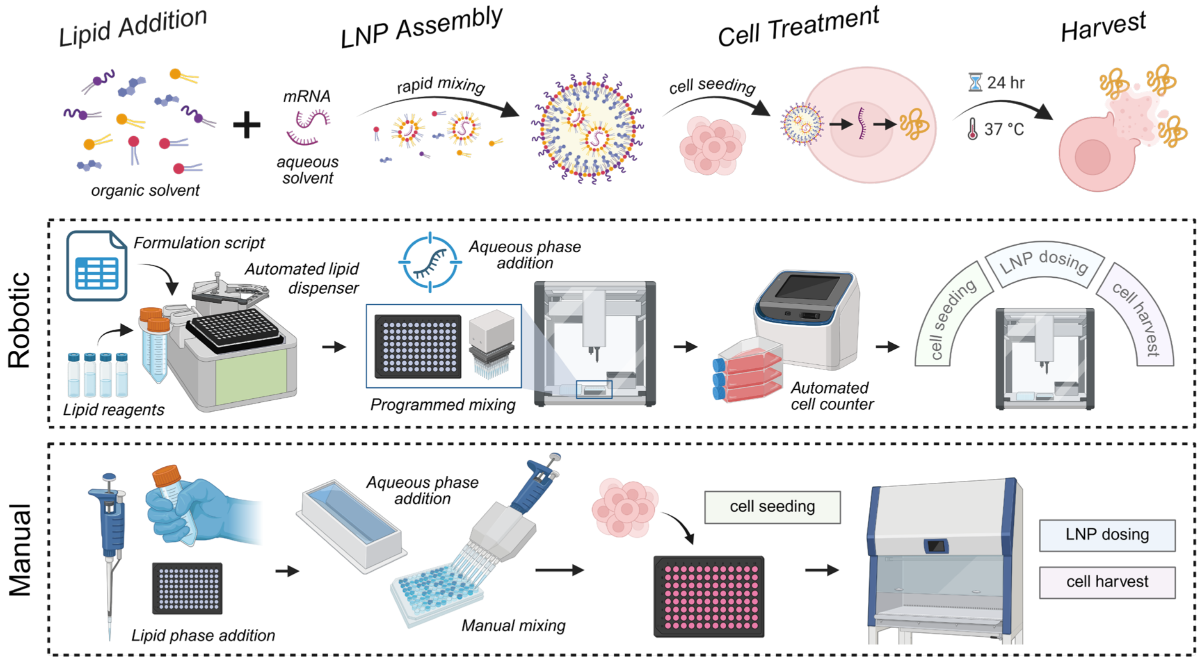


**Supplementary Fig. 2 | Process of robotic LNP assembly and screening.** In the robotic workflow, devices are utilized to assist each step of a manually performed transfection screening assay, from performing lipid component additions in a 96-well plate and self-assembly of LNPs through programmed mixing of aqueous and organic phase, to seeding, dosing, and harvesting transfected cell contents using a liquid dispenser equipped with HEPA/UV module. This figure was created with BioRender and released under a Creative Commons Attribution-NonCommercial-NoDerivs 4.0 International license.

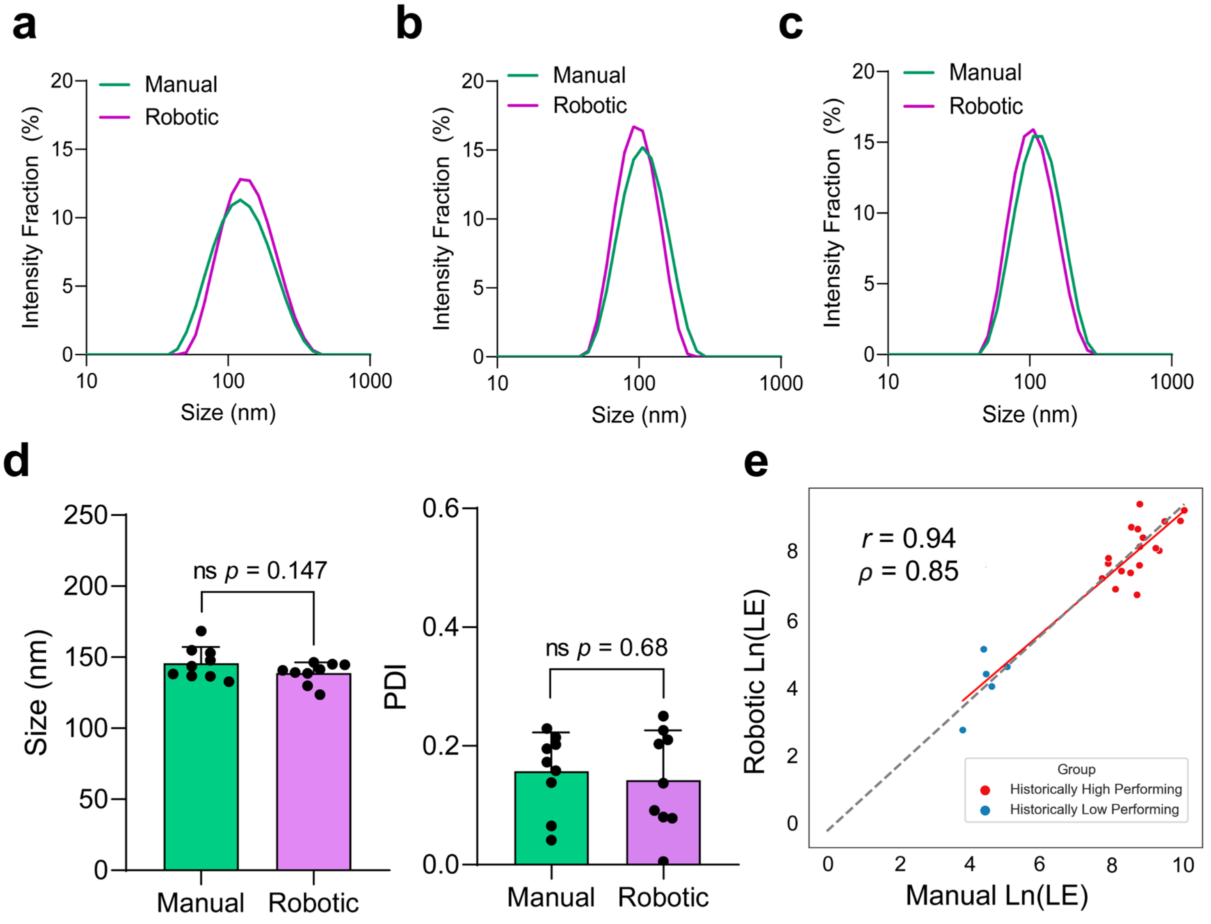


**Supplementary Fig. 3 | Comparison of physicochemical attributes and transfection performance of robotic versus manually formulated LNPs. a–c.** Representative Z-average diameter distributions measured by Dynamic Light Scattering (DLS) of selected top FALCON formulations assembled via traditional manual versus robotic methods. **d.** Z-average size and polydispersity index (PDI) measurements of a control LNP sampled at 9 random locations in formulated 96-well plates. **e.** Correlation between the transfection performance of robotic- and manually-formulated LNPs measured via luciferase reporter expression. Data are presented as mean ± SEM from a representative experiment (n = 9). The *p*-values were determined via a two-tailed unpaired Student's *t*-test. ns, $p > 0.05$.

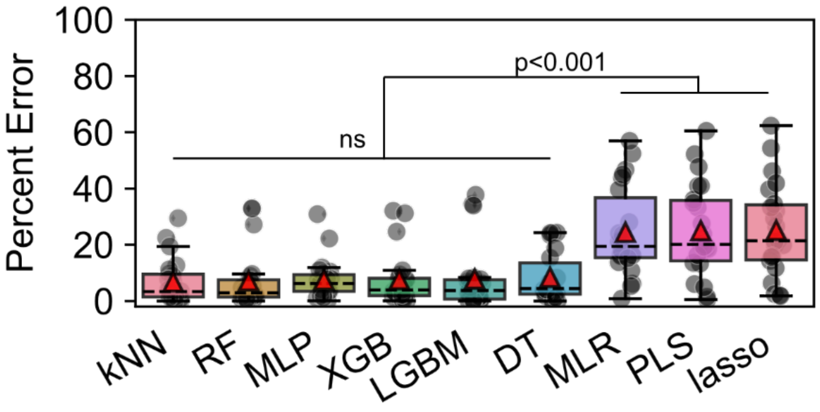


**Supplementary Fig. 4 | Boxplot of hold-out set error of panels of tested ML models.** Boxes centered around the MAE (red triangles), and dots represent individual hold-out set absolute error. Models are ranked from lowest (left) to highest (right) MAE (n = 129).

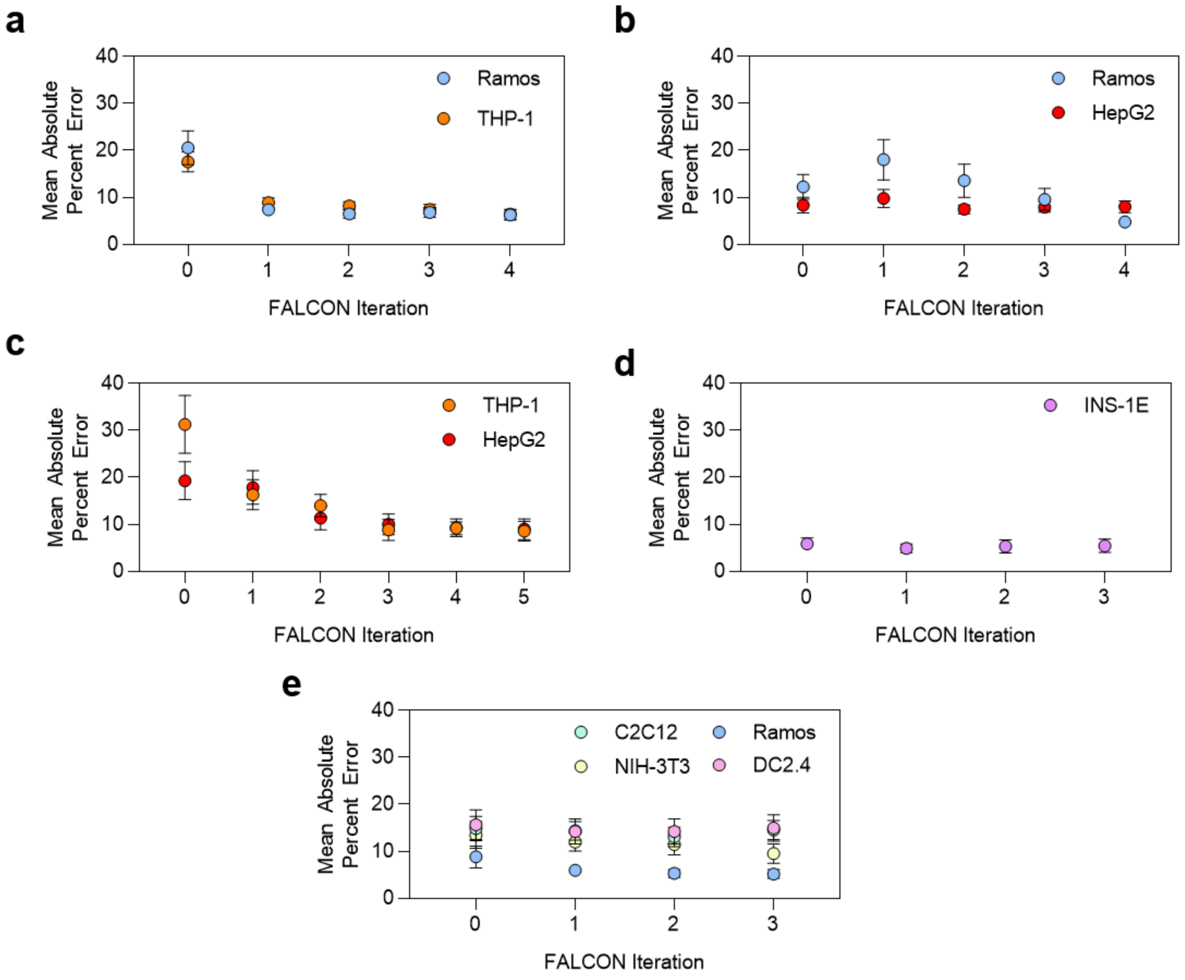


**Supplementary Fig. 5 | Global hold-out performance of models at each stage of iterative FALCON optimization.** XGBoost models were trained on various data subsets and evaluated on a fixed global hold-out set (15% of the total dataset) generated via random sampling of the final FALCON dataset for each experiment, stratified on the transfection output distribution, allowing cross-comparison of the trained model's generalizability across the full design space explored by FALCON. Mean absolute percent error metrics are shown for FALCON Ramos B single- and dual-objective optimization against THP-1 **(a)**, Ramos single- and dual- objective optimization against HepG2 **(b)**, THP-1 dual-objective optimization against HepG2 **(c)**, INS-1E single-objective optimization **(d),** and Ramos against DC24, C2C12, and NIH-3T3 quadruple-objective optimization **(e)** experiments. Plots demonstrate generally stable or improving predictive ability when retraining models across iterations. Error bars on plot denote SEM.

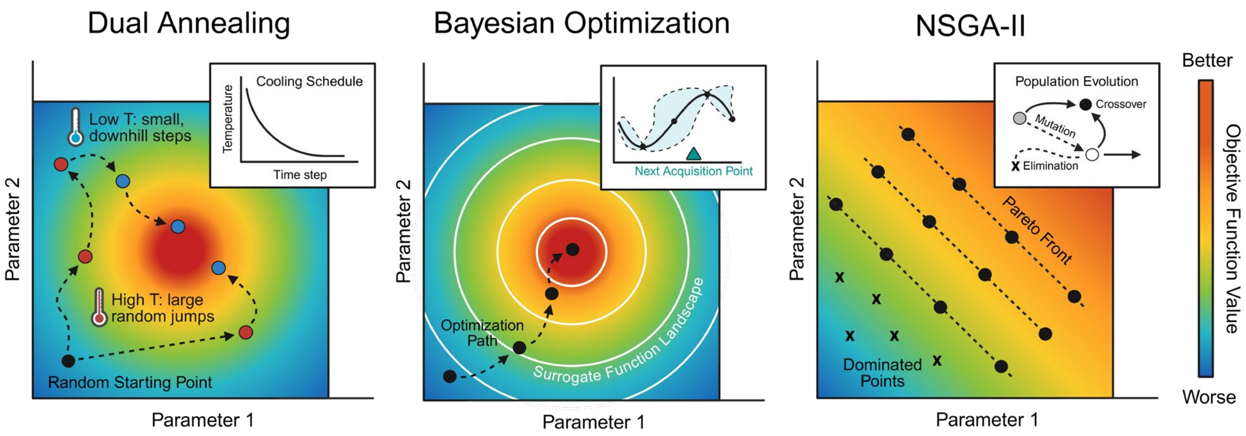


**Supplementary Fig. 6 | Illustration of search algorithms employed for *de novo* formulation search: dual annealing, Bayesian optimization, and NSGA-II.** Dual annealing is a stochastic algorithm that employs a thermodynamically inspired cooling schedule to balance random global search (at high temperature) and local refinement (at low temperature). Bayesian optimization leverages a surrogate model (concentric white rings) to sample points according to an acquisition function that balances predicted performance and uncertainty. NSGA-II performs multi-objective optimization by evolving a population of candidate solutions over multiple generations, through simulated mutation and crossover operations, and preserving non-dominated solutions that approximate the Pareto front. Objective function values for each plot are represented by color gradients. This figure was created with BioRender and released under a Creative Commons Attribution-NonCommercial-NoDerivs 4.0 International license.

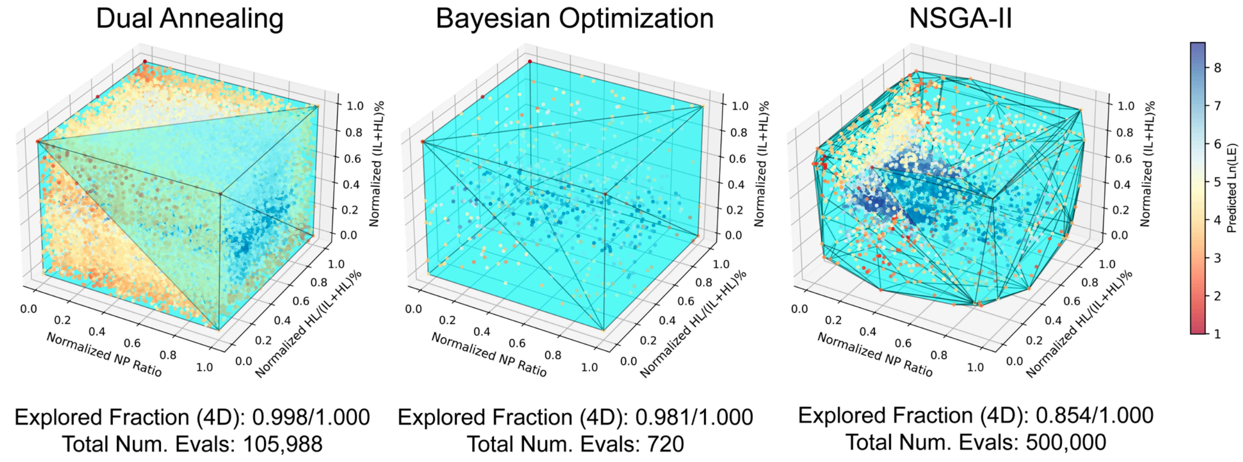


**Supplementary Fig. 7 | Visualization of the convex hull volume encompassing the evaluated compositions of each search algorithm.** 3D scatterplots of normalized composition space show the distribution of all the LNP formulations evaluated by each search algorithm during a representative in-silico optimization. Cyan mesh outlines the convex hull constructed from the evaluated points, representing the fraction of the input space explored by each algorithm. Points are colored according to predicted Ln(LE). The explored fraction (computed based on 4D convex hull volume) and the total number of evaluations for each algorithm is displayed below each plot. Dual annealing and Bayesian optimization single-objective optimization algorithms both achieved high explored fractions (0.998 and 0.981, respectively), indicating a rigorous breadth of search. Dual annealing demonstrates more uniform volumetric sampling through a greater number of evaluations, while Bayesian optimization is more efficient. NSGA-II multi-objective search algorithms also demonstrated a moderately high explored fraction (0.854), while focusing its evaluations on potential selective regions in the composition space.

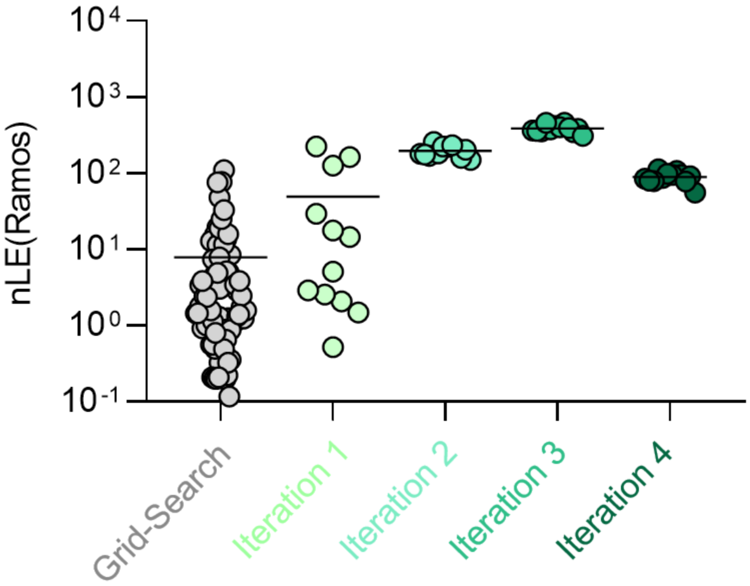


**Supplementary Fig. 8 | Scatter plot of all tested LNPs during FALCON-driven compositional optimization for B cell transfection.** 81 formulations were experimentally tested in the initial grid-search library, followed by 12 formulations per iteration, for a total of 129 tested LNPs. nLE (normalized luciferase expression) refers to raw readings from *in vitro* luciferase transfection experiments that were blank-subtracted and batch normalized against a set of internal controls.

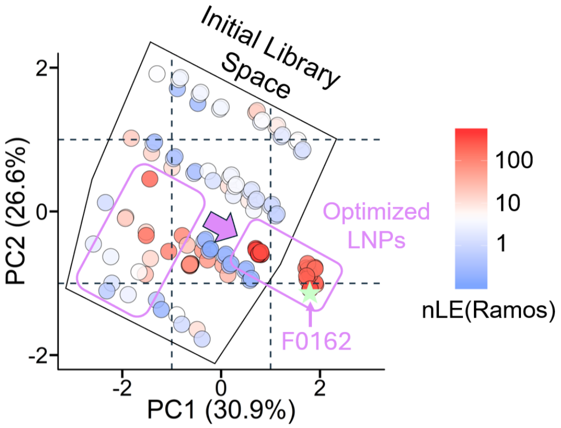


**Supplementary Fig. 9 | PCA visualization of all tested compositions for FALCON-driven single-optimization for B cells, colored by nLE(Ramos).** The search spaces encompassed by the initial grid-search library and by FALCON optimized LNPs are labelled. Arrow illustrates the progression of FALCON optimization beyond the initial library space. The green star indicates F0162, the optimized FALCON LNP nominated for further validation. nLE values are displayed on a logarithmic scale to allow visualization of the full dataset distribution.

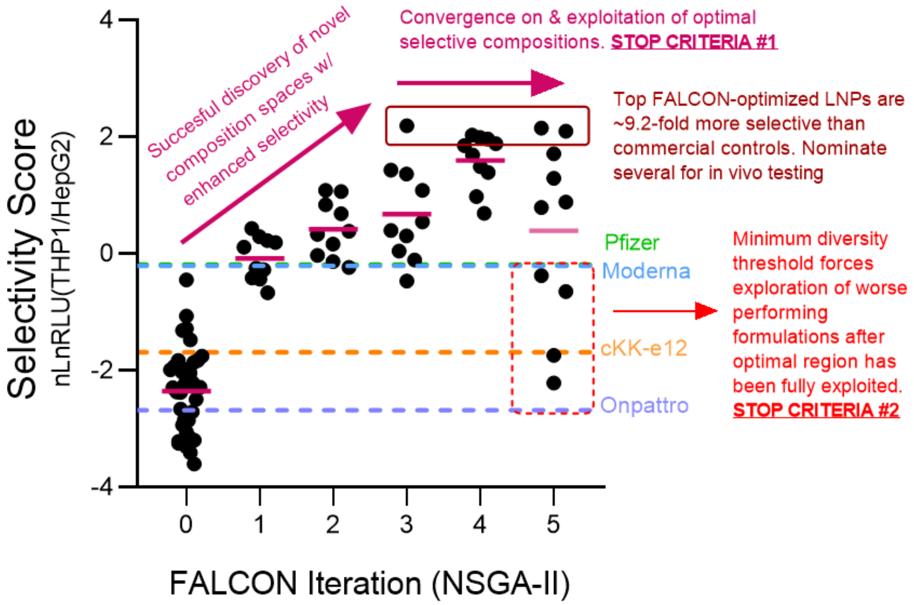


**Supplementary Fig. 10 | Example FALCON iterative optimization plot annotated with search behavior and stopping criteria used.**

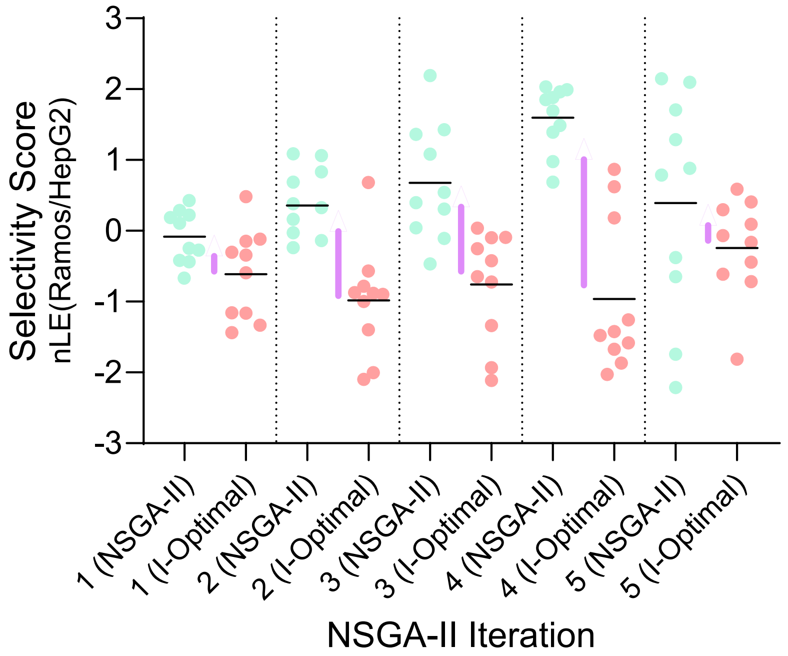


**Supplementary Fig. 11 | Comparison of NSGA-II and I-optimal sampling performance across FALCON iterations.**

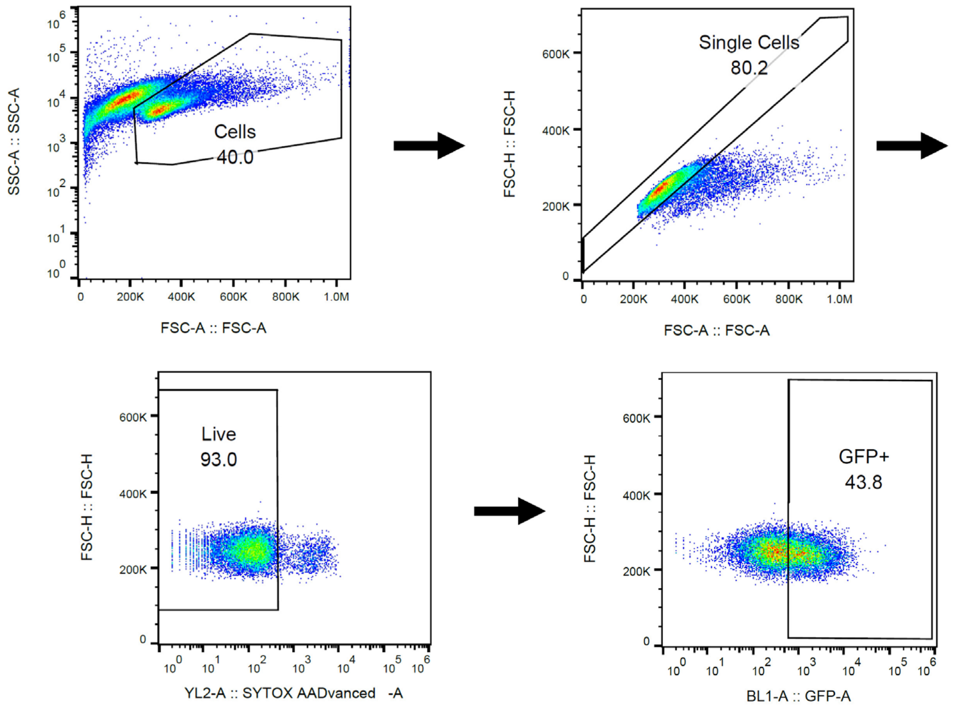


**Supplementary Fig. 12 | Gating strategy for the *in vitro* assessment of human primary B cell transfection via flow cytometry.** Representative flow plots show sequential gating: cells by size (FSC-A) and granularity (SSC-A), singlets based on FSC-H/FSC-A parameters, live cells using 7-AAD exclusion, and GFP+ cells within the live population to assess transfection efficiency.

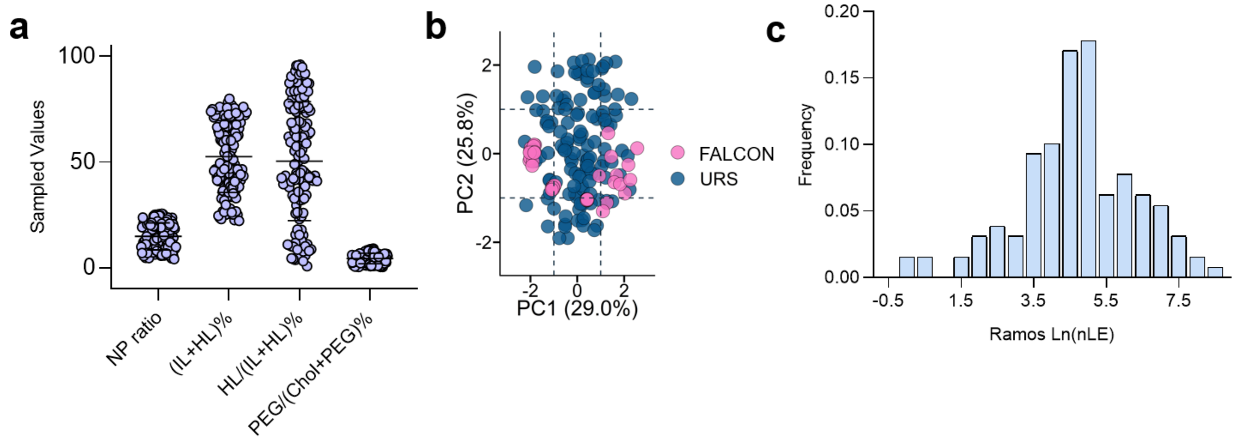


**Supplementary Fig. 13 | Unguided search control for FALCON-driven single optimization for B cells.** 129 formulations were sampled via uniform random sampling (URS) across the final parameter space covered by FALCON optimization. **a.** Scatterplot showing the sampled values and bounds for each feature parameter. **b.** PCA plot showing the composition of the URS dataset projected onto PCA axes alongside FALCON-suggested formulations. FALCON model-guided search is more targeted, whereas URS search is broader. **c.** Histogram distribution of transfection values (Ln(nLE)) obtained from *in vitro* screening in Ramos B cells. nLE values are displayed on a logarithmic scale, to allow visualization of the full dataset distribution, and are normalized within each experimental study.

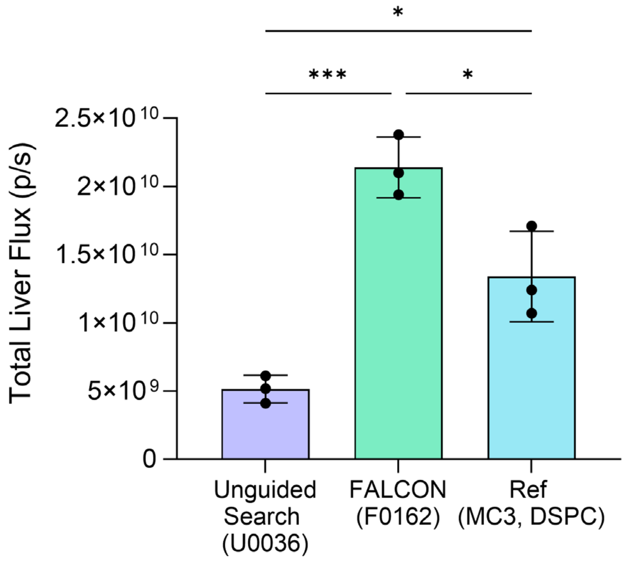


**Supplementary Fig. 14 | *Ex vivo* luminescence assay of liver after intravenous injection of MC3, DSPC-based Single-Objective FALCON LNPs.** Quantification of ex vivo luminescence flux in liver 6 h after intravenous injection of mFluc encapsulated LNPs.

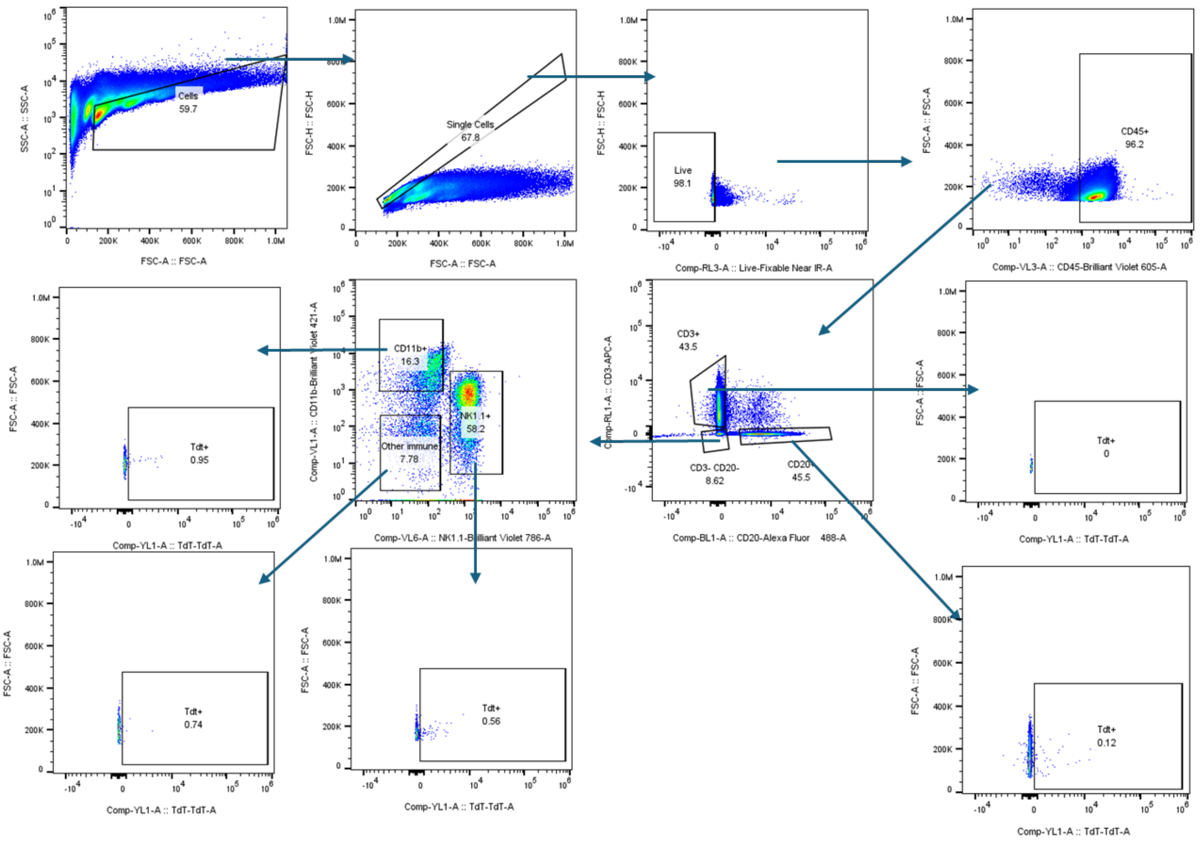


**Supplementary Fig. 15 | Gating strategy for *in vivo* assessment of spleen immune compartment cell transfection in Ai9 mice via flow cytometry.** Representative flow plots show sequential gating: cells by size (FSC-A) and granularity (SSC-A), singlets based on FSC-H/FSC-A parameters, and live cells using fixable live/dead exclusion. CD45+CD20+CD3- for B cells, CD45+CD3+ for T cells, CD45+CD3-CD20-CD11b+ for myeloid lineage, CD45+CD3-CD20-NK1.1+ for NK cells, and CD45+CD3-CD20-NK1.1-CD11b- for all other immune cells. The tdTom+ (Tdt+) value was used to assess successful transgene expression, and representative gating on PBS-treated mice is shown.

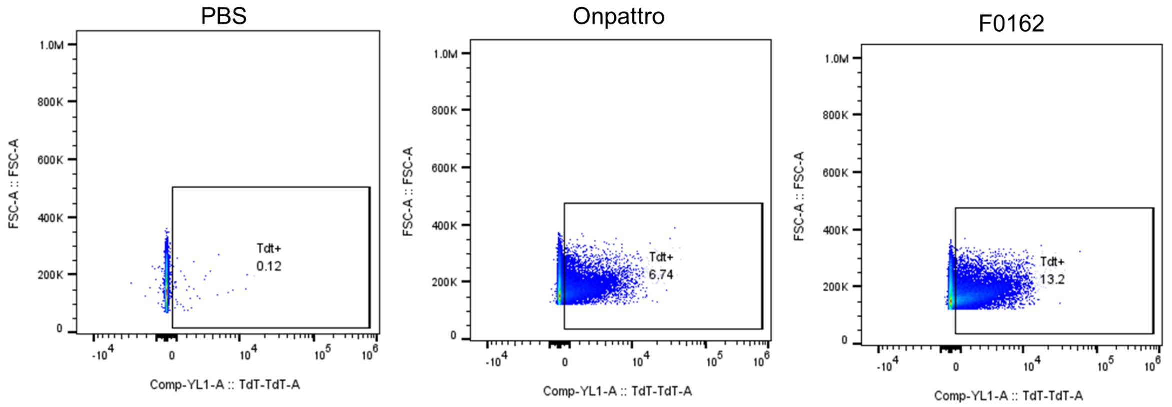


**Supplementary Fig. 16 | Representative gating for tdTomato expression in CD45+CD20+ splenocytes.** Representative flow plots showing tdTom+ (Tdt+) gating for PBS, Onpattro Ref (MC3, DSPC), and F0162 (MC3, DSPC) LNP-treated Ai9 mice.

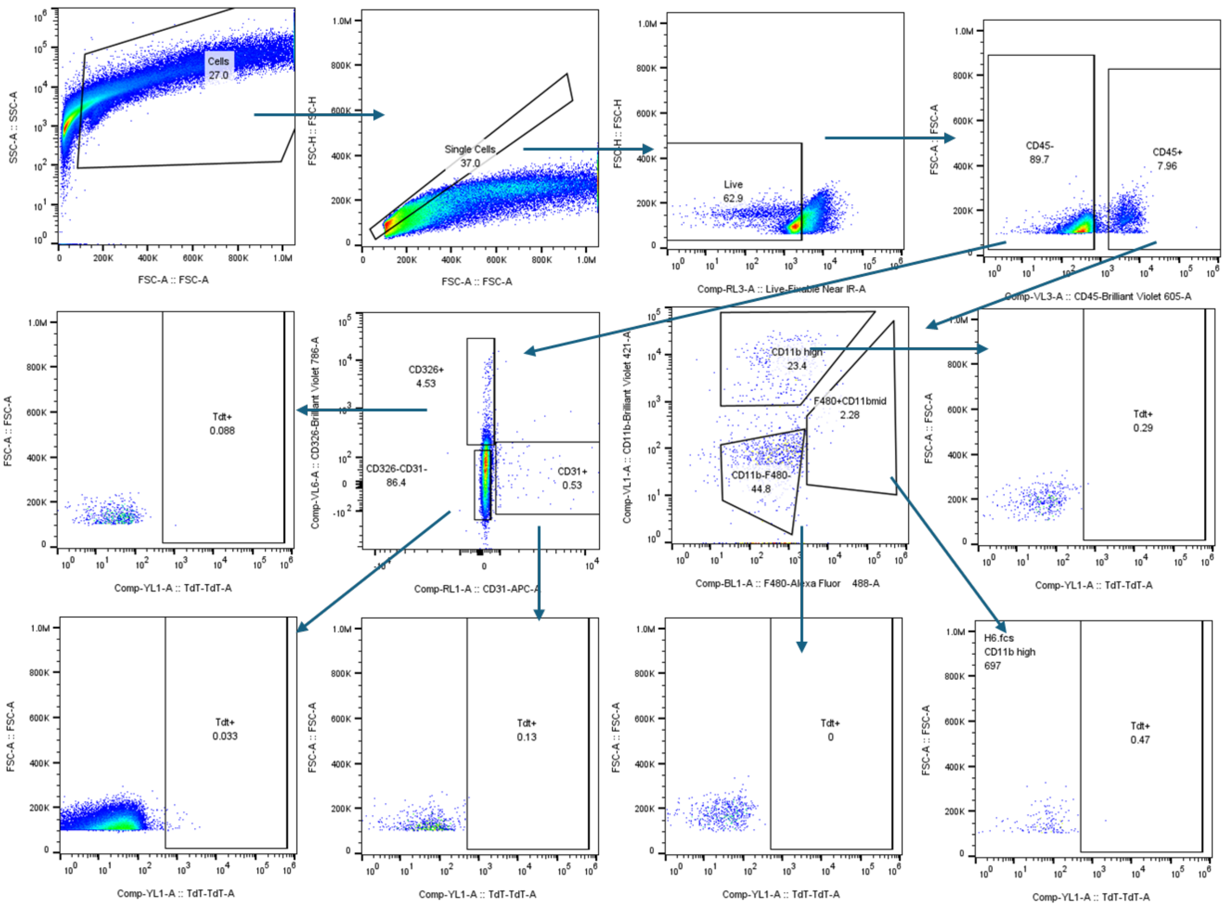


**Supplementary Fig. 17 | Gating strategy for *in vivo* assessment of liver non-immune and myeloid lineage cell transfection in Ai9 mice via flow cytometry.** Representative flow plots show sequential gating: cells by size (FSC-A) and granularity (SSC-A), singlets based on FSC-H/FSC-A parameters, and live cells using fixable live/dead exclusion. For the non-immune compartment, CD45-CD31+CD326- for endothelial cells, CD45-CD31-CD326+ for epithelial cells, and CD45-CD31-CD326- cell population, likely made up predominantly by liver hepatocytes. For the immune compartment, CD45+CD11b+ for myeloid lineage, CD45+CD11b$^{mid}$F4/80+ for Kupffer cells, and CD45-CD11b-F4/80- for other immune cells. The TdTom+ (Tdt+) value was used to assess successful transgene expression, and representative gating on PBS-treated mice is shown.

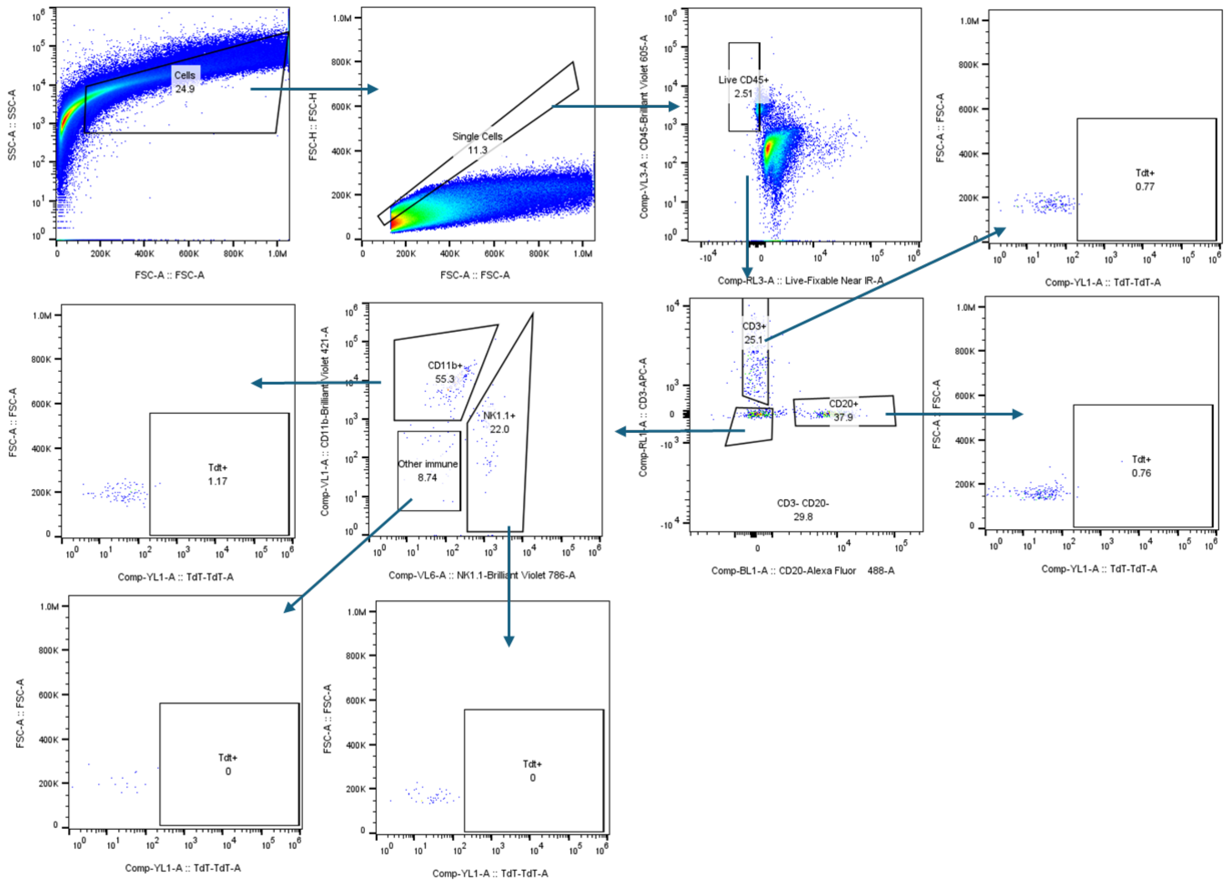


**Supplementary Fig. 18 | Gating strategy for *in vivo* assessment of liver immune compartment cell transfection in Ai9 mice via flow cytometry.** Representative flow plots show sequential gating: cells by size (FSC-A) and granularity (SSC-A), singlets based on FSC-H/FSC-A parameters, and live cells using fixable live/dead exclusion. CD45+CD20+CD3- for B cells, CD45+CD3+ for T cells, CD45+CD3-CD20-CD11b+ for myeloid lineage, CD45+CD3-CD20-NK1.1+ for NK cells, and CD45+CD3-CD20-NK1.1-CD11b- for all other immune cells. The TdTom+ (Tdt+) value was used to assess successful transgene expression, and representative gating on PBS-treated mice is shown.

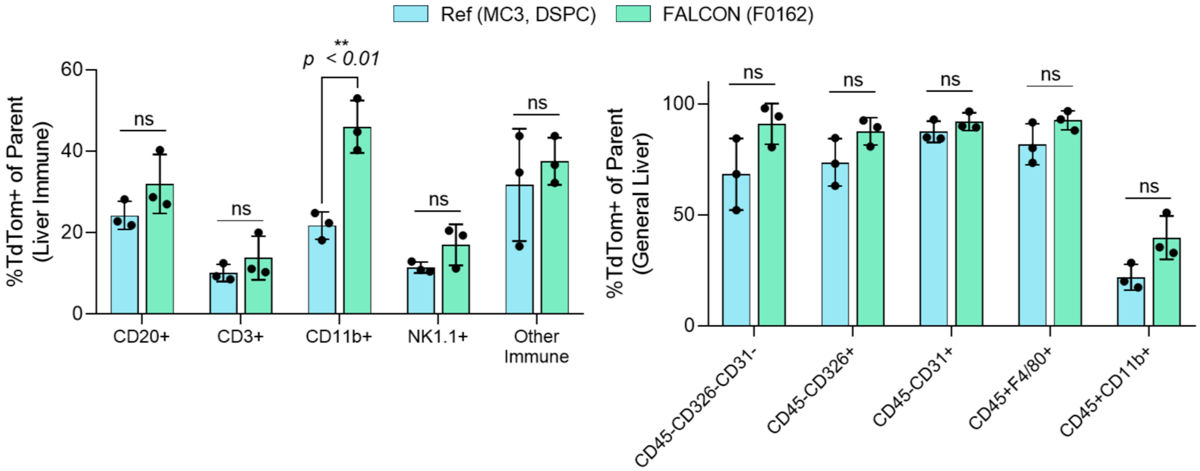


**Supplementary Fig. 19 | Liver single cell transfection of FALCON B cell single-objective optimized LNP (F0162) in Ai9 mice.** Liver single-cell suspensions from the same experiment were analyzed using two independent gating strategies: a liver immune panel (left) and a general liver (hepatocytes and Kupffer cells) panel (right). Data show the percentage of TdTomato$^{+}$ cells within each indicated population. Bars represent mean ± SEM (n = 3). Statistical comparisons were performed via multiple *t*-tests. ns, not significant, $^{**}p < 0.01$.

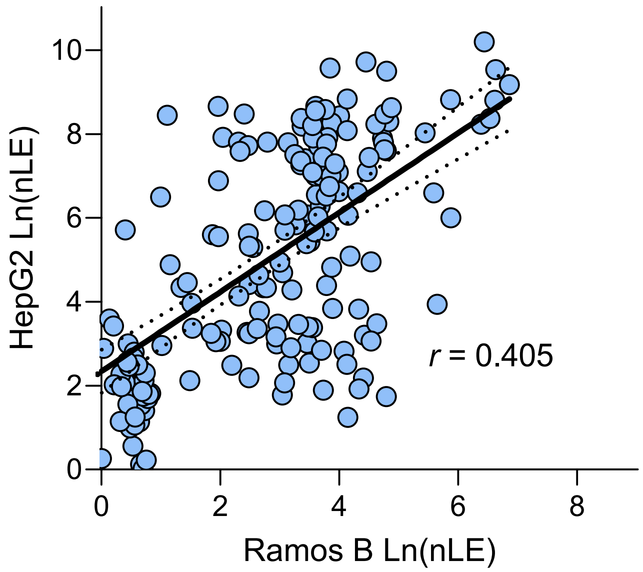


**Supplementary Fig. 20 | Transfection correlation plot between Ramos B and HepG2 cells.** Moderate positive correlation ($r$ = 0.405) highlights the complexity of isolating B cell-specific formulations and the need for multi-objective search algorithms.

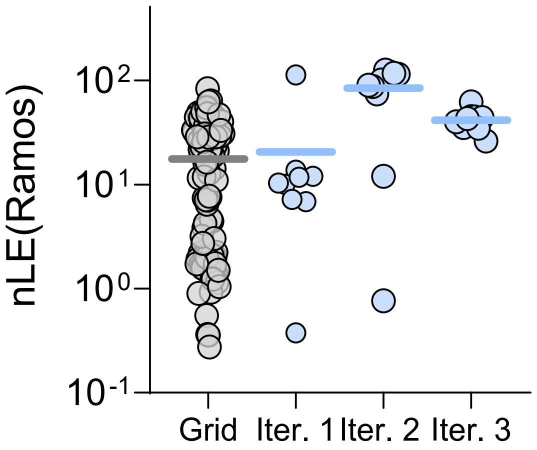


**Supplementary Fig. 21 | Scatter plot of tested LNPs during FALCON-driven single-objective optimization of anionic LNPs for B cell transfection.** 108 formulations were experimentally tested in the initial grid-search library, followed by 10 formulations per iteration, for a total of 137 tested LNPs (1 outlier formulation excluded from iteration 1). nLE (normalized luciferase expression) refers to raw readings from *in vitro* luciferase transfection experiments, that were blank-subtracted and batch normalized against a set of internal controls.

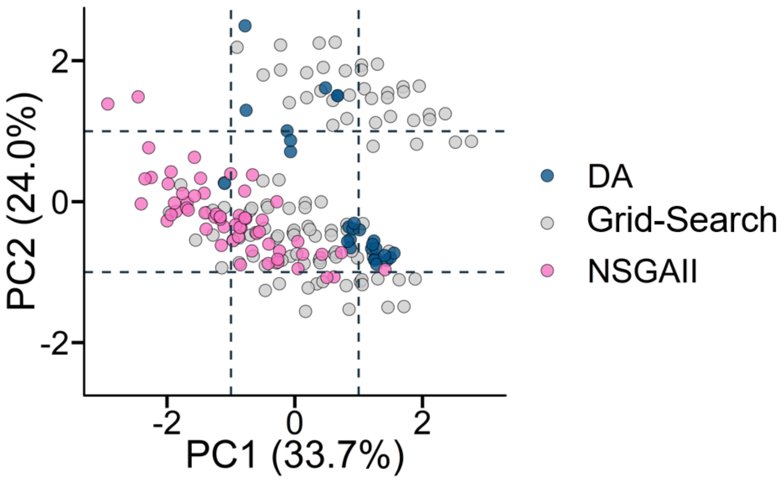


**Supplementary Fig. 22 | PCA visualization of initial grid-search, single-objective (DA)- and dual-objective (NSGAII)-optimized compositions using FALCON-driven optimization of anionic LNPs.**

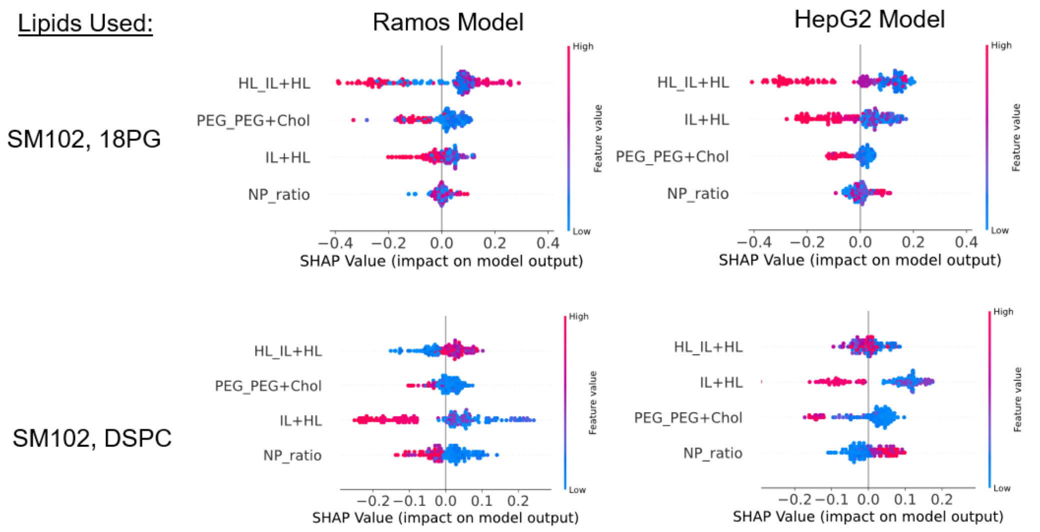


**Supplementary Fig. 23 | SHAP summary plots of models trained on anionic (18PG) or zwitterionic (DSPC) SM102 FALCON datasets for Ramos B and HepG2 transfection.** SHAP summary plots display the effect of feature values on final model predictions. Each row is defined by a specific feature, with each dot representing a unique LNP. The color displays the corresponding feature value used in the LNP, and the point's scatter on the x-axis displays the impact on the model output for that LNP, where further to the right is a positive impact (e.g., higher predicted transfection) and to the left is a negative impact (e.g., lower predicted transfection).

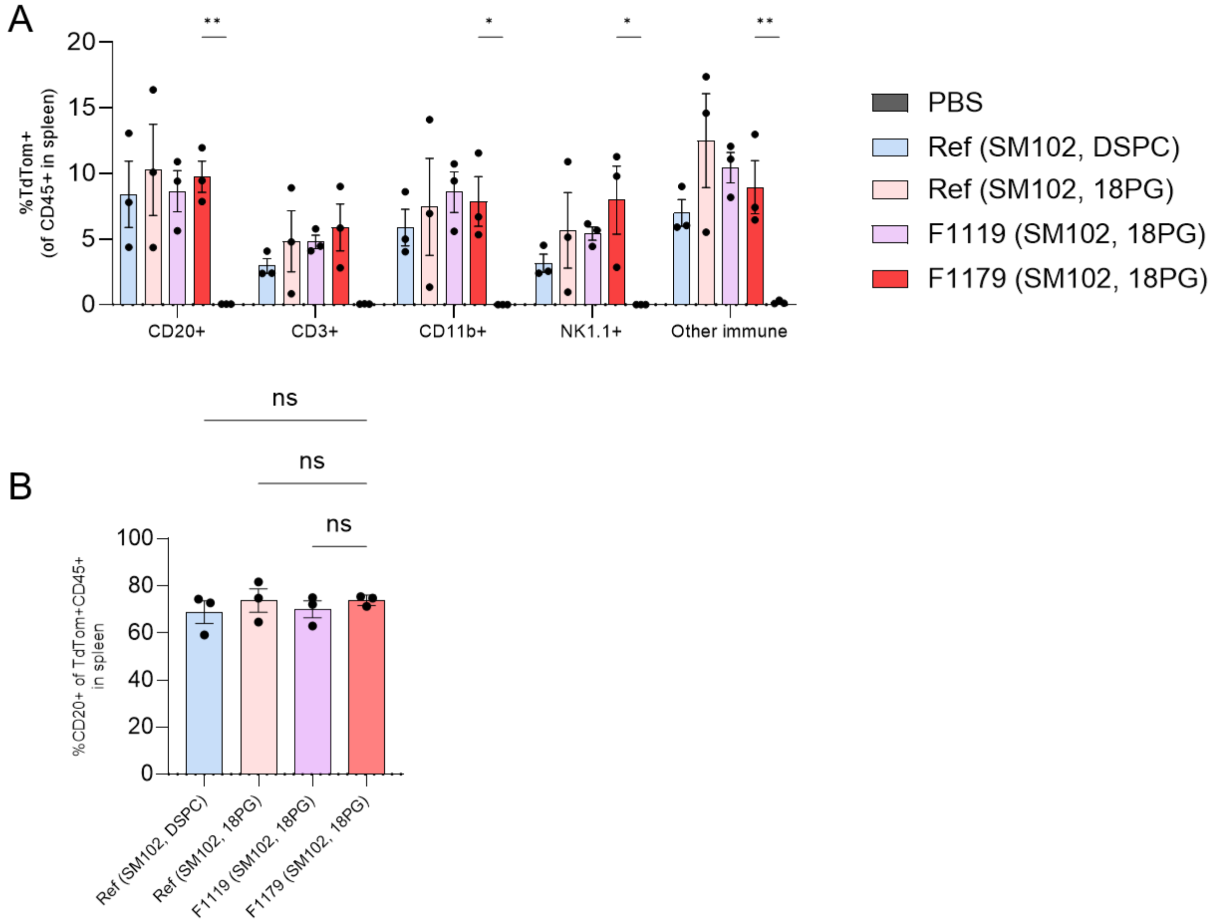


**Supplementary Fig. 24 | *In vivo* Ai9 model transgene expression in spleen immune compartment and B cell selectivity.** CD20+ B cells, CD3+ T cells, NK1.1+ NK cells, CD11b+ myeloid cells, and other (CD45+CD20-CD3-NK1.1-CD11b-) immune cells. TdTom+ cells indicate successful gene recombination.

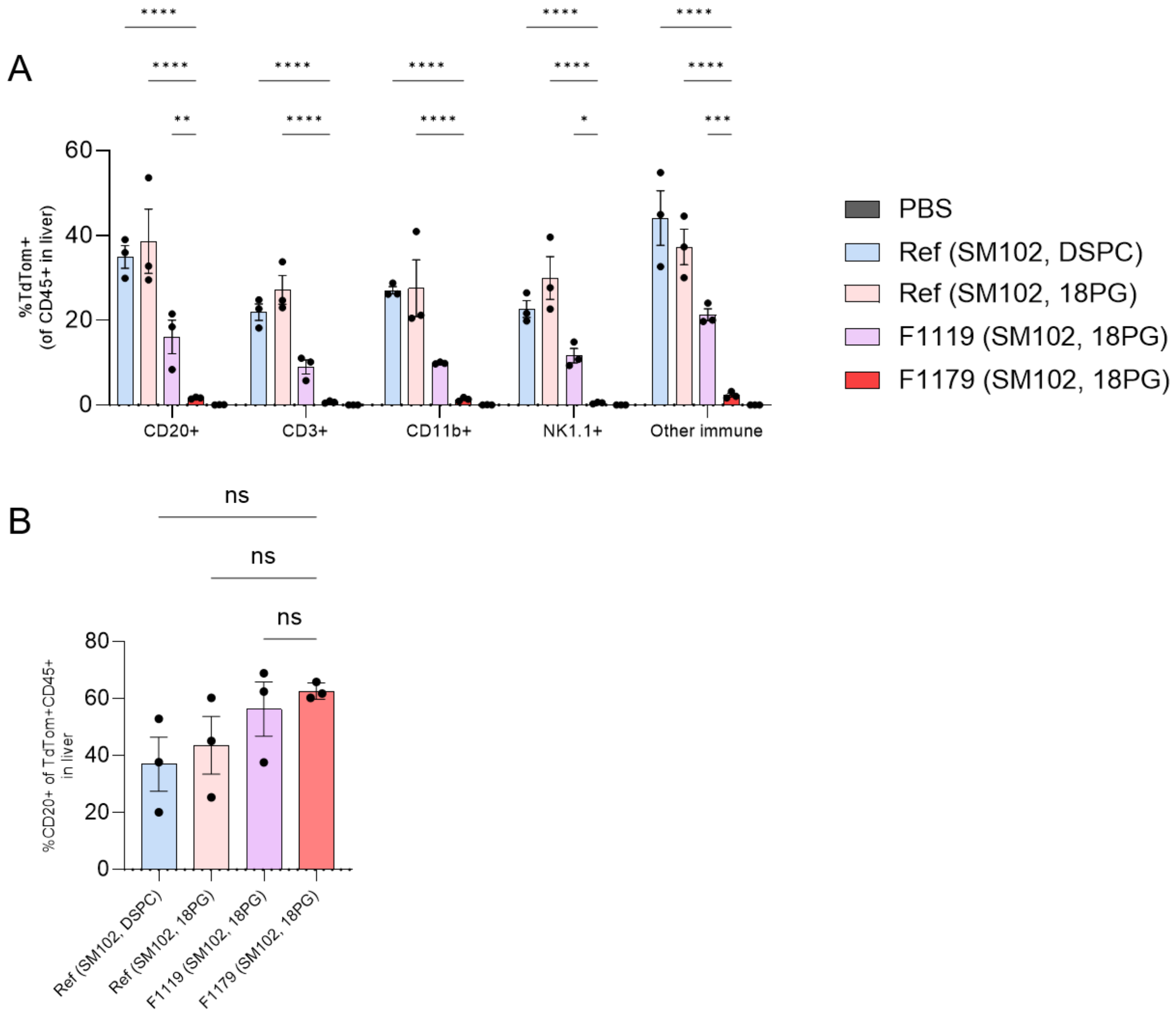


**Supplementary Fig. 25 | *In vivo* Ai9 model transgene expression in liver immune compartment and B cell selectivity.** CD20+ B cells, CD3+ T cells, NK1.1+ NK cells, CD11b+ myeloid cells, and other (CD45+CD20-CD3-NK1.1-CD11b-) immune cells. TdTom+ cells indicate successful gene recombination.

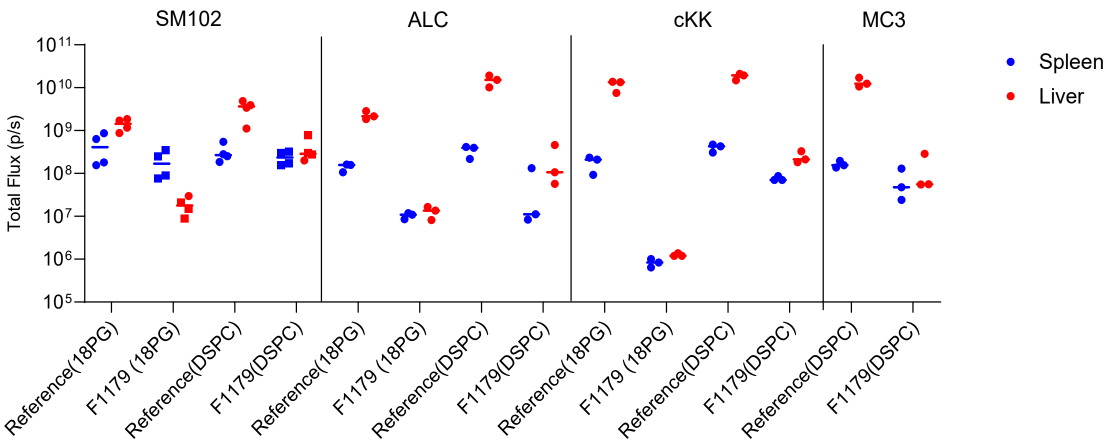


**Supplementary Fig. 26 | *Ex vivo* luminescence assay of spleen and liver after intravenous injection of F1179 and reference LNPs containing various IL and HL combinations.** Quantification of ex vivo luminescence flux in harvested organs 6 h after intravenous injection of mFluc encapsulated LNPs.

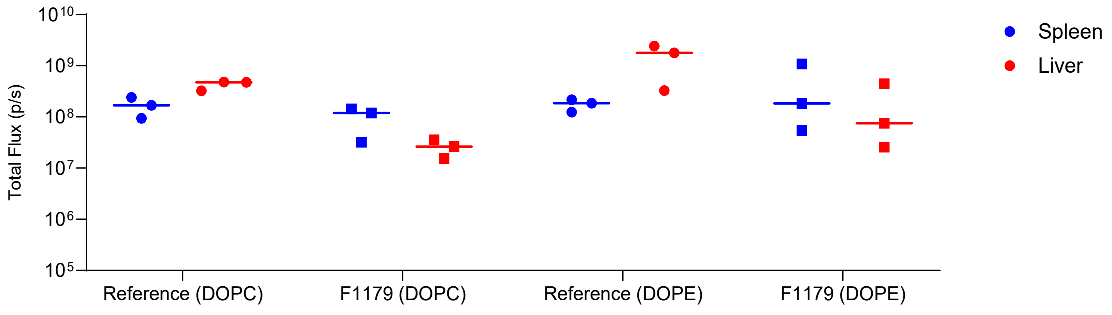


**Supplementary Fig. 27 | *Ex vivo* luminescence assay of spleen and liver after intravenous injection of SM102-based F1179 and reference LNPs containing different zwitterionic HLs.** Quantification of ex vivo luminescence flux in harvested organs 6 h after intravenous injection of mFluc encapsulated LNPs.

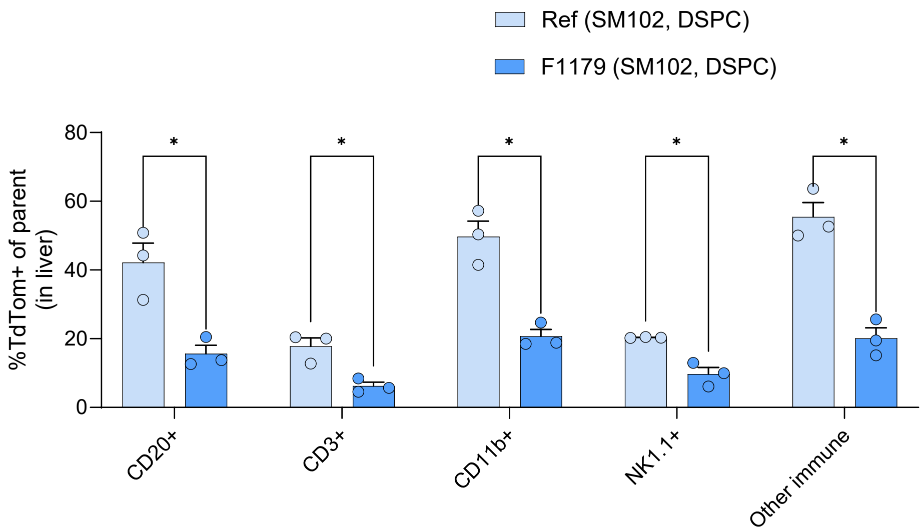


**Supplementary Fig. 28 | *In vivo* Ai9 model transgene expression in liver immune compartment cells comparing reference and F1179 LNPs with zwitterionic DSPC helper lipid.** mRNA Cre-recombinase was delivered by LNPs injected intravenously into Ai9 mice. TdTom+ cells indicate successful gene recombination.

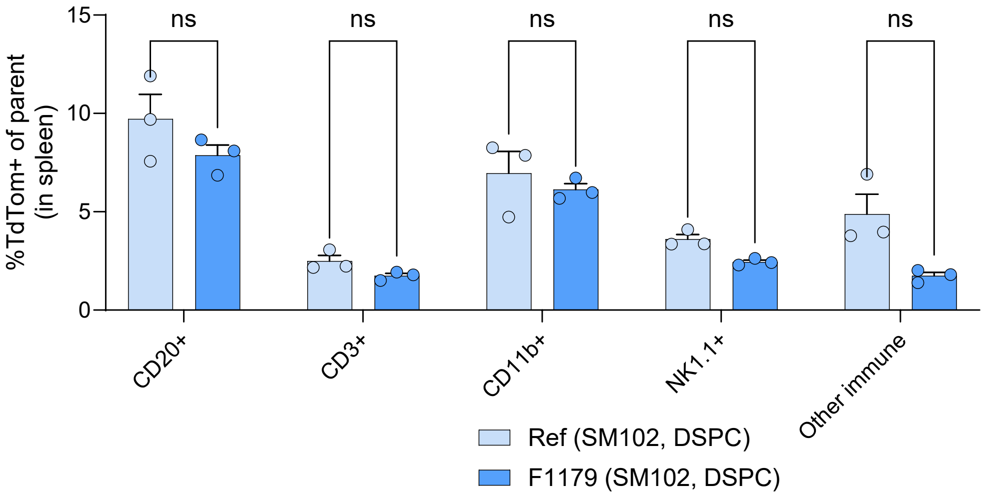


**Supplementary Fig. 29 | *In vivo* Ai9 model transgene expression in spleen immune compartment cells comparing reference and F1179 LNPs with zwitterionic DSPC helper lipid.** mRNA Cre-recombinase was delivered by LNPs injected intravenously into Ai9 mice. TdTom+ cells indicate successful gene recombination.

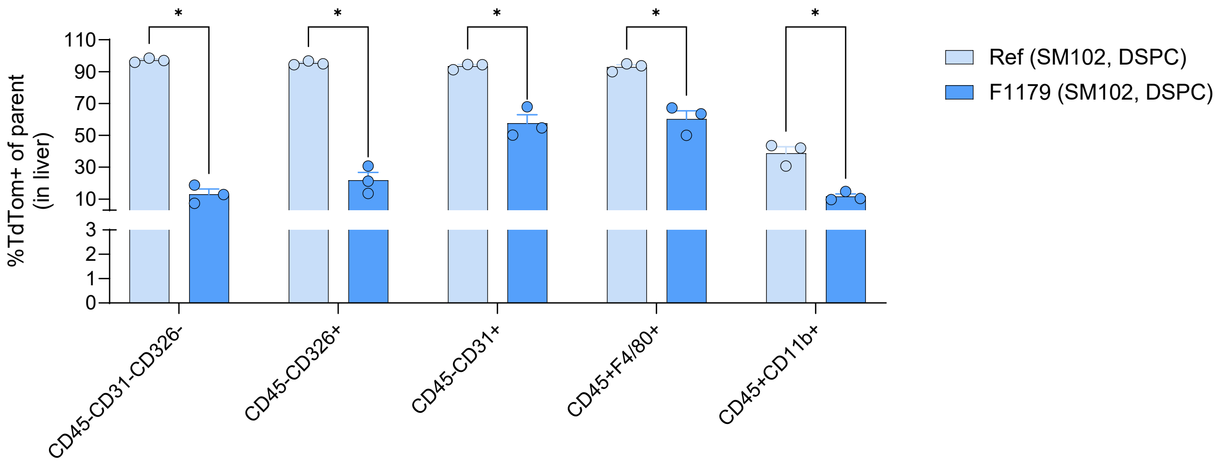


**Supplementary Fig. 30 | *In vivo* Ai9 model transgene expression in liver non-immune and myeloid lineage cells comparing reference and F1179 LNPs with zwitterionic DSPC helper lipid.** mRNA Cre-recombinase was delivered by LNPs injected intravenously into Ai9 mice. TdTom+ cells indicate successful gene recombination.

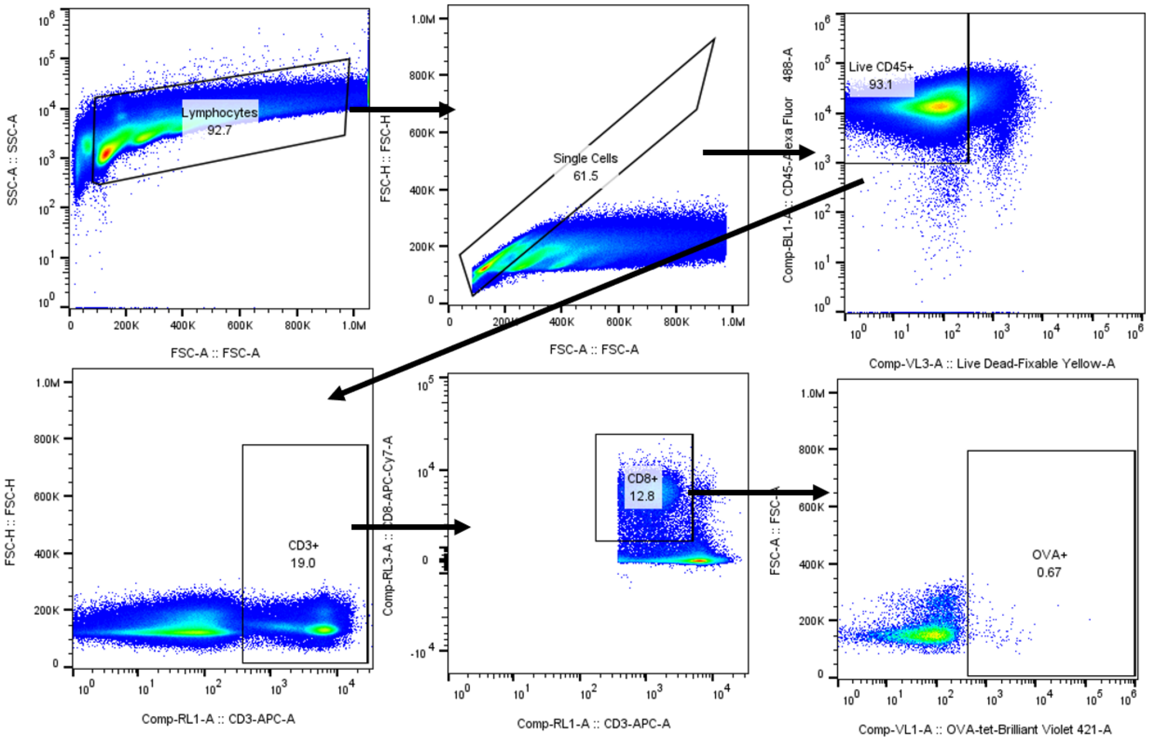


**Supplementary Fig. 31 | Gating strategy for *in vivo* assessment of splenic OVA-specific CD8+ T cells.** Representative flow plots show sequential gating: cells by size (FSC-A) and granularity (SSC-A), singlets based on FSC-H/FSC-A parameters, and live CD45+ cells using fixable live/dead exclusion. CD3+CD8+ for CD8+ T cells and OVA-tetramer positive for antigen-specific T cells. The OVA-tetramer+ representative gating on PBS-treated mice is shown.

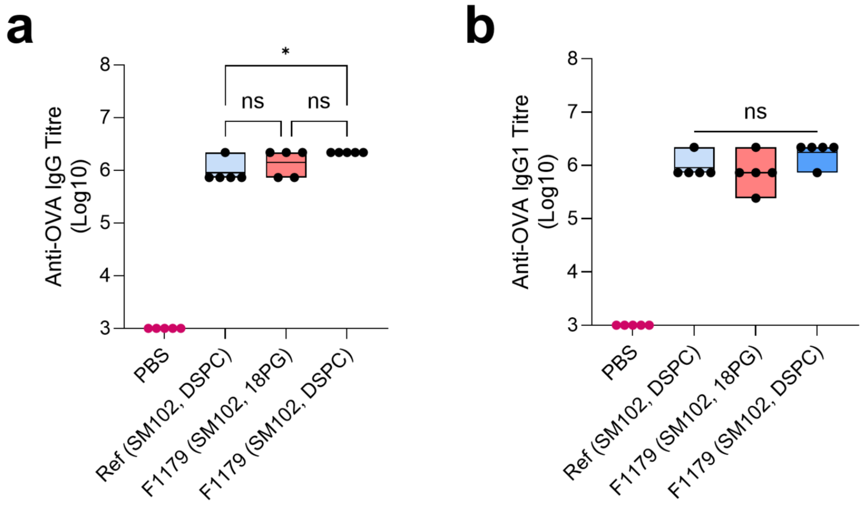


**Supplementary Fig. 32 | Serum anti-OVA IgG titer collected from mice after vaccination with reference and F1179 LNPs. a.** Total anti-OVA IgG endpoint titers. **b.** anti-OVA IgG1 endpoint titers. Endpoint titers were calculated from ELISA measurements using serial dilutions.

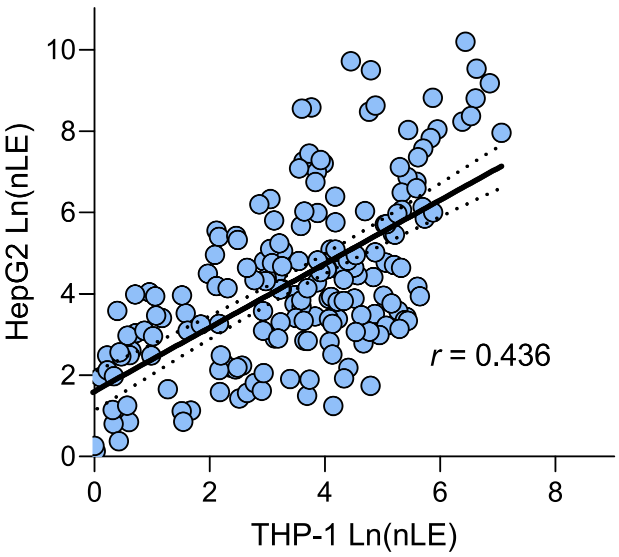


**Supplementary Fig. 33 | Transfection correlation plot between THP-1 and HepG2 cells.** Moderate positive correlation ($r$ = 0.436) highlights the complexity of isolating monocyte cell-specific formulations and the need for multi-objective search algorithms.

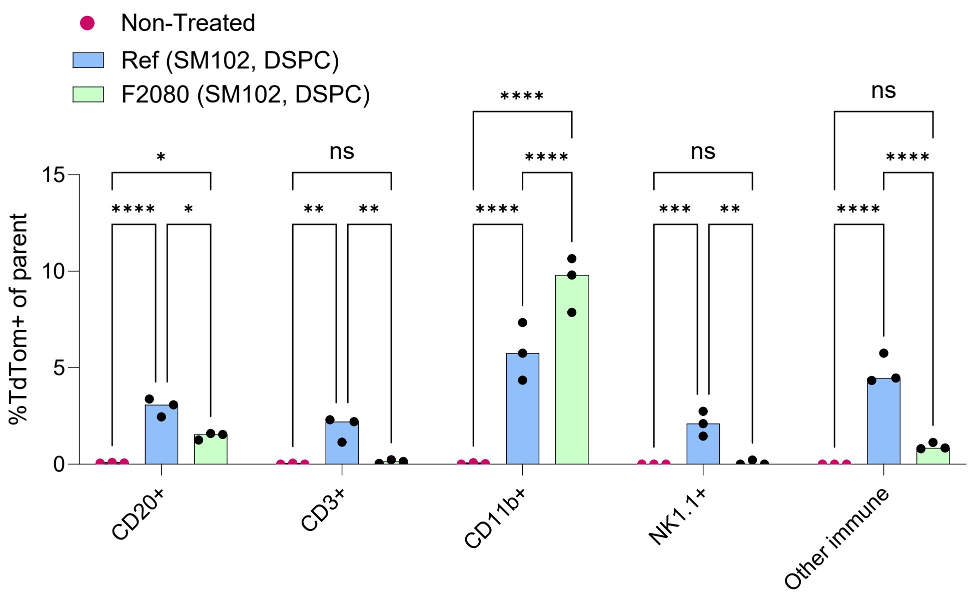


**Supplementary Fig. 34 | *In vivo* Ai9 model transgene expression in liver immune-cell populations comparing F2080 with the reference LNPs.** mRNA Cre-recombinase was delivered by LNPs injected intravenously into Ai9 mice. TdTom+ cells indicate successful transfection and gene recombination.

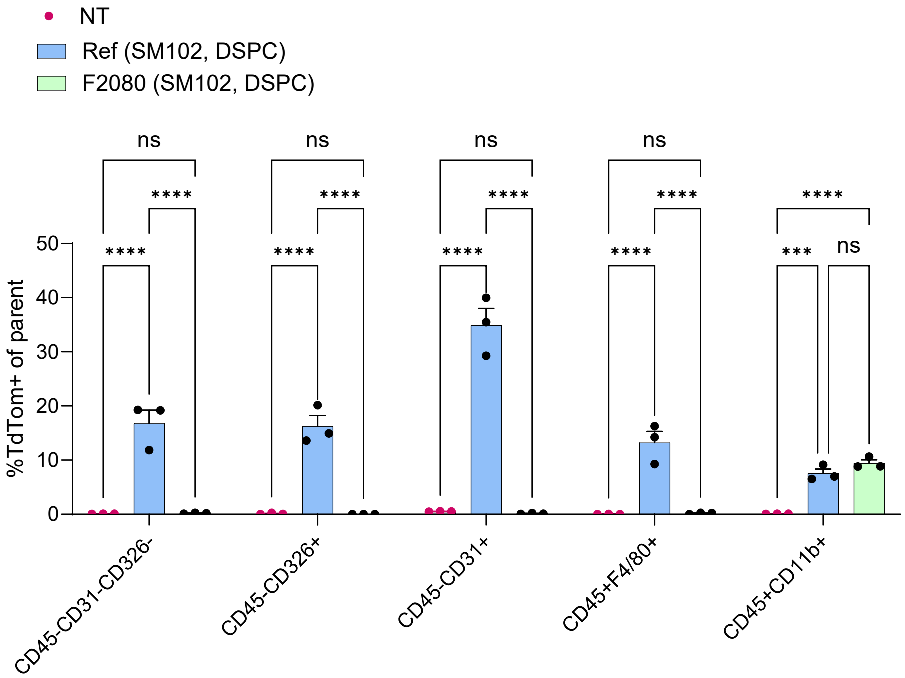


**Supplementary Fig. 35 | *In vivo* Ai9 model transgene expression in liver non-immune and myeloid cell populations comparing F2080 with the reference LNPs.** mRNA Cre-recombinase was delivered by LNPs injected intravenously into Ai9 mice. TdTom+ cells indicate successful transfection and gene recombination.

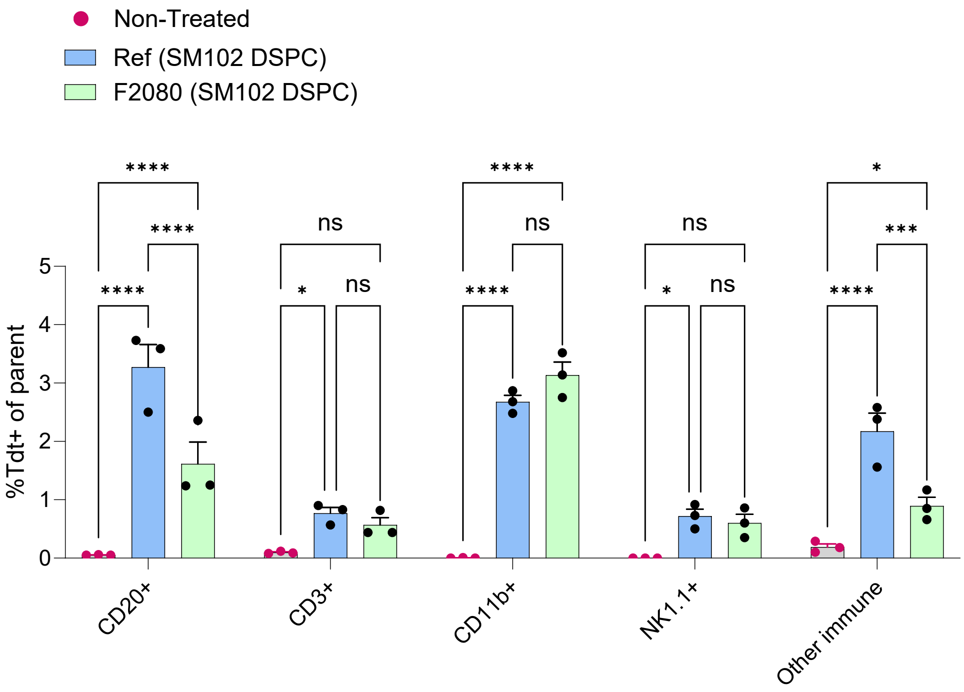


**Supplementary Fig. 36 | *In vivo* Ai9 model transgene expression in spleen immune cell populations comparing F2080 with the reference LNPs.** mRNA Cre-recombinase was delivered by LNPs injected intravenously into Ai9 mice. TdTom+ cells indicate successful transfection and gene recombination.

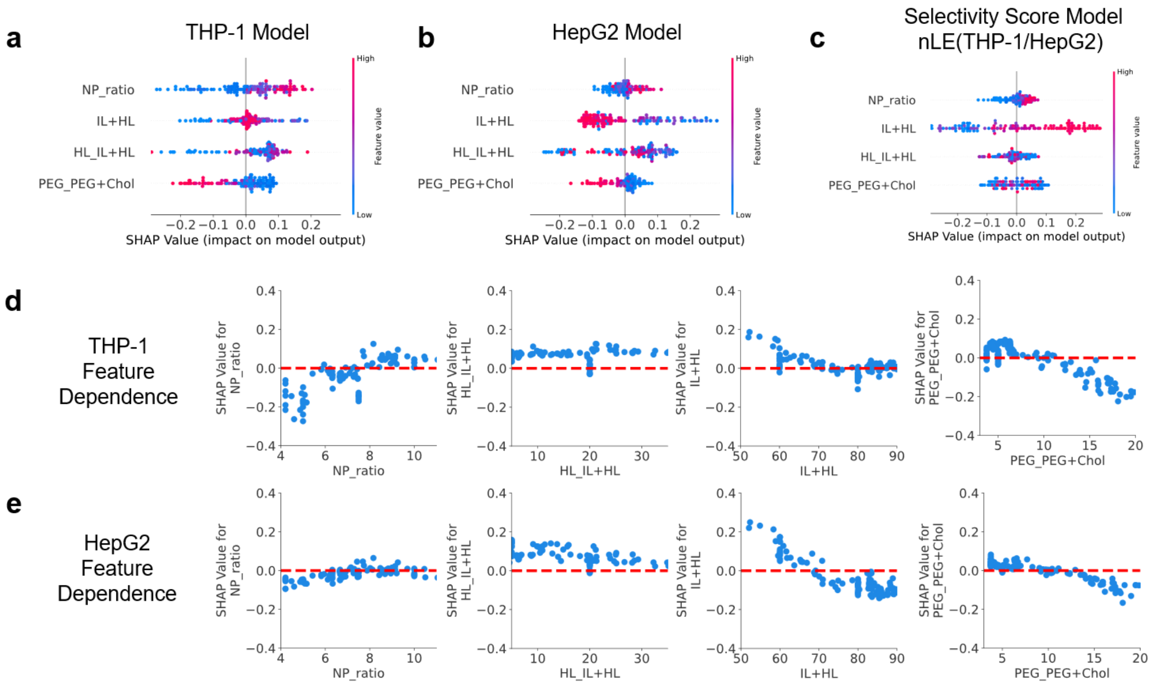


**Supplementary Fig. 37 | SHAP analysis of models trained on FALCON datasets for THP-1 and HepG2 transfection with SM102 and DSPC-based LNPs. a-c.** Summary SHAP beeswarm plots illustrating the overall impact of feature values on predicted THP-1 cell transfection (**a**), HepG2 cell transfection (**b**), and the selectivity score, calculated as the fold change in the predicted nLE of THP-1 over HepG2 (**c**). **d-e.** SHAP dependence plots showing the correlation between each compositional parameter and its corresponding SHAP value in THP-1 (**d**) and HepG2 (**e**) models.

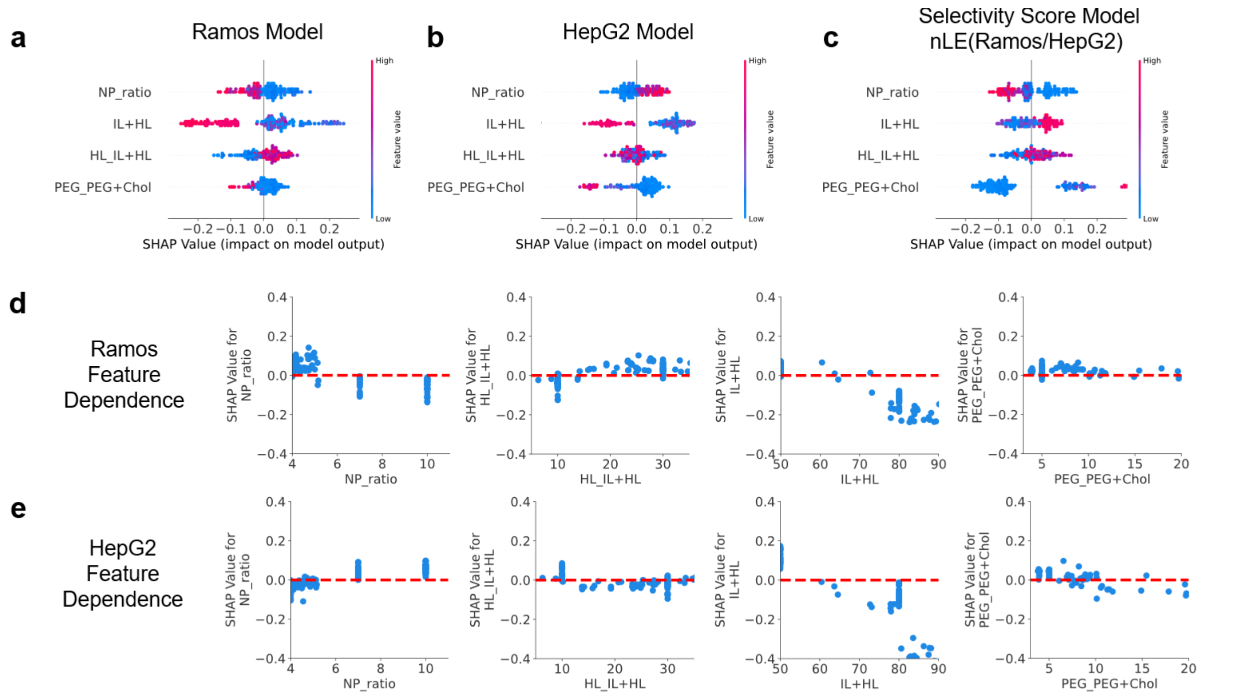


**Supplementary Fig. 38 | SHAP analysis of models trained on FALCON datasets for Ramos and HepG2 transfection with SM102 and DSPC-based LNPs. a-c.** Summary SHAP beeswarm plots illustrating the overall impact of feature values on predicted Ramos B cell transfection (**a**), HepG2 cell transfection (**b**), and the selectivity score, calculated as the fold change in the predicted nLE of Ramos over HepG2 (**c**). **d, e.** SHAP dependence plots showing the correlation between each compositional parameter and its corresponding SHAP value in Ramos (**d**) and HepG2 (**e**) models.

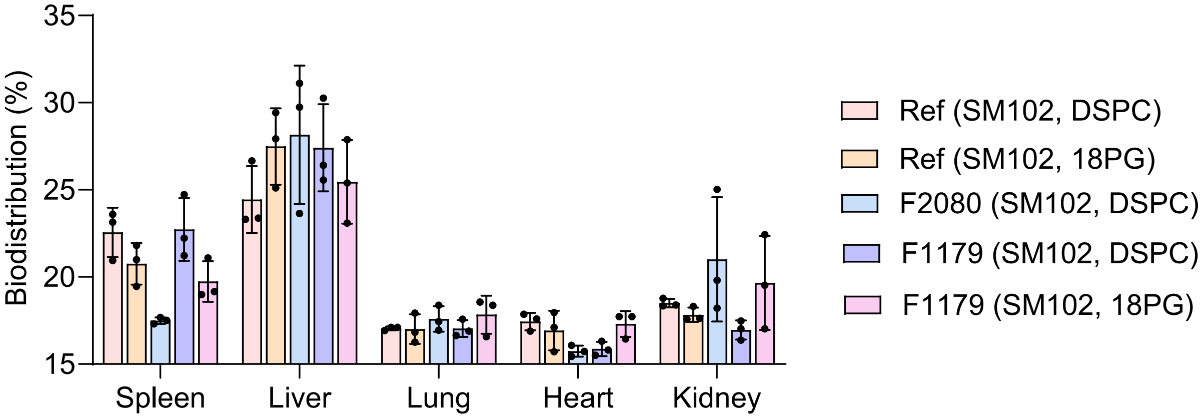


**Supplementary Fig. 39 | Relative *In Vivo* biodistribution of Top FALCON and Reference LNPs.** Mice were administered with 5 µg of Cy5-labeled mRNA LNPs via intravenous injection, and major organs were harvested and imaged after 6 h. Biodistribution is presented as the percentage of total fluorescence signal recovered from each organ.

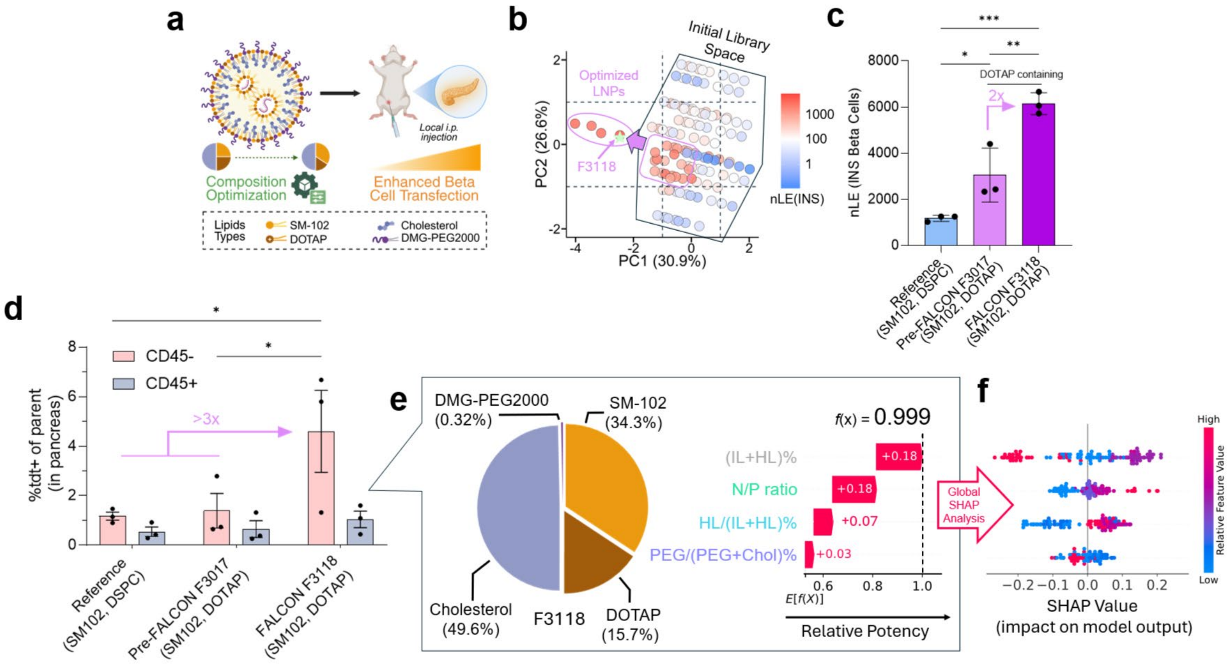


**Supplementary Fig. 40 | FALCON-guided single-objective compositional optimization of LNPs for delivery to pancreatic β cells. a.** Schematic of optimization target and lipid types used for iterative compositional optimization in INS-1E rat β cell line. **b.** PCA visualization of all tested LNP compositions, colored by selectivity score. The search spaces encompassed by the initial grid-search library and by FALCON optimized LNPs are labelled. Arrow illustrates the progression of FALCON optimization beyond the initial library space. **c.** Side-by-side comparison of the *in vitro* nLE of FALCON-identified F3118 against the commercial Moderna reference, and the top Pre-FALCON (F3017) LNP controls **d.** Single-cell transfection of cells in the pancreas, gated with CD45 pan-immune marker. **e.** Percent molar composition of F3118 and the predicted contribution of each of its individual feature values towards its relative potency (waterfall plot). **f.** Summary SHAP beeswarm plots illustrating the overall impact of feature values (global feature importance) on predicted β cell transfection. The nLE (normalized luciferase expression) refers to raw readings from *in vitro* luciferase transfection experiments, which were blank-subtracted and batch normalized against a set of internal controls. The nLE values from panel **b** are displayed on a logarithmic scale to allow visualization of the full dataset distribution. Data are presented as mean ± SEM from a representative experiment (n = 3). The *p*-values were determined via one-way ANOVA with Tukey's multiple comparisons test. ns: $p > 0.05$, *$p < 0.05$, **$p < 0.01$. Panel **a** was created with BioRender and released under a Creative Commons Attribution-NonCommercial-NoDerivs 4.0 International license.

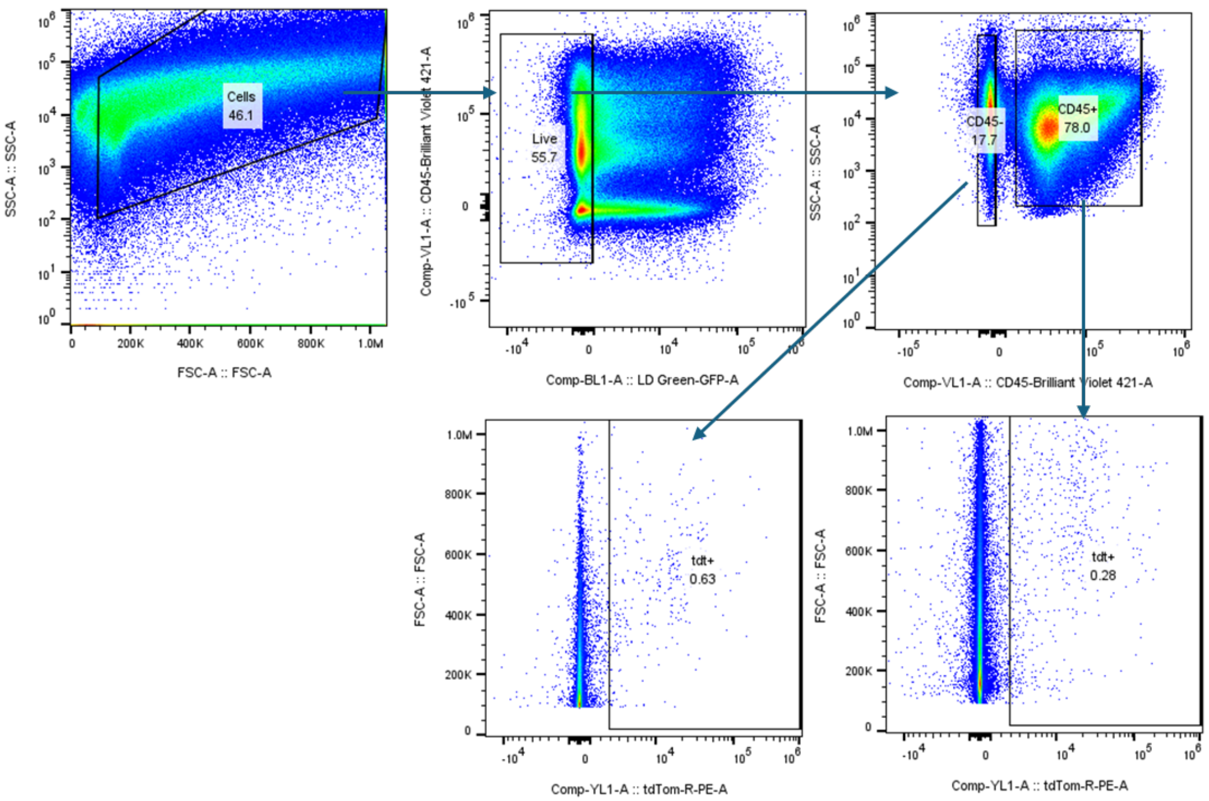


**Supplementary Fig. 41 | Gating strategy for *in vivo* assessment of pancreas cell transfection in Ai9 mice via flow cytometry after intraperitoneal administration of LNPs.** Representative flow plots show sequential gating: cells by size (FSC-A) and granularity (SSC-A), singlets based on FSC-H/FSC-A parameters, and live cells using fixable live/dead exclusion. CD45+ for pan-immune cells and CD45- for other cells (including β cells). The tdTom+ (Tdt+) value was used to assess successful transgene expression, and representative gating on PBS-treated mice is shown.

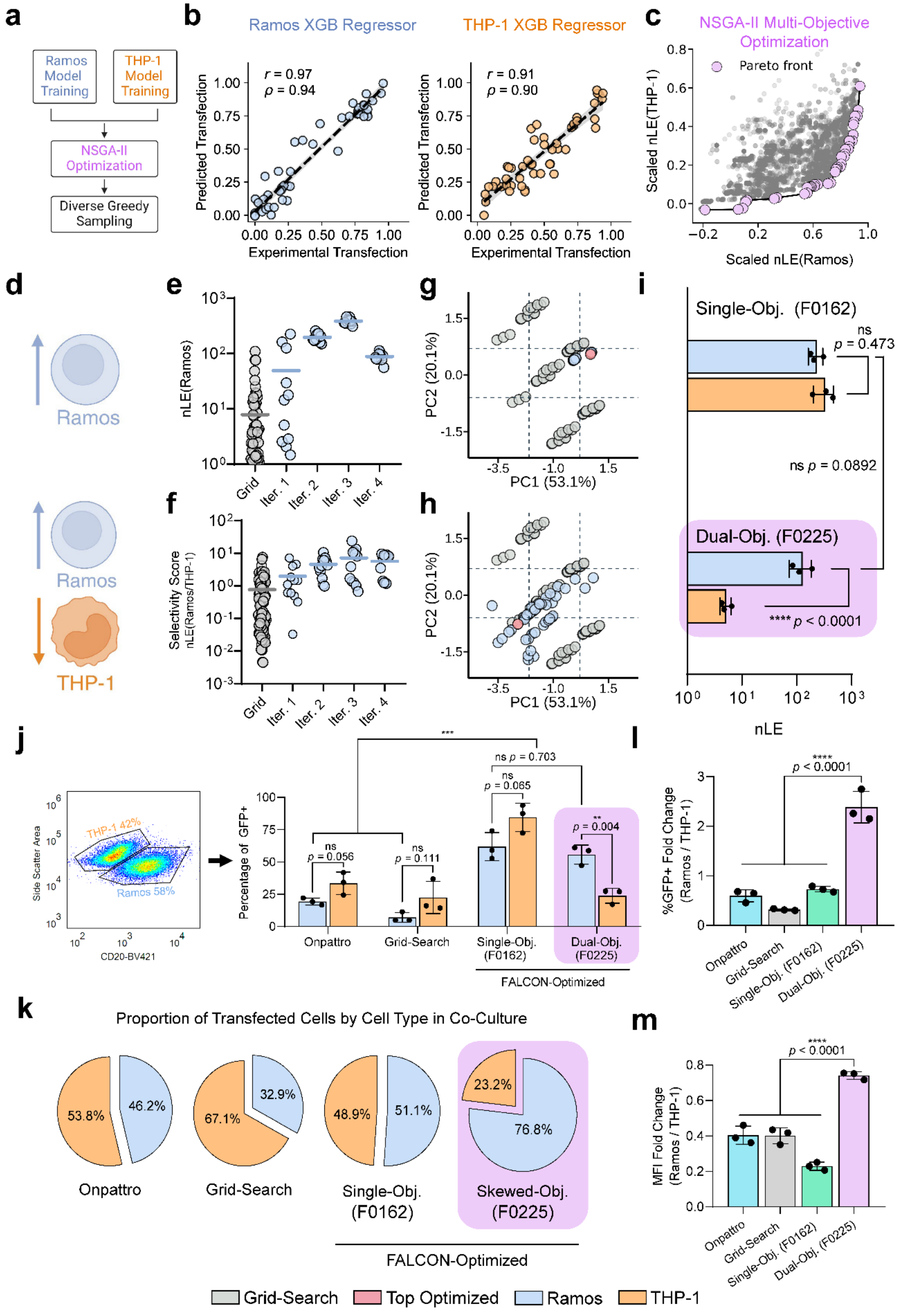
a
Ramos Model Training
THP-1 Model Training
NSGA-II Optimization
Diverse Greedy Sampling
b
Ramos XGB Regressor
THP-1 XGB Regressor
r = 0.97
ρ = 0.94
r = 0.91
ρ = 0.90
Predicted Transfection
Experimental Transfection
c
NSGA-II Multi-Objective Optimization
Pareto front
Scaled nLE(THP-1)
Scaled nLE(Ramos)
d
Ramos
Ramos
THP-1
e
nLE(Ramos)
Grid
Iter. 1
Iter. 2
Iter. 3
Iter. 4
f
Selectivity Score
nLE(Ramos/THP-1)
g
PC2 (20.1%)
PC1 (53.1%)
h
PC2 (20.1%)
PC1 (53.1%)
i
Single-Obj. (F0162)
ns p = 0.473
ns p = 0.0892
Dual-Obj. (F0225)
**** p < 0.0001
nLE
j
THP-1 42%
Ramos 58%
Side Scatter Area
CD20-BV421
Percentage of GFP+
***
ns p = 0.703
ns p = 0.065
ns p = 0.056
ns p = 0.111
** p = 0.004
Onpattro
Grid-Search
Single-Obj. (F0162)
Dual-Obj. (F0225)
FALCON-Optimized
k
Proportion of Transfected Cells by Cell Type in Co-Culture
53.8%
46.2%
67.1%
32.9%
48.9%
51.1%
23.2%
76.8%
Onpattro
Grid-Search
Single-Obj. (F0162)
Skewed-Obj. (F0225)
FALCON-Optimized
l
%GFP+ Fold Change (Ramos / THP-1)
**** p < 0.0001
Onpattro
Grid-Search
Single-Obj. (F0162)
Dual-Obj. (F0225)
m
MFI Fold Change (Ramos / THP-1)
**** p < 0.0001
Onpattro
Grid-Search
Single-Obj. (F0162)
Dual-Obj. (F0225)
Grid-Search
Top Optimized
Ramos
THP-1

**Supplementary Fig. 42 | Dual-objective FALCON-driven compositional optimization of LNPs for selective B cell and monocyte transfection. a.** Flowchart depicting multi-objective computational workflow. **b.** Representative hold-out performance of both Ramos B and THP-1 transfection models. **c.** Representative Pareto front illustrating optimal trade-off solutions for skewed Ramos B cell transfection identified via the NSGA-II algorithm. **d.** Parallel FALCON-driven LNP optimization configured for Ramos B cell transfection only (Row 1) and selective Ramos B cell transfection against THP-1 cell transfection (Row 2). **e**, **f.** Dot plot of normalized LE values achieved by LNPs tested in the initial grid-search library and four subsequent FALCON-guided iterative cycles. **g, h.** PCA plots of FALCON LNPs showing regions of convergence in the compositional search space for each optimization task. **i.** Comparison of normalized LE and Selectivity (Ramos/THP-1) in Ramos B cells and THP-1 cells achieved by the top FALCON-optimized LNPs from each task. **j–m.** Cell-type selectivity of FALCON LNPs in co-culture. **j.** Representative gating showing the relative percentage of Ramos B and THP-1 cell populations present in co-culture, used to compute absolute transfection efficiency (GFP+ percentage) of LNPs in each cell type. **k.** The transfection profile for each LNP formulation is shown as a pie chart, illustrating the proportion of total transfected cells contributed by each cell type. **l, m.** Fold change in GFP+ percentage (**l**) and MFI (**m**) in Ramos B cells relative to THP-1 cells. The nLE (normalized luciferase expression) refers to raw readings from *in vitro* luciferase transfection experiments, which were blank-subtracted and batch normalized against a set of internal controls. Data are presented as mean ± SEM from a representative experiment (n = 3). The *p*-values were determined via one-way ANOVA with Tukey's multiple comparisons test, and multiple *t*-tests with Benjamini-Krieger-Yekutieli correction (FDR Q = 1%). ns: $p > 0.05$, **$p < 0.01$, ***$p < 0.001$, ****$p < 0.0001$. Panels **a** and **d** were created with BioRender and released under a Creative Commons Attribution-NonCommercial-NoDerivs 4.0 International license.

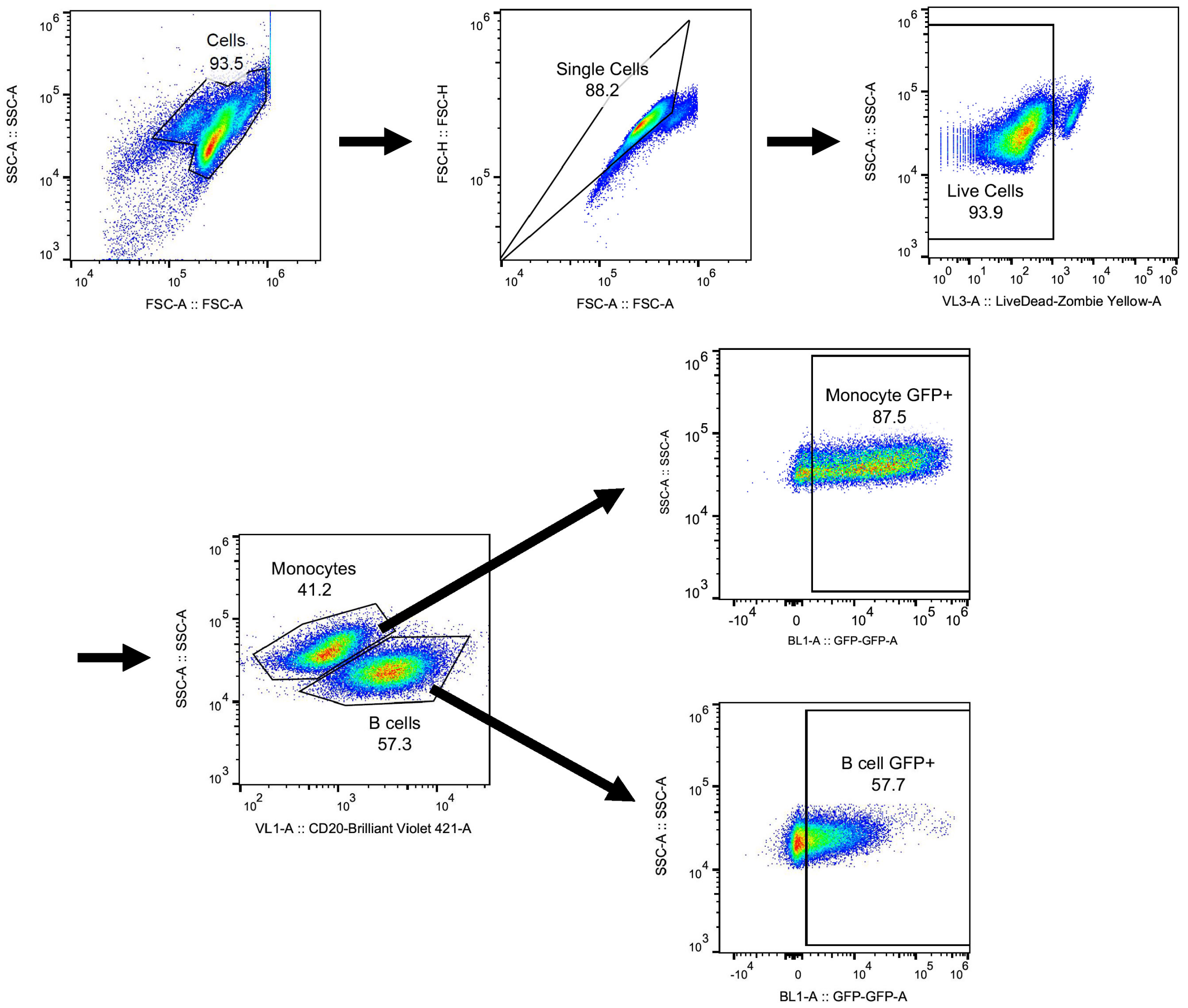


**Supplementary Fig. 43 | Gating strategy for co-culture assessment of LNP selectivity via flow cytometry.** Representative flow plots show sequential gating: cells by size (FSC-A) and granularity (SSC-A), singlets based on FSC-H/FSC-A parameters, and live cells using Zombie Yellow exclusion. Monocytes and B cells in co-culture were distinguished based on CD20 (BV421) expression. For each gated cell type, GFP expression was measured to assess transfection efficiency.

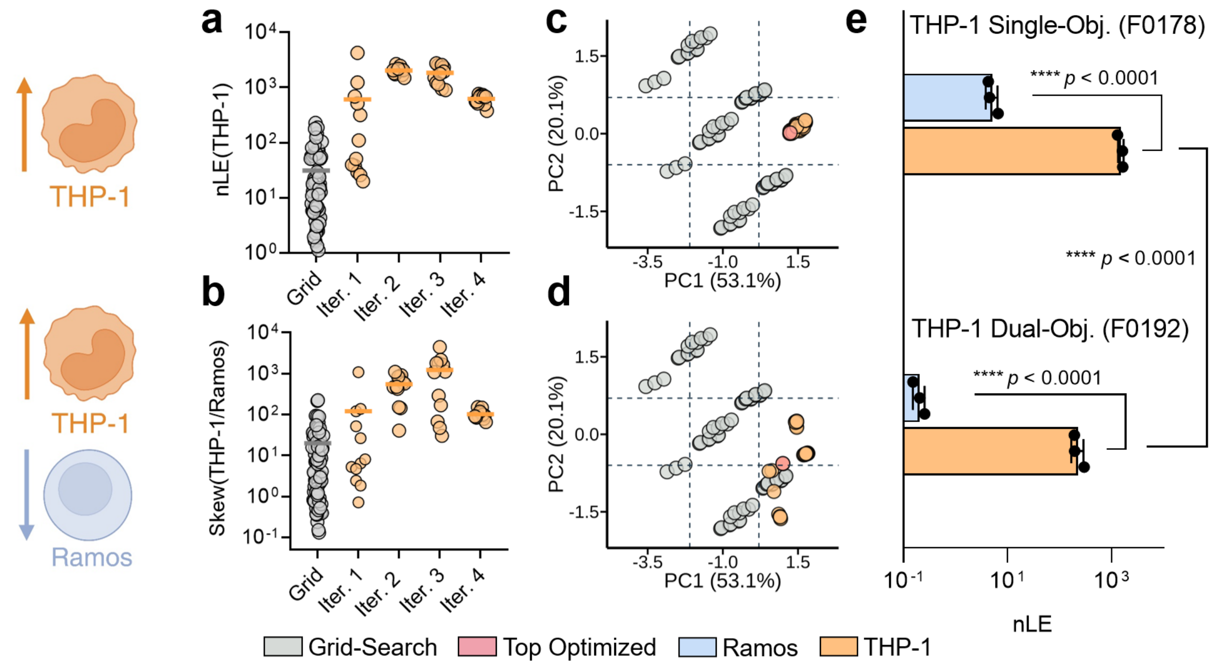


**Supplementary Fig. 44 | Dual-objective optimization of LNPs for selective monocyte transfection *in vitro*.** Parallel FALCON-driven LNP optimization configured for THP-1 cell transfection only (**a**) and selective THP-1 cell transfection against Ramos B cell transfection (**b**). **a, b.** Dot plot of normalized LE values achieved by LNPs tested in the initial grid-search library and four subsequent FALCON-guided iterative cycles. **c, d.** PCA plots of FALCON LNPs showing regions of convergence in the compositional search space for each optimization task. **e.** Comparison of normalized LE in Ramos B cells and THP-1 cells achieved by the top FALCON-optimized LNPs from each task. nLE (normalized luciferase expression) refers to raw readings from *in vitro* luciferase transfection experiments, which were blank-subtracted and batch normalized against a set of internal controls. Data are presented as mean ± SEM from a representative experiment (n = 3). The p-values were determined via one-way ANOVA with Tukey's multiple comparisons test, and multiple t-tests with Benjamini-Krieger-Yekutieli correction (FDR Q = 1%).

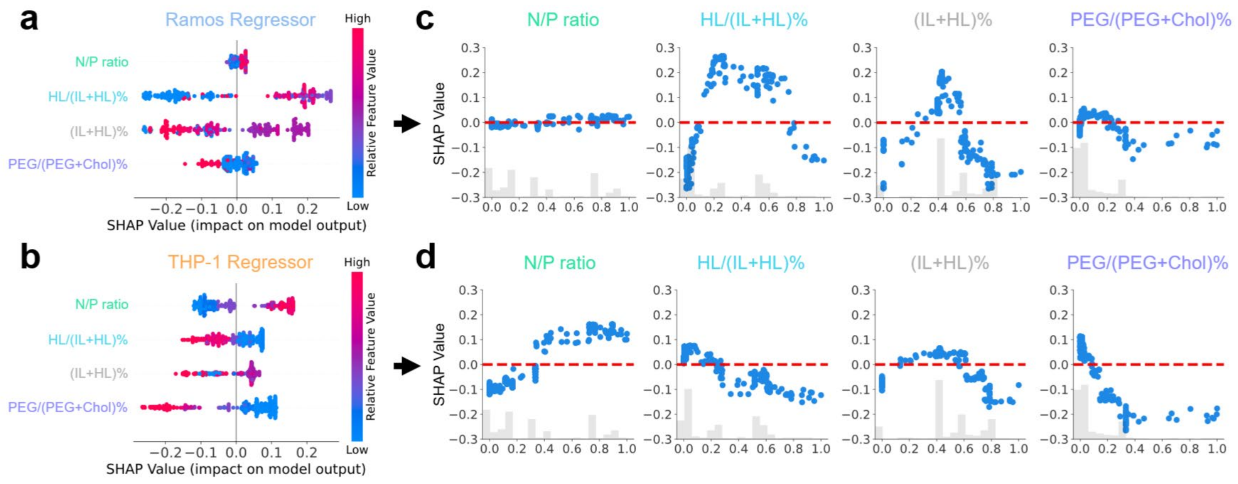


**Supplementary Fig. 45 | SHAP feature importance analysis of Ramos B and THP-1 FALCON models. a, b.** Summary SHAP beeswarm plots illustrating the overall impact of feature values on the final transfection predictions of trained Ramos B cell (**a**) and THP-1 cell (**b**) XGBoost models. Each point represents a single LNP formulation; color indicates relative feature value. **c, d.** SHAP dependence plots showing the correlation between each compositional parameter and its corresponding SHAP value in Ramos B (**c**) and THP-1 (**d**) models.

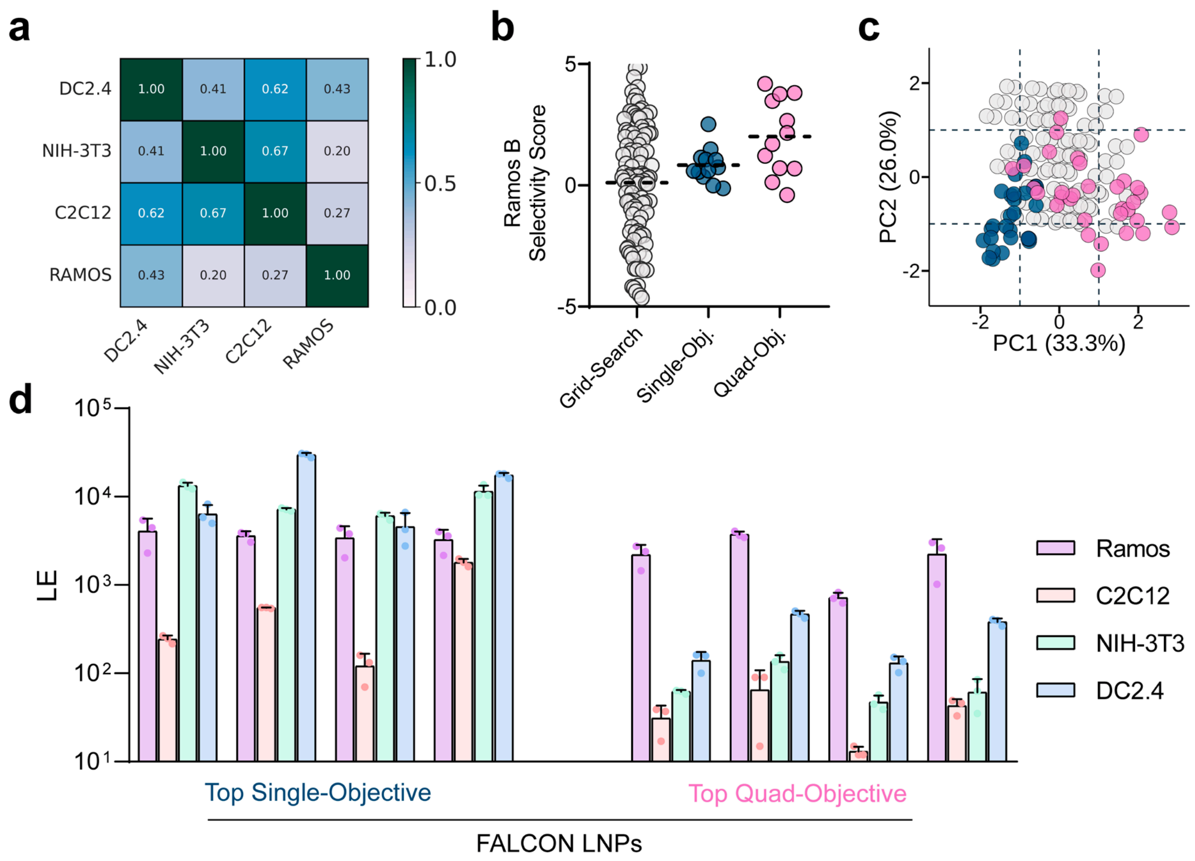


**Supplementary Fig. 46 | Quadruple-objective optimization of LNPs in Ramos, C2C12, DC2.4, and NIH-3T3 cells.** FALCON was deployed to optimize the composition of LNPs from a starting library consisting of SM-102 ionizable lipid, 18PG, cholesterol, and DMG-PEG2000 for high Ramos B cell transfection while minimizing transfection of C2C12, NIH-3T3, and DC2.4 cell lines. **a.** Heatmap of Pearson correlation coefficients (*r*) between LE measurements across four tested cell types (DC2.4, NIH-3T3, C2C12, and Ramos B cells). Strong positive correlations indicate shared trends in LNP-mediated transfection, whereas low correlations reflect contrasting transfection responses across cell types. **b.** Selectivity of formulations tested through grid-search, as well as FALCON-driven single-objective and quad-objective optimization. Selectivity score was calculated as LE_Ramos/1/3×LE_(C2C12+NIH-3T3-DC2.4), where LE refers to raw luciferase luminescence readings. Dotted lines indicate the average selectivity score of each group. Quadruple-objective optimized FALCON LNPs demonstrated greater selectivity on average than grid-search or single-objective optimized LNPs. **c.** The PCA plot depicts the compositional space explored by each search algorithm. **d.** The LE values of the top four FALCON formulations from each optimization task were tested in the four cell types of interest. Quadruple-objective optimized FALCON LNPs maintained a high Ramos transfection while demonstrating decreased transfection in all other off-target cell types.

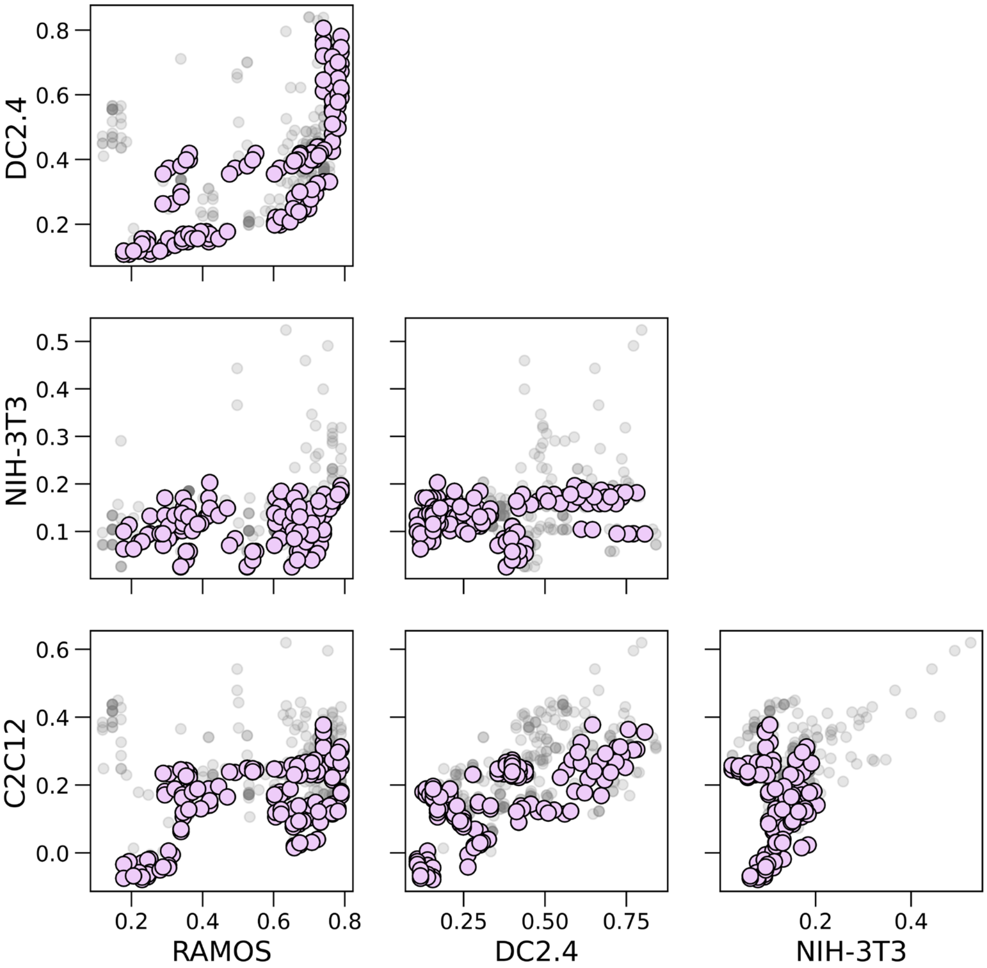


**Supplementary Fig. 47 | Representative pairwise Pareto plots illustrating the FALCON quadruple-objective optimization process.** NSGA-II was configured to identify LNPs with high Ramos B cell transfection, while minimizing transfection of NIH-3T3, C2C12, and DC2.4 cells. Each subplot shows a 2D projection of the multi-objective search space, where the Pareto front (pink points) represents LNPs that achieve the best possible tradeoffs. Selectivity is the most challenging to achieve against DC 2.4 cells, as they display a strong positive correlation with Ramos B cell transfection.

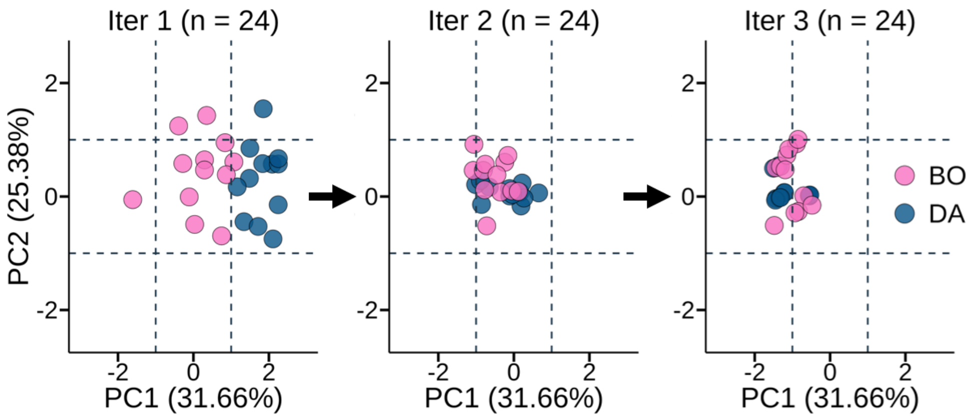


**Supplementary Fig. 48 | Comparison of dual annealing and Bayesian optimization search algorithm performance.** A preliminary iterative optimization study was conducted using both Bayesian Optimization (pink) and Dual Annealing (blue) to suggest 12 formulations each to test during each iteration. By the third iteration, both algorithms converged to focus on the same concentrated area.
